# Long COVID manifests with T cell dysregulation, inflammation, and an uncoordinated adaptive immune response to SARS-CoV-2

**DOI:** 10.1101/2023.02.09.527892

**Authors:** Kailin Yin, Michael J. Peluso, Xiaoyu Luo, Reuben Thomas, Min-Gyoung Shin, Jason Neidleman, Alicer Andrew, Kyrlia Young, Tongcui Ma, Rebecca Hoh, Khamal Anglin, Beatrice Huang, Urania Argueta, Monica Lopez, Daisy Valdivieso, Kofi Asare, Tyler-Marie Deveau, Sadie E. Munter, Rania Ibrahim, Ludger Ständker, Scott Lu, Sarah A. Goldberg, Sulggi A. Lee, Kara L. Lynch, J. Daniel Kelly, Jeffrey N. Martin, Jan Münch, Steven G. Deeks, Timothy J. Henrich, Nadia R. Roan

**Affiliations:** Gladstone Institutes, University of California, San Francisco, USA; Department of Urology, University of California, San Francisco, USA; Division of HIV, Infectious Diseases, and Global Medicine, University of California, San Francisco, USA; Core Facility Functional Peptidomics, Ulm University Medical Center, Meyerhofstrasse 1, Ulm, Germany; Department of Epidemiology and Biostatistics, University of California, San Francisco, USA; Zuckerberg San Francisco General Hospital and the University of California, San Francisco, USA; Division of Laboratory Medicine, University of California, San Francisco, USA; Division of Experimental Medicine, University of California, San Francisco, USA

**Author notes:** Equal Contribution (Kailin Yin, Michael J. Peluso).

## Abstract

Long COVID (LC), a type of post-acute sequelae of SARS-CoV-2 infection (PASC), occurs after at least 10% of SARS-CoV-2 infections, yet its etiology remains poorly understood. Here, we used multiple “omics” assays (CyTOF, RNAseq/scRNAseq, Olink) and serology to deeply characterize both global and SARS-CoV-2-specific immunity from blood of individuals with clear LC and non-LC clinical trajectories, 8 months following infection and prior to receipt of any SARS-CoV-2 vaccine. Our analysis focused on deep phenotyping of T cells, which play important roles in immunity against SARS-CoV-2 yet may also contribute to COVID-19 pathogenesis. Our findings demonstrate that individuals with LC exhibit systemic inflammation and immune dysregulation. This is evidenced by global differences in T cell subset distribution in ways that imply ongoing immune responses, as well as by sex-specific perturbations in cytolytic subsets. Individuals with LC harbored increased frequencies of CD4+ T cells poised to migrate to inflamed tissues, and exhausted SARS-CoV-2-specific CD8+ T cells. They also harbored significantly higher levels of SARS-CoV-2 antibodies, and in contrast to non-LC individuals, exhibited a mis-coordination between their SARS-CoV-2-specific T and B cell responses. RNAseq/scRNAseq and Olink analyses similarly revealed immune dysregulatory mechanisms, along with non-immune associated perturbations, in individuals with LC. Collectively, our data suggest that proper crosstalk between the humoral and cellular arms of adaptive immunity has broken down in LC, and that this, perhaps in the context of persistent virus, leads to the immune dysregulation, inflammation, and clinical symptoms associated with this debilitating condition.

## Introduction

Intense efforts are underway to determine the pathophysiology of post-acute sequelae of SARS-CoV-2 infection (PASC), a set of conditions affecting at least 10% of individuals recovering from COVID-19 ^1, 2, 3^. PASC, which includes the unexplained, debilitating symptoms that characterize LC, remains a major public health challenge despite the availability of SARS-CoV-2 vaccination and treatment ^4, 5, 6, 7, 8^. Although the underlying cause or causes of LC are incompletely understood, multiple mechanisms including microvascular dysregulation ^9, 10^, autoimmune phenomena ^11, 12, 13^, and reactivation of latent human herpesviruses ^12, 14, 15^ have been proposed as contributors to inflammatory responses, particularly in tissues, which could in turn drive symptoms that individuals experience. In addition, persistence of SARS-CoV-2 antigen can occur months after infection ^16^, and has recently been demonstrated in a subset of immunocompetent individuals with LC ^17, 18, 19^. However, there are currently no accepted therapies for LC, in part due to limited insight into the underlying mechanisms of the condition to date ^2^.

To try to better understand the molecular underpinnings of LC, multiple “omics”-based approaches have recently been implemented on plasma specimens. Such studies have revealed individuals with LC to more often have elevated levels of inflammatory cytokines such as IFNβ and IL8, but low levels of cortisol ^12, 20, 21^. These results are consistent with the ongoing immunologic perturbations that have been consistently observed in individuals experiencing LC ^12, 21, 22, 23, 24, 25, 26^. Serological analyses have also found the presence of auto-antibodies during the acute or post-acute phases of infection to be associated with LC ^11, 12, 13^, although this has not been observed consistently ^20, 27, 28^. Indeed, recent proteome-wide autoantibody analysis by PhIP-Seq revealed a clear autoreactivity signature associated with prior COVID-19, but no unique autoreactivity signature comparing people with and without LC ^28^. Intriguingly, a subset of auto-antibodies against chemokines have even been reported recently to associate with protection against LC ^29^. Elevated levels of SARS-CoV-2 antibodies have also been associated with LC ^20^. “Omics” analyses of immune cells in the form single-cell transcriptomics on PBMCs have likewise been performed, resulting in the classification of LC into multiple endotypes, and uncovering the persistent elevation of select immune subsets – including myeloid and NK subsets – in some phenotypes of LC ^12^. This study, however, did not examine individuals whose symptoms persisted beyond three months, and did not examine LC resulting from initial mild-to-moderate (non-hospitalized) cases of COVID-19, which comprise the vast majority of those experiencing this condition.

T cells play an important role in SARS-CoV-2 immunity and pathogenesis, yet relatively little is known about their role in LC. A limited set of studies that have examined SARS-CoV-2-specific T cell responses have implicated these cells in LC, albeit with conflicting results. While some studies have found elevated SARS-CoV-2-specific T cell responses in LC as compared to non-LC individuals ^26, 30^, we have observed faster decay of subsets of SARS-CoV-2-specific CD8+ T cells in the context of LC ^31^. Apart from a transcriptomic/CITE-seq analysis of SARS-CoV-2-specific CD8+ T cells by MIRA, which identified unique features associated with LC two to three months after COVID-19 hospitalization ^12^, in-depth analyses of the phenotypic and functional features of SARS-CoV-2-specific T cells from individuals with LC are lacking. In particular, the profile of CD4+ T cells, key orchestrators of adaptive immunity, in individuals with LC is currently unknown.

We have previously used deep phenotypic characterization of SARS-CoV-2-specific T cells by CyTOF to identify differentiation states, effector functions, and/or homing properties of SARS-CoV-2-specific T cells associated with long-lived memory responses, fatal COVID-19, vaccination, and hybrid immunity ^32, 33, 34, 35^, and to characterize pulmonary T cell responses in a mouse model of severe COVID-19 ^36^. As these insights into COVID-19 immunity and pathogenesis were obtained using these next-generation T cell characterization assays in ways that would not have been captured using solely more traditional T cell assays, we reasoned that a similar in-depth analysis could identify T cells that protect or contribute to the symptoms of LC.

Therefore, in this study we deeply characterized T cell immunity during the post-acute phase of infection by CyTOF. We then combined these data with standard serological analyses, as well as additional “omics” techniques: RNAseq/scRNAseq, and high-dimensional plasma proteomics using the Olink Explore Proximity Extension Assay (PEA), the latter of which enables simultaneous quantitation of 384 analytes from plasma. We leveraged a cohort of LC and non-LC individuals with detailed longitudinal characterization and biospecimen collection prior to SARS-CoV-2 vaccination or reinfection, which could confound interpretation of SARS-CoV-2-specific T cell and antibody responses, to identify clues to the immunologic processes that might drive LC. By performing this holistic, integrative analysis on a well-matched set of LC and non-LC individuals with consistent phenotypes for 8 months after infection, we were able to identify unique immune features associated with LC that inform on the mechanistic underpinnings of this debilitating disease.

## Results

### LC and non-LC participants

To study the phenotypes and effector functions of immune cells from individuals experiencing Long COVID-19 symptoms, we analyzed blood samples from 27 LC and 16 non-LC individuals from the San Francisco-based Long-term Impact of Infection with Novel Coronavirus (LIINC) cohort (NCT04362150) ^37^. Specimens were collected 8 months following infection, but individuals had been followed since at least 4 months post-infection to characterize LC over time. Individuals with LC were defined as those that consistently met the case definition for LC (at least one COVID-19-attributed symptom that was new or worsened since the time of SARS-CoV-2 infection, and was at least somewhat bothersome) at both 4 and 8 months, while clinically matched non-LC individuals did not experience any lingering symptoms for the entire 8 months after SARS-CoV-2 infection. Importantly, at the time of specimen collection (in 2020-2021), none of the participants had yet received a SARS-CoV-2 vaccine, which would confound the SARS-CoV-2-specific serological and T cell analyses (Fig. S1A).

Overall, the enrolled individuals had a median age of 46 years (range 19 to 71) and 58.1% identified as White (Table 1, Table 2, Table S1). Individuals with LC were more likely to be female (63% vs 44%) (Fig. S1B) and included those previously hospitalized during the acute phase of COVID-19 (26% vs 13%) (Fig. S1C). The individuals with LC analyzed herein were all highly symptomatic, and consistently exhibited LC symptoms over an 8-month period (Fig. S1D). Although the overall population was relatively healthy with few pre-existing comorbidities, when present these tended to be more common in the LC group (Fig. S1E), consistent with the current understanding that certain comorbid conditions such as obesity, diabetes, and pre-existing lung disease are likely to be risk factors for LC. This is also consistent with our observation that in our cohort, individuals with LC had higher BMI than those without LC (Fig. S1F).

**Table 1:**
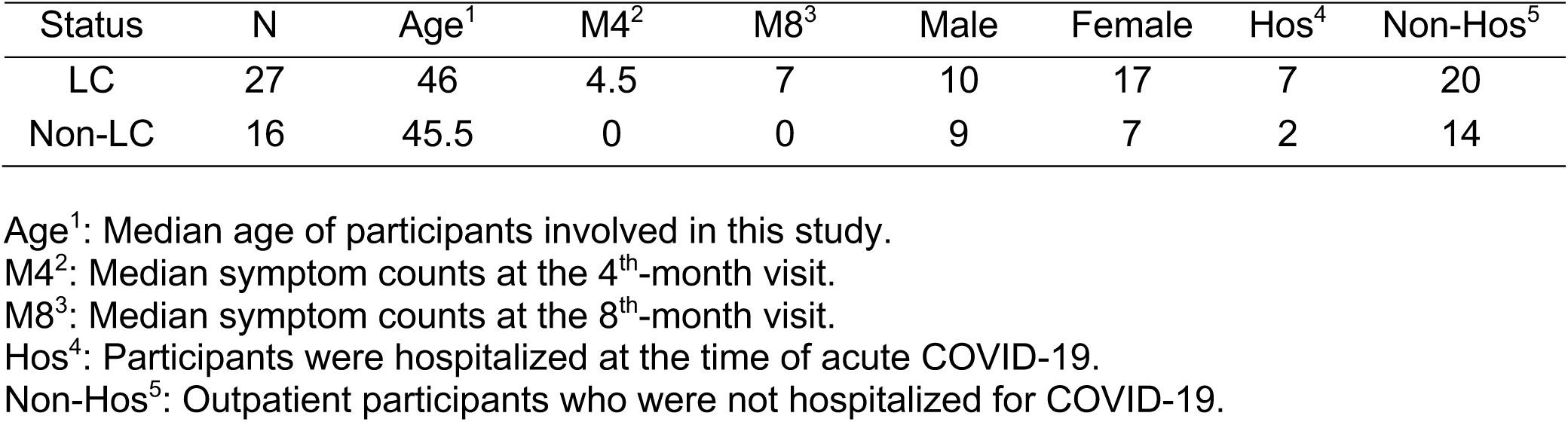
Participants’ sequelae status at the 8^th^-month visit.

**Table 2:** Detailed participants’ demographics and hospitalization status at the 8^th^-month visit.

| Participants | All | Sex |  | Hospitalized |  |
| --- | --- | --- | --- | --- | --- |
|  |  | Male | Female | Yes | No |
| N | 43 | 19 | 24 | 9 | 34 |
| Age (Median) | 46 | 53 | 43 | 46 | 48 |
| M4<sup>1</sup> (Median) | 2.5 | 0 | 5 | 7 | 2 |
| M8<sup>2</sup> (Median) | 5 | 2 | 7 | 10 | 3 |
| Female | 24 | - | - | 6 | 18 |
| Hospitalized | 9 | 3 | 6 | - | - |
| **Race (N)** |  |  |  |  |  |
| White | 25 | 13 | 12 | 0 | 25 |
| Latinx* | 11 | 6 | 5 | 7 | 4 |
| Black | 2 | 2 | 0 | 1 | 1 |
| Asian | 3 | 1 | 2 | 1 | 2 |
| NA<sup>#</sup> | 2 | 2 | 0 | 0 | 2 |

M4<sup>1</sup>: Symptom counts at the 4<sup>th</sup>-month visit.
M8<sup>2</sup>: Symptom counts at the 8<sup>th</sup>-month visit.
Latinx\*: Hispanic or Latino.
NA<sup>#</sup>: Data are not available here.

### Experimental design

SARS-CoV-2 serological analysis and five “omics” assays were performed on the same blood specimens from our cohort of LC and non-LC individuals (Fig. 1). Plasma/sera were analyzed for RBD-specific antibody levels, and for the levels of 394 analytes using the Olink platform. PBMCs from the same specimens were subjected to bulk and single-cell RNAseq (scRNAseq), as well as in-depth CD4+ and CD8+ T cell phenotyping using a 39-parameter CyTOF panel designed to simultaneously interrogate the differentiation states, activation states, effector functions, and homing properties of T cells (Table S2). Cells were phenotyped by CyTOF at baseline and following a 6-hour stimulation with SARS-CoV-2 peptides to identify and characterize SARS-CoV-2-specific T cells at the single-cell level through intracellular cytokine staining. The RNAseq/scRNAseq and Olink datasets, as well as the CyTOF datasets corresponding to total and SARS-CoV-2-specific T cells, were visualized and analyzed using a variety of integrative high-dimensional analysis approaches (Fig. 1). In total, we obtained 6 distinct datasets, enabling us to assess humoral response (serology), plasma analytes (Olink), transcriptional signatures at the bulk (RNAseq) and single-cell (scRNAseq) levels, T cell features (CyTOF), and SARS-CoV-2-specific T cell phenotypes and effector functions (CyTOF).

**Fig. 1.**
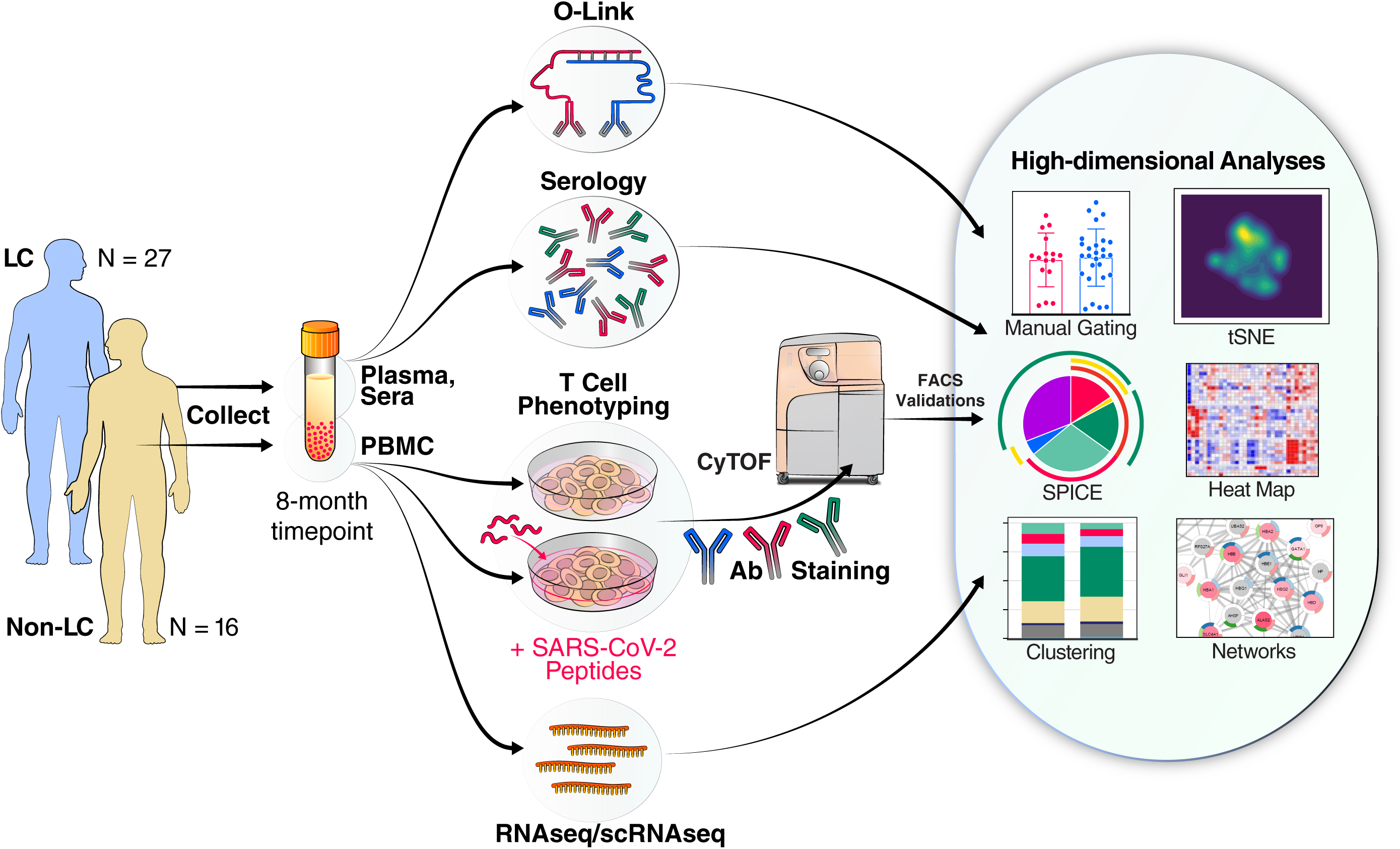
Study Design. Schematic of experimental design and data analyses. Plasma and sera from 27 individuals with Long COVID (LC) and 16 individuals without LC (Non-LC) were subjected to Olink and serological analyses. PBMCs from the same individuals were subjected to RNAseq/scRNAseq analysis, as well as to CyTOF analysis at baseline, or following a 6-hour stimulation with peptides derived from SARS-CoV-2 spike proteins (see Methods) to analyze T cell responses. The cells for CyTOF were treated with viability marker, fixed, and stained with a 39-parameter panel (Table S2) prior to analysis on a CyTOF instrument. The indicated tools on the right were then used for analyses of the resulting high-dimensional datasets.

### SARS-CoV-2-specific T cells exhibit similar frequencies and effector profiles in LC and non-LC individuals

To quantitate total and SARS-CoV-2-specific T cells, normalized events from the CyTOF datasets were gated on intact, live, singlet events, followed by gating for CD4+ and CD8+ T cells (Fig. S2A, B). The T cells were assessed for expression of all our panel’s effector molecules, which were chosen because of their roles in T-cell immunity and pathogenesis. These consisted of the cytokines IFNγ, TNFα, IL2, IL4, IL6, IL17, and MIP1β, and cytolytic markers including granzyme B and perforin (Fig. S2C, D). To determine which T cells were SARS-CoV-2-specific, we established a stringent set of rules based on the frequencies of cells expressing these effectors in samples treated or not with SARS-CoV-2 peptides (details in Methods). This analysis revealed that a combination of IFNγ, TNFα, and/or IL2 specifically identified the vast majority of SARS-CoV-2-specific CD4+ T cells (Fig. 2A, S2C), while a combination of IFNγ, TNFα, and/or MIP1β specifically identified the vast majority of SARS-CoV-2-specific CD8+ T cells (Fig. 2B, S2D).

**Fig. 2.**
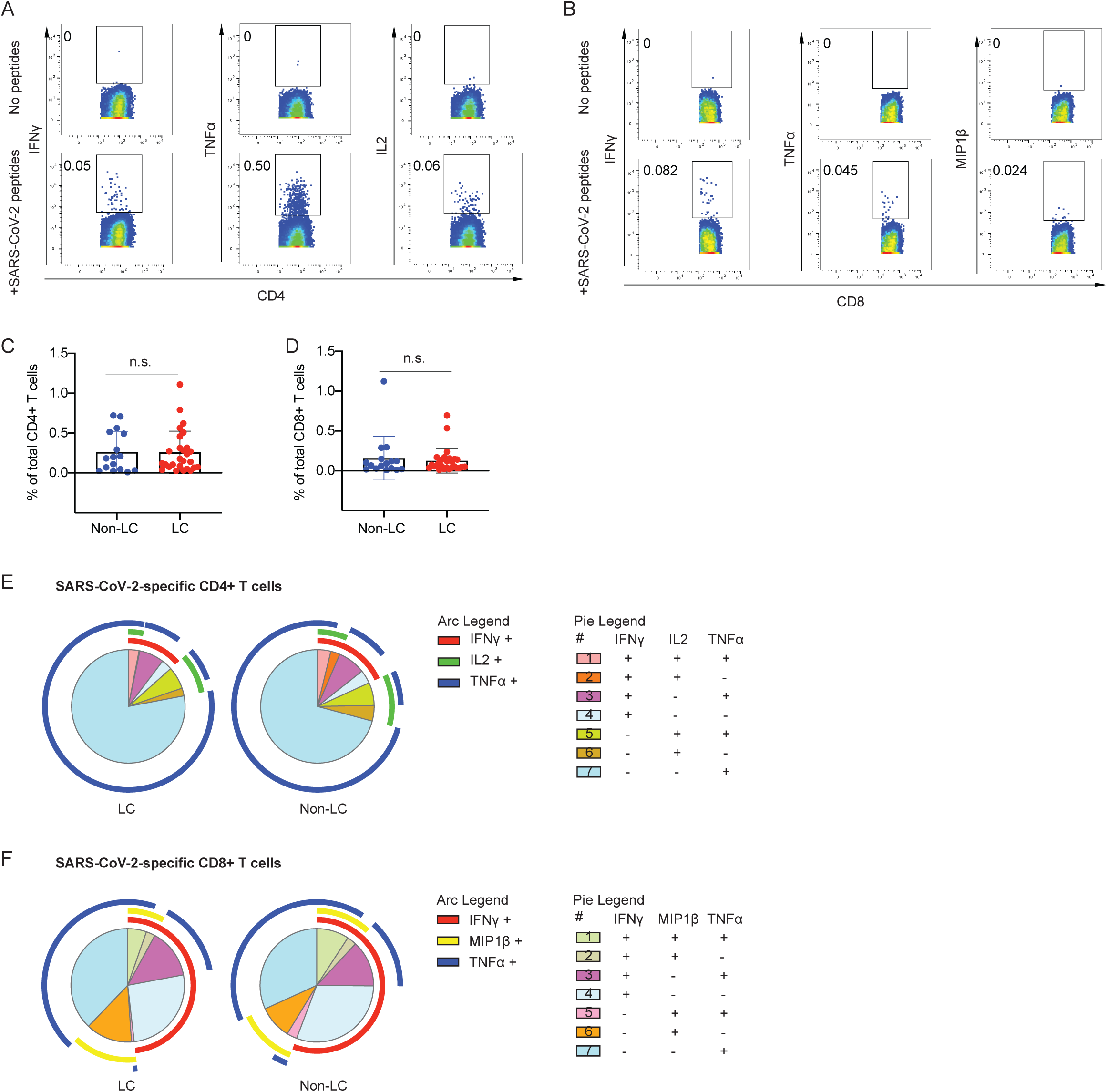
Identification and characterization of SARS-CoV-2-specific T cells in individuals from the LIINC cohort. **A.** SARS-CoV-2-specific CD4+ T cells can be identified as those producing IFNγ, TNFα, or IL2 in response to SARS-CoV-2 peptide stimulation. Cells were analyzed by intracellular cytokine staining in the absence (*top row*) or presence (*bottom row*) of SARS-CoV-2 peptides. **B.** SARS-CoV-2-specific CD8+ T cells can be identified as those producing IFNγ, TNFα, or MIP1β in response to SARS-CoV-2 peptide stimulation. Cells were analyzed by intracellular cytokine staining in the absence (*top row*) or presence (*bottom row*) of SARS-CoV-2 peptides. **C, D.** No significant differences in the magnitude of the T cell responses were observed between LC and non-LC individuals within the CD4+ (**C**) or CD8+ (**D**) T cell compartments (student’s t-tests). **E.** Analysis of polyfunctionality of SARS-CoV-2-specific CD4+ T cells. SPICE analysis revealed that polyfunctional SARS-CoV-2-specific CD4+ T cells co-expressing IFNγ, IL2, and TNFα (*category 1*) trended higher in non-LC than LC individuals albeit insignificantly (permutation test). TNFα single positive cells (*category 7*) made up the vast majority of SARS-CoV-2-specific CD4+ T cells in both LC and non-LC individuals. **F.** Analysis of polyfunctionality of SARS-CoV-2-specific CD8+ T cells. SPICE analysis revealed that polyfunctional SARS-CoV-2-specific CD8+ T cells co-expressing IFNγ, MIP1β, and TNFα (*category 1*) trended higher in non-LC than LC individuals albeit insignificantly (permutation test). TNFα single positive cells (*category 7*) made up the majority of SARS-CoV-2-specific CD8+ T cells in both LC and non-LC individuals, but to a lesser extent than for SARS-CoV-2-specific CD4+ T cells. Relative to SARS-CoV-2-specific CD4+ T cells, SARS-CoV-2-specific CD8+ T cells more frequently produced IFNγ.

Using Boolean gating, we then compared the frequencies of SARS-CoV-2-specific T cells between LC and non-LC individuals. No significant differences were observed when looking at the total population of SARS-CoV-2-specific T cells (expressing any combination of IFNγ, TNFα, IL2, and/or MIP1β) (Fig. 2C, D), or when looking at SARS-CoV-2-specific T cells producing any of the individual effector cytokines (Fig. S3A, B).

To quantitate polyfunctional cells, we implemented Simulation Program with Integrated Circuit Emphasis (SPICE) analyses. Overall, the distribution of polyfunctional SARS-CoV-2-specific T cells was similar between the LC and non-LC individuals, among both the CD4+ and CD8+ T cell compartments (Fig. 2E, F). The most polyfunctional T cells (IFNγ+TNFα+IL2+ for SARS-CoV-2-specific CD4+ T cells, and IFNγ+TNFα+MIP1β+ for SARS-CoV-2-specific CD8+ T cells) tended to be more abundant in non-LC individuals, but this trend did not reach statistical significance (Fig. 2E, F). TNFα single-positive cells made up the majority of SARS-CoV-2-specific T cells in both LC and non-LC individuals, particularly among CD4+ T cells where >50% of the responding cells singly produced this effector cytokine (Fig. 2E, F). By contrast, SARS-CoV-2-specific CD8+ T cells more frequently produced IFNγ than SARS-CoV-2-specific CD4+ T cells, independent of LC status (Fig. 2E, F). Overall, these analyses suggest that SARS-CoV-2-specific T cells have similar effector profiles in LC and non-LC individuals. However, one interesting exception was found in that IL6 was found to be induced within CD4+ T cells after SARS-CoV-2 peptide stimulation (Fig. S3C) exclusively among individuals with LC, albeit only in a small subset of these people (14%) (Fig. S3D).

### Individuals with LC exhibit different distributions of T cell subsets including in sex-dimorphic fashion

T cells can be classified not only by the effector molecules they produce, but also by T cell lineage markers. We next took advantage of the deep phenotyping capabilities of CyTOF to compare classical subset distributions among total and SARS-CoV-2-specific T cells. Naïve T cells (Tn), stem cell memory cells (Tscm), central memory T cells (Tcm), effector memory T cells (Tem), transitional memory cells (Ttm), and effector memory RA (Temra) cells were identified from both CD4+ and CD8+ T cells through sequential gating strategies (Fig. S4A, B). In addition, T follicular helper cells (Tfh) and regulatory T cells (Treg) were identified from the CD4+ compartment (Fig. S4A). Among total CD4+ T cells, the Tcm, Tfh, and Treg subsets were all significantly elevated among the individuals with LC (Fig. 3A). The other CD4+ T cell subsets were not different between the LC and non-LC groups, and among SARS-CoV-2-specific CD4+ T cells none of the subsets were significantly different (Fig. 3A, B). Among both total and SARS-CoV-2-specific CD8+ T cells, no subsets were statistically different between the LC and non-LC groups; however, there was a trend (p=0.09) for the Tem subset to be higher among total CD8+ T cells in the LC group (Fig. S5).

**Fig 3.**
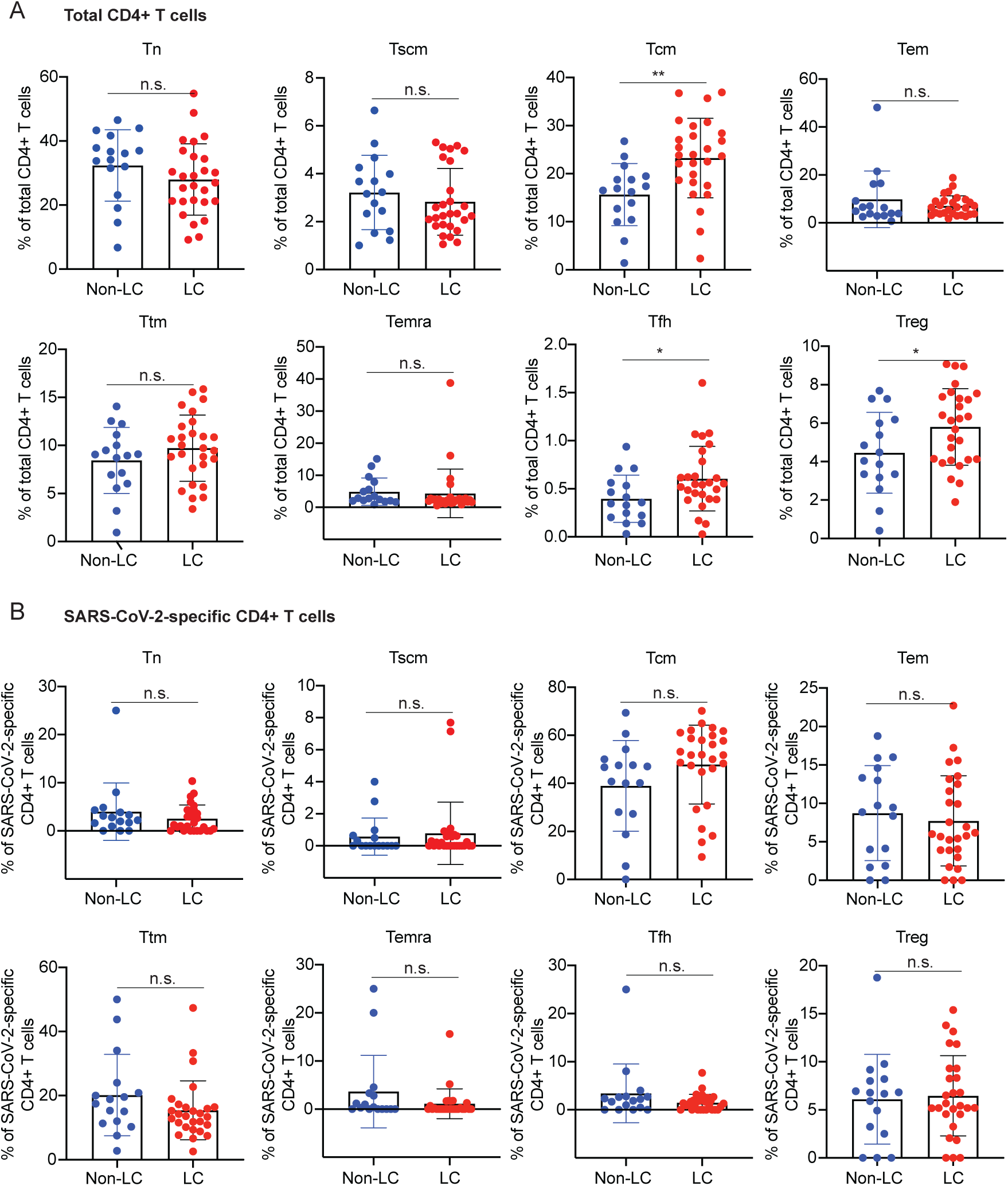
Tcm, Tfh, and Treg frequencies differ between LC and Non-LC individuals. **A.** CD4+ T cell subset analysis reveals higher proportions of Tcm, Tfh, and Treg in LC vs. non-LC individuals. **p<0.01, *p<0.05 (student’s t-test). **B.** No significant differences were observed in the proportion of the indicated SARS-CoV-2-specific CD4+ T cell subsets between LC vs. non-LC individuals. Phenotypic definition of subsets were as follows: naïve T cells (Tn): CD45RA+CD45RO-CCR7+CD95-, stem cell memory T cells (Tscm): CD45RA+CD45RO-CCR7+CD95+, central memory T cells (Tcm): CD45RA-CD45RO+CCR7+CD27+, effector memory T cells (Tem): CD45RA-CD45RO+CCR7-CD27-, transitional memory T cells (Ttm): CD45RA-CD45RO+CCR7-CD27+, effector memory RA T cells (Temra): CD45RA+CD45RO-CCR7-, T follicular helper cells (Tfh): CD45RA-CD45RO+PD1+CXCR5+, and regulatory T cells (Treg): CD45RA-CD45RO+CD127-CD25+.

To examine T cell distribution between LC and non-LC individuals using not only the T cell lineage markers, but all the phenotyping and effector markers analyzed in our CyTOF panel, we examined the mean expression levels of all these markers among both total and SARS-CoV-2-specific T cells. Following multiple correction adjustment, no antigens were significantly differentially expressed between the individuals with or without LC (Fig. S6-S9). As excess inflammation has been implicated in the context of both severe COVID-19 and LC, we additionally performed manual gating to assess for evidence of increased T cell activation in individuals with LC. This revealed no significant differences in the percentages of cells expressing acute activation markers CD38, HLADR, and Ki67 between individuals with and without LC (Fig. S10). T cells expressing all three activation markers were also not differentially represented among the two groups of individuals (Fig. S10). To analyze the complexity of our high-dimensional datasets in ways not captured through such one-dimensional analyses of individual marker expression, we performed clustering analyses. CD4+ and CD8+ T cells fell into six (Clusters A1-A6) and five (Clusters B1-B5) clusters, respectively (Fig. S11A, S12A), that did not differ significantly between the LC and non-LC groups, except when we stratified the participants by sex. Among CD4+ T cells, cluster A1 was significantly over-represented in LC than non-LC females (but not males), while cluster A4 was significantly over-represented in LC than non-LC males (but not females) (Fig. S11B). Cluster A1 was composed of naïve CD4+ T cells, and expressed low levels of activation markers and inflammatory tissue homing receptors, and high levels of the lymph node homing receptors (Fig. S11C). By contrast, cluster A4 was composed of terminally differentiated effector memory CD4+ T cells and expressed high levels of receptors associated with homing to inflamed tissues but not those associated with homing to lymph nodes (Fig. S11D). They also expressed elevated levels of the cytolytic markers including perforin and granzyme B (Fig. S11D). Among CD8+ T cells, cluster B1 was significantly under-represented in LC while cluster B2 was significantly over-represented, but only among females (Fig. S12B). Interestingly, the phenotypic features of cluster B1 mirrored those of cluster A1, while the features of cluster B2 mirrored those of cluster A4 (Fig. S12 C, D). These observations, together with the observation that cluster A4 trended higher in the female LC group, suggest that female individuals with LC harbor relatively low frequencies of resting naïve T cells expressing low levels of inflammatory tissue-homing receptors, and high frequencies of terminally differentiated effector memory T cells expressing inflammatory tissue homing receptors and cytolytic markers; this was true among both CD4+ and CD8+ T cells. More broadly, the results suggest that there are sex-dimorphic differences in the subset distribution of T cells between LC and non-LC individuals.

### Preferential expression of some tissue-homing receptors on CD4+ T cells from individuals with LC

We then focused on the phenotypic features of SARS-CoV-2-specific T cells. Although clustering analyses did not reveal significant differences between the LC vs. non-LC groups (not shown), contour-based tSNE visualization revealed that the cells from the LC vs. non-LC groups tended to concentrate in different areas suggesting some phenotypic differences (Fig. 4A, 5A). Focusing first on the SARS-CoV-2-specific CD4+ T cells, we found that the chemokine receptors CXCR4, CXCR5, and CCR6 were all expressed at higher levels in the cells from the LC as compared to the non-LC individuals (Fig. 4B). Gating on SARS-CoV-2-specific CD4+ T cells co-expressing various pairs of these chemokine receptors revealed that those that were CXCR4+CXCR5+ and CXCR5+CCR6+ were significantly increased, while those that were CXCR4+CCR6+ trended higher, in individuals with LC (Fig. 4C). Interestingly, this same pattern was observed among total CD4+ T cells (Fig. 4D), in a manner confirmed by FACS (FACS-based assessment of receptor expression significantly associated with measurements made by CyTOF, Fig. S13A), but now where the elevated frequencies of CXCR4+CCR6+ CD4+ T cells in LC also reached statistical significance (Fig. S13B, C). These results together suggest that expression of tissue-homing receptors is elevated in CD4+ T cells – including but also beyond those with specificity for SARS-CoV-2 – in individuals with LC.

**Fig. 4.**
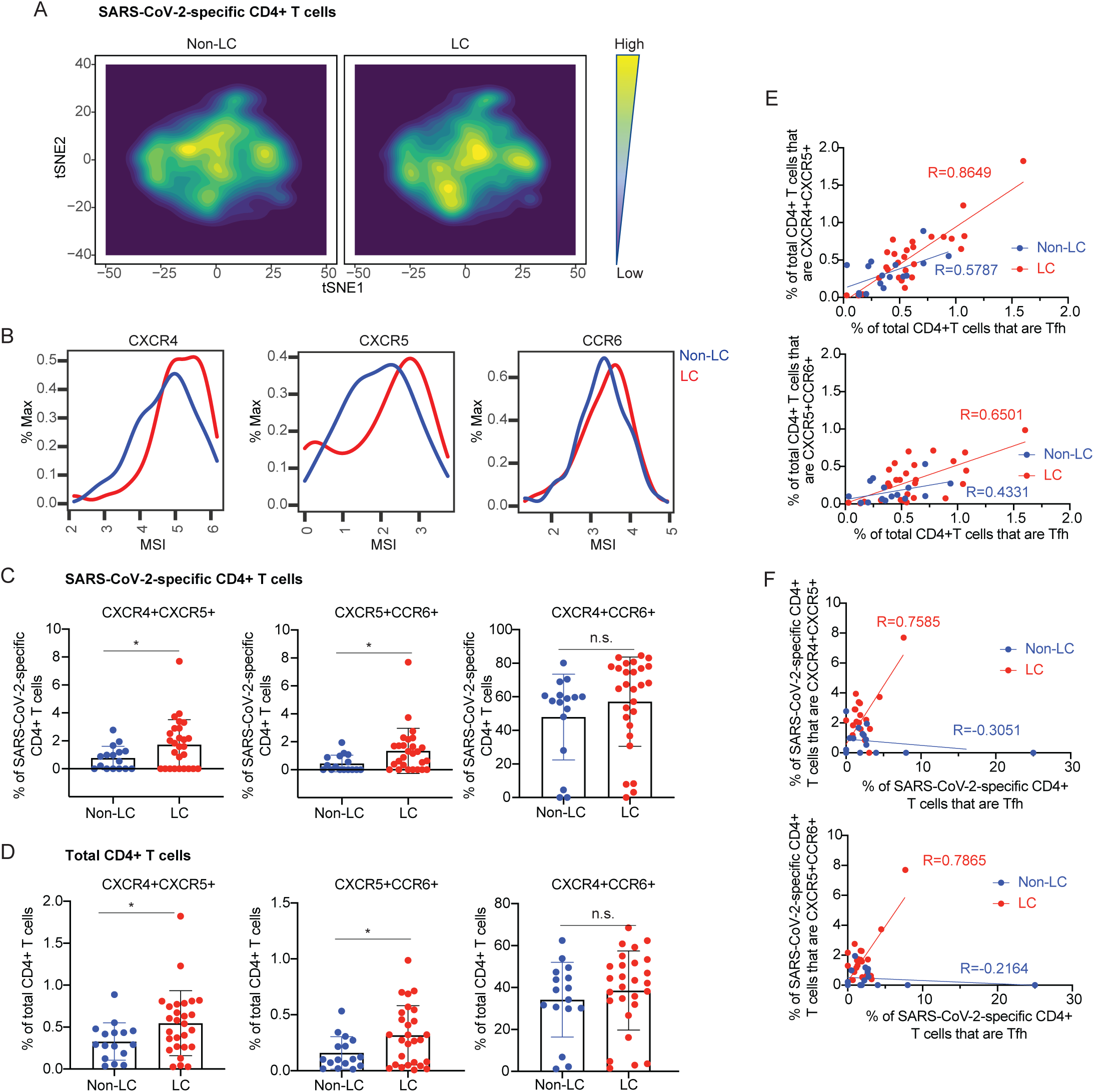
SARS-CoV-2-specific CD4+ T cells from individuals with LC preferentially express homing receptors associated with migration to inflamed tissues. **A.** tSNE contour depiction of SARS-CoV-2-specific CD4+ T cells from LC and non-LC individuals, highlighting different distribution of cells from the two groups. **B.** Expression of the chemokine receptors CXCR4, CXCR5, and CCR6 are elevated in SARS-CoV-2-specific CD4+ T cells from LC as compared to non-LC individuals. MSI corresponds to mean signal intensity of the indicated markers’ expression level, reported as arcsinh-transformed CyTOF data as detailed in the Methods. **C, D.** CXCR4+CXCR5+ and CXCR5+CCR6+ SARS-CoV-2-specific *(C)* and total *(D)* CD4+ T cells are significantly elevated in LC as compared to non-LC individuals, while their CXCR4+CCR6+ counterparts trended higher in the LC group. *p<0.05 (student’s t-test). **E**. The frequencies of CXCR4+CXCR5+ and CXCR5+CCR6+ CD4+ T cells are significantly positively associated (p<0.0001 for CXCR4+CXCR5+, p<0.001 for CXCR5+CCR6+) with the frequencies of Tfh in LC, but not non-LC, individuals. **F**. The frequencies of SARS-CoV-2-specific CXCR4+CXCR5+ and CXCR5+CCR6+ CD4+ T cells are significantly positively associated (p<0.0001 for CXCR4+CXCR5+, p<0.0001 for CXCR5+CCR6+) with the frequencies of SARS-CoV-2-specific Tfh in LC, but not non-LC, individuals. In *panels E-F*, correlation estimates were identified as R values (Pearson correlation coefficients).

**Fig. 5.**
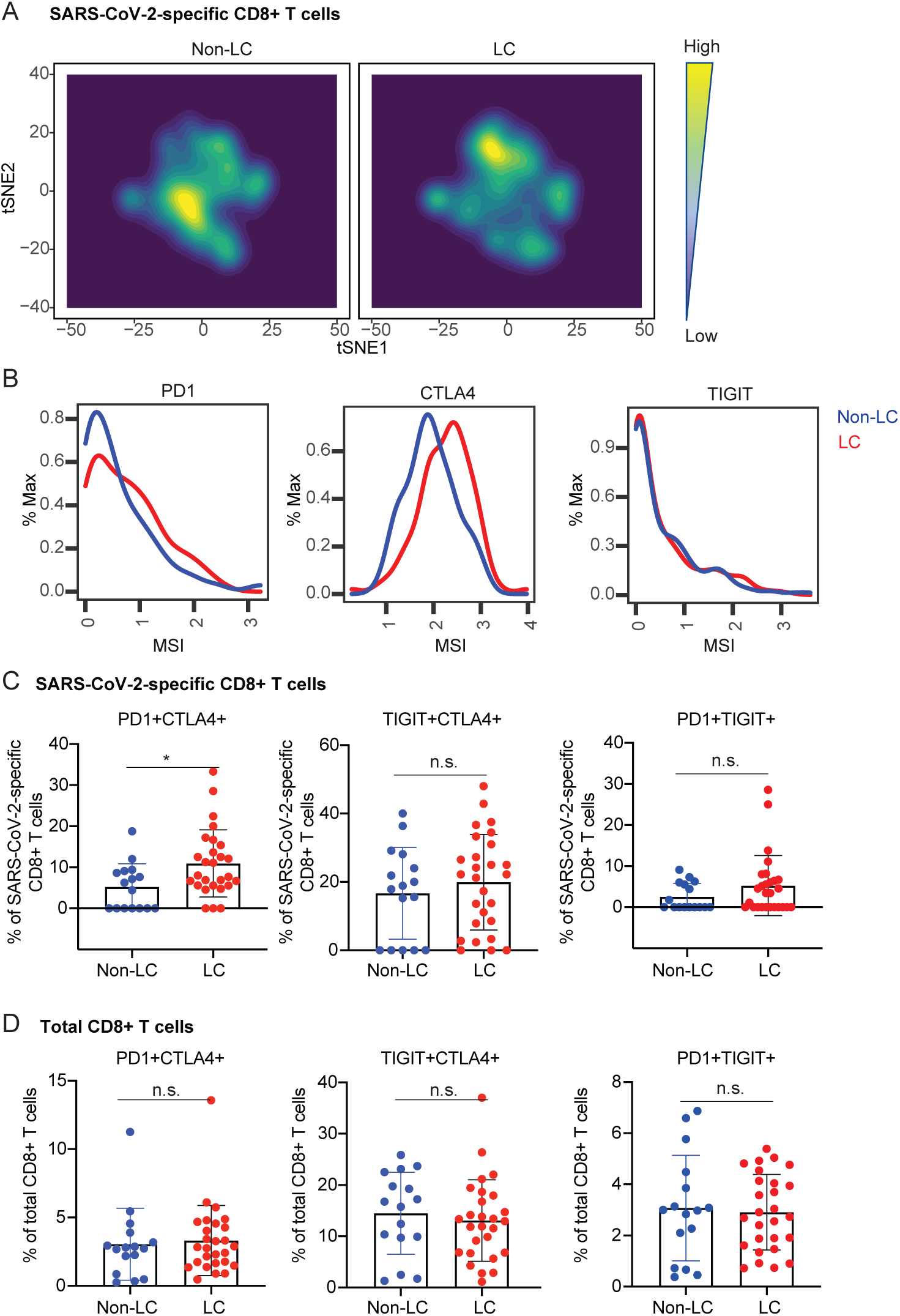
SARS-CoV-2-specific CD8+ T cells from individuals with LC preferentially express exhaustion markers PD1 and CTLA4. **A.** tSNE contour depiction of SARS-CoV-2-specific CD8+ T cells from LC and non-LC individuals, highlighting different distribution of cells from the two groups. **B.** Expression of exhaustion markers PD1 and CTLA4, but not TIGIT, are elevated on SARS-CoV-2-specific CD8+ T cells from LC as compared to non-LC individuals. MSI corresponds to mean signal intensity of the indicated markers’ expression level. **C, D.** PD1+CTLA4+ cells are significantly enriched among SARS-CoV-2-specific CD8+ T cells (**C**) but not total CD8+ T cells (**D**) in LC as compared to non-LC individuals. By contrast, TIGIT+CTLA4+ and PD1+TIGIT+ total and SARS-CoV-2-specific CD8+ T cells were equivalently distributed between LC and non-LC individuals. *p<0.05 (student’s t-test).

It was intriguing to us that CXCR5 was a common marker among the CXCR4+CXCR5+, CXCR5+CCR6+, and Tfh CD4+ T cell subsets, both of which were found to be significantly elevated in LC (Fig. 4, S13). This prompted us to examine whether there was an association between the frequencies of Tfh and these other chemokine receptor-expressing subsets. Intriguingly, we observed significant associations between the frequencies of Tfh and these subsets, particularly for the LC group (Fig. 4E). When we performed this correlation analysis among SARS-CoV-2-specific instead of total CD4+ T cells, significant positive associations were observed only among the LC group, while in the non-LC group there was a negative association trend (Fig. 4F).

### SARS-CoV-2-specific CD8+ T cells from individuals with LC preferentially co-express the checkpoint molecules PD1 and CTLA4

A similar manual inspection of CD8+ T cell data revealed the checkpoint/exhaustion markers PD1 and CTLA4 to be expressed at elevated levels on SARS-CoV-2-specific CD8+ T cells from the LC as compared to non-LC individuals, while the exhaustion marker TIGIT was not differentially expressed (Fig. 5B). Consistent with these data, SARS-CoV-2-specific CD8+ T cells that were PD1+CTLA4+, but not those that were TIGIT+CTLA4+ or PD1+TIGIT+, were significantly elevated in individuals with LC (Fig. 5C). Interestingly, however, frequencies of total PD1+CTLA4+ CD8+ T cells were not different between the LC and non-LC individuals (Fig. 5D). These results were confirmed by FACS analyses of the same patient specimens (Fig. S14). Although PD1 and CTLA4 are also activation markers, the fact that we didn’t see elevation of acute activation markers CD38, HLADR, or Ki67 on SARS-CoV-2-specific CD8+ T cells from individuals with LC (Fig. S10) suggests that what is being observed is not simply elevated T cell activation. These results together suggest that LC-derived CD8+ T cells recognizing SARS-CoV-2 uniquely exhibit phenotypic features of exhaustion (but not acute activation), as reflected by co-expression of PD1 and CTLA4, perhaps due to persistence of SARS-CoV-2 antigen that cannot be eliminated in individuals with LC.

### Individuals with LC exhibit a mis-coordinated T and antibody response

Serological analysis revealed significantly higher SARS-CoV-2 RBD antibody levels in the LC group than in the non-LC group (Fig. 6A). Interestingly, the two individuals with LC with the highest antibody levels (green oval) were not those that had the highest frequencies of exhausted (PD1+CTLA4+) SARS-CoV-2-specific CD8+ T cells (purple oval) (Fig. 6B), even though both features are consistent with a persistent SARS-CoV-2 reservoir. Interestingly, however, the individuals with the highest frequencies of exhausted SARS-CoV-2-specific CD8+ T cells did have the lowest frequencies of SARS-CoV-2-specific CD4+ Treg cells, and the frequencies of these two subsets of cells negatively correlated in LC but not non-LC individuals (Fig. 6C). When we assessed the association between antibody levels and SARS-CoV-2-specific T cell frequencies, we found a significant (p=0.0418 for CD4, p=0.0007 for CD8) positive correlation, but only in the non-LC individuals (Fig. 6D, F). The frequencies of SARS-CoV-2-specific Tfh cells also significantly (p=0.0014) positively correlated with antibody responses in non-LC individuals but not in individuals with LC (Fig. 6E). These results suggest that a mis-coordinated humoral and cell-mediated response, previously implicated in severe COVID-19 ^38^, may also be a hallmark of LC.

**Fig. 6.**
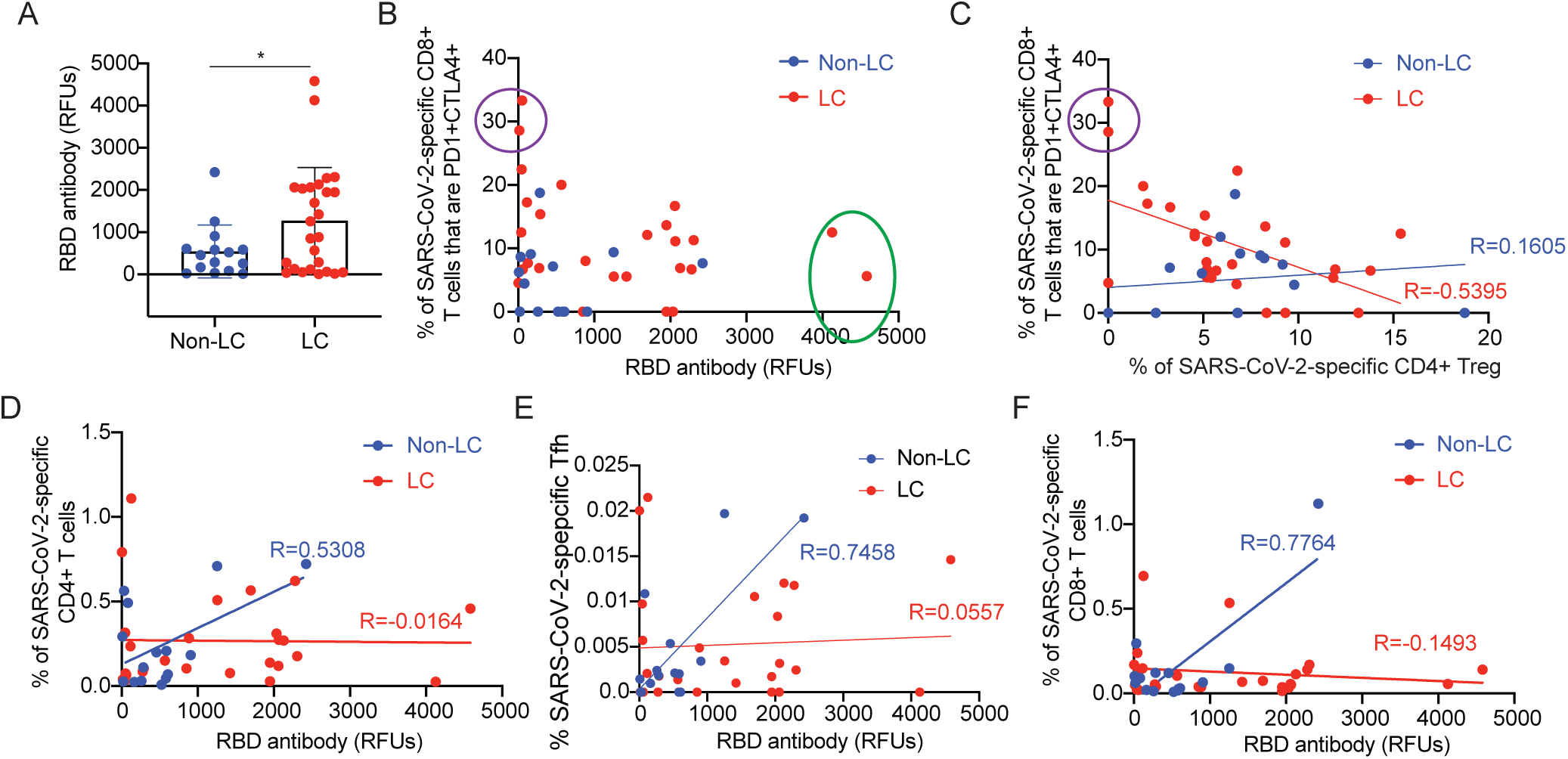
Dis-coordinated humoral and adaptive immunity in individuals with LC. **A.** SARS-CoV-2 spike RBD antibody levels are elevated in LC as compared to non-LC individuals. *p<0.05 (student’s t-test). **B.** Individuals with LC harboring the highest humoral response (green oval) are not those exhibiting highest levels of exhausted PD1+CTLA4+ SARS-CoV-2-specific CD8+ T cells (purple oval). **C**. Frequencies of PD1+CTLA4+ SARS-CoV-2-specific CD8+ T cells and SARS-CoV-2-specific CD4+ Treg cells are negatively associated only in individuals with LC. **D-F.** SARS-CoV-2 spike RBD antibody levels are significantly positively associated with the frequencies of SARS-CoV-2-specific CD4+ T cells *(**D**)*, SARS-CoV-2-specific Tfh *(**E**)*, and SARS-CoV-2-specific CD8+ T cells *(**F**)* in non-LC individuals, but not in individuals with LC. In *panels D-F*, the correlation estimates (identified as R in the figures) for the non-LC group were significantly (p<0.05 for the CD4+ T cells, p<0.01 for Tfh cells, p<0.001 for the CD8+ T cells) different from zero while the corresponding estimates were n.s. (p>0.05) for the LC group (Pearson r t-tests).

### Individuals with LC exhibit global alterations in PBMCs reflecting immune and non-immune dysregulation

To determine whether the differences between LC and non-LC individuals extended beyond T cells and the humoral response, we examined the transcriptome of the PBMCs. Assessing for total changes in gene expression in PBMCs by bulk RNAseq, we found only two genes that remained significantly differentially expressed after multiple comparison adjustments: OR7D2 (Olfactory Receptor 7D2) and ALAS2 (5’-Aminolevulinate Synthase 2), both of which were over-expressed in the individuals with LC (Fig. 7A). Interestingly, this was observed in a non-overlapping manner in that the four individuals with highest OR7D2 expression did not overlap with the four individuals with highest ALAS2 expression. OR7D2 encodes a G-protein-coupled receptor that responds to odorant molecules in the nose, while ALAS2 encodes a protein that catalyzes the first step in heme synthesis, defects in which can lead to anemia.

**Fig. 7.**
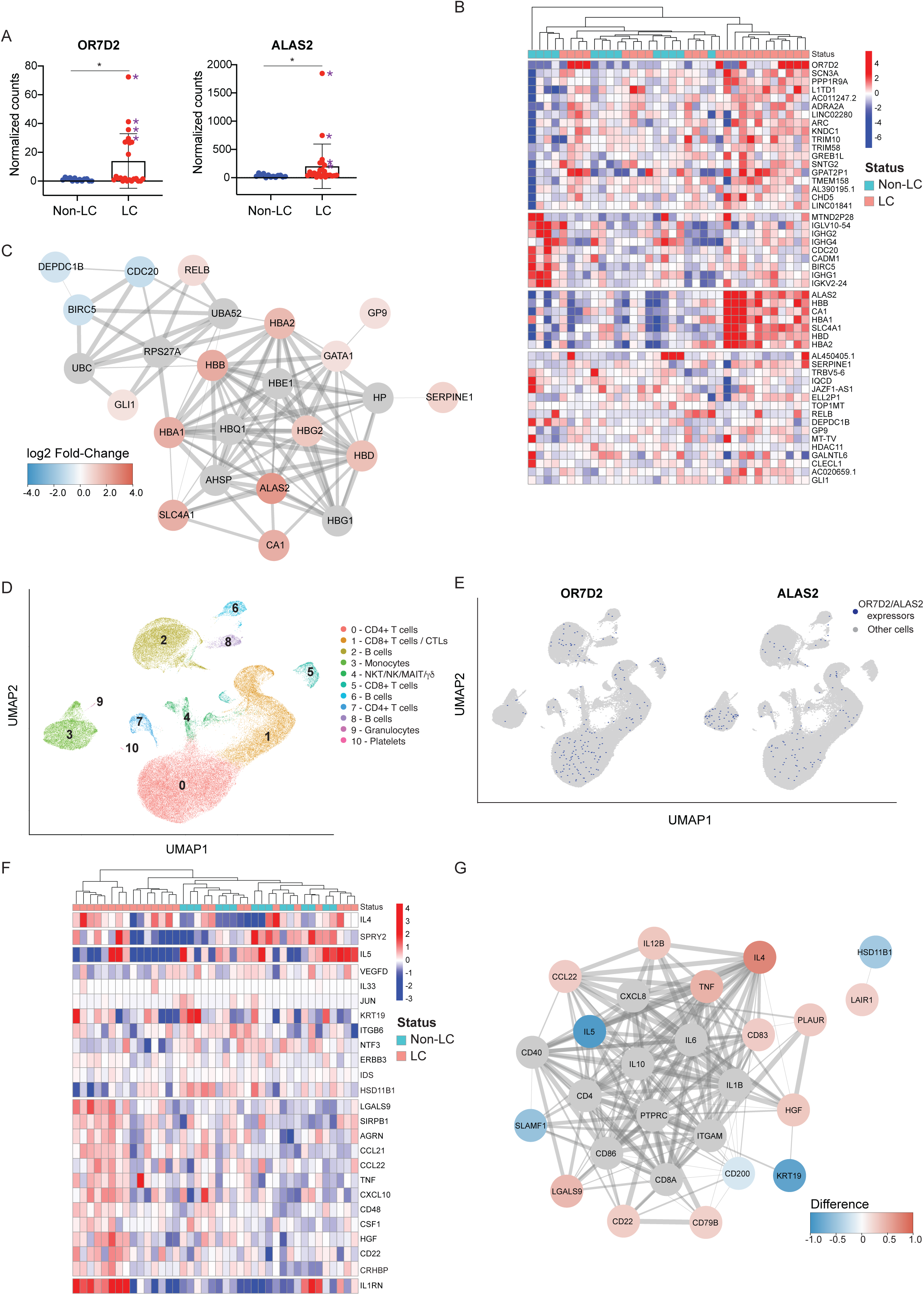
Global differential gene and gene product expression in participants with and without LC. **A.** Relative gene expression levels of top two significantly differentially expressed genes (DEGs) from bulk RNAseq analysis of LC vs. non-LC individuals. OR7D2 corresponds to Olfactory Receptor family 7D2 (log_2_ fold-change=3.63), and ALAS2 to 5’Aminolevulinate Synthase 2 (log_2_ fold-change=2.58). *p < 0.05 (Wald test, with Benjamini-Hochberg correction). Purple asterisks identify the female donors selected for follow-up scRNAseq analyses. **B.** Clustered heatmap of the top 50 DEGs in PBMCs in LC compared to non-LC individuals. Genes are grouped into k-clusters based on similarity. Note four modules of gene expression, with the second corresponding to immunoglobulin and T cell genes (under-expressed in LC), and the third corresponding to heme synthesis and carbon dioxide transport (over-expressed in LC). **C.** Network mapping of related DEGs from bulk RNAseq analysis. Each node corresponds to a gene, and colors of nodes indicate the extent of change as indicated in the heatmap scale bar, with red corresponding to upregulation in individuals with LC, and blue corresponding to downregulation in individuals with LC. Edges depict the functional relevance between pairs of genes, where the thickness of the edge corresponds to the confidence of the evidence. The highly networked nature of the indicated genes supports their association with LC. **D.** UMAP of annotated clusters from samples analyzed by scRNAseq. **E.** Both OR7D2 and ALAS2 are broadly expressed among PBMC subsets of individuals with LC. Shown are UMAP depictions of cells expressing (*blue*) or not expressing (*grey*) OR7D2 or ALAS2 as indicated. **F.** Clustered heatmap of the top 25 differentially expressed plasma proteins from Olink Proximity Extension Assay with markers grouped into k-clusters based on similarity. Note a dominant module of inflammatory-related genes including LGALS9, CCL21, CCL22, TNF, CXCL10, and CD48. **G.** Network mapping of related differentially expressed proteins as detected by Olink. Graph representations are as described in *panel C*. Note the simultaneous over-expression of IL4 and CCL22 (in red) with under-expression of IL5 (in blue), all three proteins of which are involved in Th2 immune responses.

Further analysis of the RNAseq datasets revealed that a number of genes other than OR7D2 and ALAS2 were upregulated in individuals with LC, although not significantly so after conservative adjustments for multiple comparisons; these included a module of genes that regulate heme synthesis and carbon dioxide transport (ALAS2, HBB, CA1, HBA1, SLC4A1, HBD, HBA2) (Fig. 7B, S15A). By contrast, a module consisting of immunoglobin kappa, lambda, and heavy chain genes, along with BIRC5, which plays an important role in T cell survival and function ^39^, were more highly expressed in non-LC individuals (Fig. 7B, S15A). Gene ontology analysis revealed that genes from both of these modules were highly networked together (Fig. 7C), strongly suggesting that these genes are indeed linked to LC.

In order to determine which subsets of cells were expressing our top differentially expressed genes (DEGs) OR7D2 and ALAS2, as well as to gain a more granular view of the transcriptional features of cells from individuals with LC, we selected a subset of the participants for further analyses by scRNAseq. Intriguingly, females were vastly over-represented among the LC individuals expressing high levels of OR7D2 or ALAS2: the top five OR7D2 expressors were all female, and among the top six ALAS2 expressors, five were female (the third highest ALAS2 expressor was male). To avoid sex-associated confounders, we limited our scRNAseq studies to female donors. We selected the four LC females with the highest OR7D2 expression and the four LC females with the highest ALAS2 expression (purple asterisks in Fig. 7A), and four randomly selected non-LC females for comparison. Integrative analysis of all 12 donors identified 11 clusters of cells, including subsets of CD4+ T cells, CD8+ T cells, B cells, monocytes, and innate-like cells (Fig. 7D). In addition, small numbers of granulocytes and platelets were identified; their low numbers were expected as PBMCs and not whole blood were analyzed. While the major clusters did not differ in frequency between the LC and non-LC groups, the granulocyte cluster was significantly less abundant (p=0.006) while the platelet cluster was significantly more abundant (p=0.01) in individuals with LC. Consistent with lack of major perturbations at the transcriptional level, visualization of the datasets by LC vs. non-LC status (Fig. S16A), and by the two groups of LC (OR7D2^high^ and ALAS2^high^) (Fig. S16B), did not reveal profound differences. Consistent with the bulk RNAseq data, OR7D2 was expressed at the highest levels in the OR7D2^high^ LC cells, and ALAS2 was expressed at the highest levels in the ALAS2^high^ LC cells (Fig. S16C). When we assessed which clusters of cells expressed OR7D2 and ALAS2, we found that these transcripts were broadly expressed in all subsets except the granulocyte and platelet clusters (Fig. 7E, S16D), with the caveat that the low numbers of cells in the latter two clusters likely precluded ability to detect these transcripts. Interestingly, among clusters expressing OR7D2 or ALAS2, only a small fraction of cells (<0.5%) expressed these genes (Fig. S16D), suggesting that small numbers of cells harboring high levels of OR7D2/ALAS2 were responsible for the original identification of these DEGs from the bulk RNAseq analysis. Clusters expressing OR7D2 generally expressed this gene at the highest levels in the OR7D2^high^ LC group (Fig. S16E). Similarly, clusters expressing ALAS2 generally expressed this gene at the highest levels in the ALAS2^high^ LC group (Fig. S16F). Interrogating cluster-specific gene expression beyond just OR7D2 and ALAS2 expression revealed only 3 genes as significantly differentially expressed between the LC and non-LC groups, two within a CD4+ T cell cluster and one within the monocyte cluster (Table S3). Notably, however, using a less stringent cutoff of p<0.1, we found that the activation marker GITR was more highly expressed (p=0.07) in a cluster of CD4+ T cells from the individuals without LC. Reasoning that LC is heterogeneous and that the OR7D2^high^ and ALAS2^high^ groups may represent different subtypes of individuals with LC, we then compared these groups individually with our non-LC controls. This revealed 35 DEGs in the OR7D2^high^ group (Table S4), and 14 DEGs in the ALAS2^high^ group (p<0.05) (Table S5). With a less stringent cutoff of p<0.1, a number of additional immune-related genes of interest were additionally identified, including downregulation of the Th17 lineage marker RORC (p=0.06) and upregulation of immune checkpoint KLRG1 (p=0.08) in subsets of CD4+ T cells from the OR7D2^high^ LC as compared to non-LC individuals. Overall, our scRNAseq results support immune dysregulation in LC, and validate the OR7D2 and ALAS2 bulk RNAseq data.

### Individuals with LC exhibit global alterations in sera reflecting immune dysregulation and inflammation

Finally, we implemented Olink analysis to characterize the proteome of the sera from our participant specimens. This also revealed global changes associated with LC, including a module consisting of elevated levels of proteins associated with inflammation (LGALS9, CCL21, CCL22, TNF, CXCL10, CD48) (Fig. 7F, S15B). Proteins associated with immune regulation (IL1RN, CD22) were also elevated in individuals with LC (Fig. 7F, S15B). Interestingly, although IL4 and IL5 are both canonical cytokines for Th2 responses, these two cytokines exhibited very different expression patterns (Fig. 7F, S15B), and individuals with LC overall exhibited elevated levels of IL4 yet lower levels of IL5 (Fig. 7G). CCL22, a ligand for the Th2 marker CCR4, was expressed at elevated levels in individuals with LC (Fig. 7G). Together, these results suggest an elevated yet mis-coordinated Th2 response (elevated IL4 and CCL22 but diminished IL5) in individuals with LC. Intriguingly, IL4, but not IL5 or CCL22, significantly positively associated with the percentages of CXCR4+CXCR5+ and CXCR5+CCR6+ CD4+ T cells (more frequent in LC, Fig. 4), but only in the individuals with LC (Fig. S17). By contrast, individuals without LC exhibited a significant negative association (Fig. S17). As for the RNAseq data, networking of genes from the inflammatory and immunoregulatory modules, as well as from the Th2 markers IL4, IL5, and CCL22, suggests a biologically-relevant plausible association of all of these genes with LC. Overall, our findings suggest that LC is associated with unique, and likely complex, global immune dysregulation.

## Discussion

Using multiple “omics” analytical approaches on specimens from individuals exhibiting consistent LC trajectories, we demonstrate that individuals with LC exhibit perturbations in both total and SARS-CoV-2-specific T cells, which manifests at a global level as mis-coordination between the two main arms of adaptive immunity and overall changes in gene expression. In this analysis, we took care to limit several confounders that often constrain studies of LC. First, we carefully selected a cohort of individuals who consistently met the case definition for LC over an 8-month period and compared them with individuals who, when measured in the same way, using the same study instruments at the same timepoints, consistently demonstrated complete recovery. Second, to avoid surveillance bias, all assays were applied on samples from the same timepoint (8 months post-COVID), and we chose this relatively late timepoint so that we would not be confounded by individuals only exhibiting shorter-term LC (e.g., which resolve spontaneously after 4-6 months). Third, we restricted our analysis to only those individuals who prior to the time of sampling had not yet received a SARS-CoV-2 vaccine and who had not had a known or suspected SARS-CoV-2 re-infection, as either could markedly affect our SARS-CoV-2-specific antibody and T cell measurements.

Our CyTOF data revealed profound changes in classical subset distribution among total CD4+ T cells in individuals with LC, specifically a significantly higher proportion of CD4+ Tcm, Tfh, and Treg cells. Elevated frequencies of Tcm in LC have been reported previously ^30^, although another group reported the opposite observation that Tcm frequencies were decreased in LC ^20^. The reason for these discrepancies is not clear, but may reflect the composition of the clinical cohorts studied: while our study and the other one which also reported elevated Tcm cells examined only non-vaccinated individuals, the study where Tcm were decreased included some individuals who received a SARS-CoV-2 vaccine prior to sampling. Higher frequencies of Tfh and Treg cells in individuals with LC have, to our knowledge, not been previously reported. Interestingly, however, a prior study reported that elevated frequencies of activated Treg cells during acute SARS-CoV-2 infection predicted development of LC two to three months later ^12^, which together with our findings is consistent with Tregs being involved in both LC initiation and maintenance.

Elevated frequencies of Tcm, Tfh, and Treg in individuals with LC indicate an ongoing immune response persisting at 8 months post-infection. This immune response, however, may not necessarily be directed against SARS-CoV-2, and could potentially be directed against other viruses (*e.g.,* reactivated EBV or other herpes viruses) or auto-antigens ^12^. Indeed, we did not find significantly higher magnitude of the SARS-CoV-2-specific T cell response as determined by intracellular cytokine staining in individuals with LC, consistent with prior observations reported from the activation-induced marker (AIM) assay ^30^. We also did not find individuals with LC to harbor more polyfunctional SARS-CoV-2-specific T cells, and in fact polyfunctionality trended lower in both the CD4 and CD8 compartments. At the same time, other studies have reported higher ^26^ or lower ^31^ SARS-CoV-2-specific T cell responses in the context of LC. Discrepancies may stem from differences in the LC cohorts analyzed, and in the assays used to quantitate T cell responses (including the SARS-CoV-2 proteins examined, and the approaches used to identify responding cells). We note that our approach was comprehensive in that we monitored expression levels of 10 different effectors, and settled on a subset of five of these (IFNγ, TNFα, IL2, MIP1β, IL6) using strict criteria, to define SARS-CoV-2-specific T cells.

One aspect highly consistent between all studies to date is the ability to detect SARS-CoV-2-specific T cells in both LC and non-LC individuals, months after infection. This could simply be attributed to the long-term persistence of memory T cells elicited by SARS-CoV-2, but may also indicate the persistence of a long-lived tissue viral reservoir as has been documented^16^. Indeed, we found that in LC relative to non-LC individuals, SARS-CoV-2-specific CD8+ T cells, but not total CD8+ T cells, more frequently expressed the exhaustion markers PD1 and CTLA4, which is consistent with ongoing stimulation with viral antigens. Also in support of a potential persistent reservoir is our observation of higher SARS-CoV-2 antibody levels in LC as compared to non-LC individuals, which has also been previously seen with Spike-specific IgG levels ^30^. Interestingly, our data revealed that the individuals with the highest frequencies of SARS-CoV-2-specific CD8+ T cells co-expressing the exhaustion markers PD1 and CTLA4 were not those with the highest SARS-CoV-2 antibody levels, suggesting that there may be multiple endotypes of LC being driven by persisting virus. Consistent with this possibility, a recent RNAseq study identified two types of LC: one being driven by high expression of Ig-related genes, and the other being associated with low levels of Ig-related genes ^40^. Based on these observations as well as case reports of improvement in LC symptoms following antiviral treatment ^41, 42, 43^, nirmatrelvir-ritonavir treatment as an antiviral strategy to clear this putative LC-associated SARS-CoV-2 reservoir is underway (NCT05576662, NCT05595369, NCT05668091). Future studies could evaluate other antivirals or monoclonal antibodies, and in light of our PD1 and CTLA4 expression data might consider incorporating checkpoint inhibition in conjunction with antivirals to reinvigorate T cells’ ability to help eliminate residual viremia.

One intriguing aspect of LC that emerged from our study is sex-dimorphism in T cell phenotypes. This perhaps is not so surprising given the different trajectories of COVID-19 between males and females ^44^ and the observation that LC is more common in females ^5, 45^. Our data revealed that, among females, a subset of activated and cytotoxic T cells was more elevated in LC than in non-LC individuals; intriguingly, the opposite pattern was observed in males. The presence of cytotoxic T cells has been associated with gastrointestinal LC symptoms ^12^ and it will be of interest in future studies to establish whether biological sex impacts LC-associated cytotoxic T cell function. We also found in our RNAseq analysis that individuals expressing the highest levels of the top LC-associated DEGs (OR7D2 and ALAS2) were heavily biased towards female sex, again emphasizing the sex dimorphism of LC. Intriguingly, biological sex was recently shown to manifest in the context of differential responses to influenza vaccines after COVID-19 convalescence ^46^, although individuals with LC were not examined therein.

While SARS-CoV-2-specific CD8+ T cells from individuals with LC showed phenotypic features of exhaustion, their CD4+ counterparts preferentially expressed the tissue-homing receptors CXCR4, CXCR5, and CCR6. Of note, these receptors can all direct immune cells to the lung, and CXCR4 is of particular interest as its expression on bystander T cells has been associated with severe/fatal COVID-19 ^34^. Elevated expression of CXCR4 was also observed on pulmonary neutrophils from severe COVID-19 cases, suggesting it as a potential target for constraining ARDS induced by SARS-CoV-2 infection ^47^. It has also recently been observed to be elevated on pulmonary immune cells in the context of post-acute sequelae following SARS-CoV-2 infection of mice ^48^. As we found elevated expression of CXCR4 not only on SARS-CoV-2-specific but also total CD4+ T cells in the context of LC, targeting of this receptor as well as other chemokine receptors may be useful to limit immune cell infiltration into the lung, which may persist in an elevated state of inflammation in individuals with LC. Intriguingly, targeting of chemokine-mediated signaling has also been recently suggested as a therapy for LC based on observations that auto-antibodies against a variety of chemokines associate with protection from LC ^29^.

Another intriguing observation we made about the SARS-CoV-2-specific CD4+ T cells from individuals with LC is their production of IL6 in response to spike peptide stimulation. Although this was observed in only a small minority of individuals with LC, it suggests that a highly inflammatory response directed against the virus, persisting for at least 8 months post-infection, could be a driver of the sequelae. Although IL6 is not typically produced by lymphocytes, such atypical production of IL6 by T cells has been observed in the context of severe COVID-19 ^34^ and by B cells in the context of HIV pathogenesis ^49^. Interestingly, elevated IL6 levels have been associated with pulmonary, cardiac, and neurological LC ^22, 26, 50, 51^ and IL6 production induced by broad-spectrum mitogen PMA was found to be elevated in individuals with LC ^20^. These data together bolster the notion of targeting IL6 as a potential LC therapeutic strategy.

Most striking from our study was the finding that while fully recovered individuals exhibited coordinated humoral and cellular immune responses to SARS-CoV-2, this coordination was lost in the LC group. This finding is consistent with observations that about half of individuals with LC with no detectable SARS-CoV-2 antibodies have detectable SARS-CoV-2-specific T cell responses ^52^. That improper crosstalk between T and B cells may be involved in the etiology of LC is also supported by our RNAseq data, which showed that a cluster of genes including both immunoglobulin synthesis and T cell function were co-upregulated in those without LC, but not in individuals with LC. The downregulation of immunoglobulin-related genes in the context of LC has previously been reported and shown to be independent of spike antibody levels ^40^, which is in line with our finding higher levels of spike antibodies in our individuals with LC. How the humoral response becomes divorced from the cellular response is unclear, and could potentially involve a mis-alignment between IL4 and IL5 production by Th2 cells which emerged from our Olink analysis. Potential upstream initiators leading to the mis-coordination include a long-lived SARS-CoV-2 reservoir, reactivation of viral co-infections, or autoimmune responses.

Finally, our datasets taken together point to not only a dysregulated but also a highly pro-inflammatory signature in LC, consistent with prior data suggesting elevated and persistent inflammation in LC ^22, 23, 31, 50, 51^. Of particular interest was the elevation of the SGALS9 gene product in LC. LGALS9 encodes for Galectin 9, which has previously been shown to be upregulated during acute COVID-19 and may be a contributing factor in cytokine release and subsequent disease severity ^53, 54, 55^. The high inflammatory state observed in our individuals with LC may be in part driven by immune dysregulation, which could initiate from improper cross-talk between T and B cells as discussed above, or potentially faulty regulatory mechanisms as supported by our observation that the individuals with LC with the highest frequencies of exhausted SARS-CoV-2-specific CD8+ T cells were those that had the lowest frequencies SARS-CoV-2-specific CD4+ Treg cells. Non-immune mechanisms may also be at play, as supported by our findings that genes involved in olfactory sensing and heme synthesis were also upregulated in those with LC. The findings of increased heme synthesis were interesting in light of the fact that higher expression of genes involved in heme biosynthesis (among T cells, B cells, and monocytes) are observed during acute COVID-19 ^56, 57^, and that SARS-CoV-2 can bind hemoglobin and dysregulate heme metabolism ^58, 59, 60^. It is also possible that increased heme synthesis may reflect fibrin amyloid microclot formation that has been observed in individuals with LC ^10, 61^, and is of interest in light of our finding increased platelet numbers in the LC donors we analyzed by scRNAseq. These microclots appear to be resistant to fibrinolizes and may trap potential circulating biomarkers of the coagulopathy ^62, 63^. As a result, heme synthesis may play a useful role in determining the extent of microclot formation. Further studies of iron metabolism and red blood cell function, and their relationships to coagulopathy in the setting of LC, are warranted.

Our study has several limitations. The analysis cohort included only 43 participants due to our strict definitions of LC and complete recovery as detailed above. However, such rigor with which the participants were characterized, and their consistency in meeting the case definition longitudinally, mitigates the limitations of our small sample size, and we believe is a major strength of our study. We also note that the parent cohort is a convenience sample and certainly not representative of all individuals with a history of SARS-CoV-2 infection, although it did reflect the characteristics of the pandemic in our geographic region. Some findings we report were driven by small subsets of LC patients, which is consistent with the notion of LC being a very heterogeneous disease, and requires validation in larger cohorts. A second limitation was our focus on blood specimens, when the source of immune dysregulation, including SARS-CoV-2 persistence, likely originates from tissues, and changes in subset frequencies in blood could reflect migration into tissues. The infrastructure supporting LIINC has the ability for non-invasive tissue sampling via gut biopsies and fine needle aspirates ^64, 65^, and future studies will take advantage of these capabilities to better understand the tissue-based mechanisms underlying the immune dysregulation that manifests in LC. Finally, we note that our study is for the most part descriptive, but as for any such new and poorly-understood disease for which currently there is no good animal model, such in-depth “omics”-based characterization of a well-annotated cohort is the critical first step for better understanding the condition’s etiology and mechanistic underpinnings.

Overall, we found using multiple analytical approaches in a carefully selected cohort of individuals with consistent post-COVID symptom trajectories that LC is associated with dysregulation between humoral and cellular immunity. While LC exhibits both clinical and biological complexity, this work contributes to a growing understanding of the potential pathophysiological contributors and suggests several mechanisms warranting further exploration and/or disruption in future therapeutic trials.

## Methods

### Study participants

Participants were volunteers in the University of California, San Francisco (UCSF)-based Long-term Impact of Infection with Novel Coronavirus (LIINC) cohort (www.liincstudy.org; NCT04362150). Details of cohort recruitment, enrollment, and measurement procedures have been described in detail previously ^37^. Briefly, LIINC is a prospective observational study enrolling individuals with prior nucleic acid-confirmed SARS-CoV-2 infection in the San Francisco Bay Area, regardless of the presence or absence of post-acute symptoms. At each study visit, participants underwent an interviewer-administered assessment of 32 physical symptoms that were newly developed or had worsened since COVID-19 diagnosis, as well as assessment of mental health and quality of life. Pre-existing and unchanged symptoms were not considered to be attributable to COVID-19. In addition, detailed data regarding medical history, COVID-19 history, SARS-CoV-2 vaccination, and SARS-CoV-2 reinfection were collected. The majority of participants in LIINC did not require hospitalization for COVID-19 and were treated with symptomatic management or observation during the acute phase of infection ^37^. Consistent with broader epidemiologic trends ^66^, individuals with LC in LIINC are more likely to be female and have higher body mass index (BMI). Other pre-existing medical comorbidities are relatively uncommon in the LIINC, although we generally observe a higher rate of hypertension among those with LC compared to those who fully recovered ^23^. Two participants enrolled in LIINC had biospecimens collected previously via the UCSF COVID-19 Host Immune Response Pathogenesis (CHIRP) study, which utilizes identical procedures for ascertainment of clinical history as the LIINC study ^35^.

Because of challenges in the measurement of LC as outlined in prior work from the LIINC cohort, including within-participant symptom variability as well as the fact that some individuals with LC demonstrate symptomatic improvement and resolution of symptoms over time ^37^, we selected for this analysis participants who consistently met a case definition for LC based on the presence or absence of at least one symptom attributable to COVID-19 for the 8-month period following SARS-CoV-2 infection (Fig. S1A). The LC group (n=27) included individuals who consistently reported at least 1 COVID-19 attributed symptom during the entire study period, while the non-LC group (n=16) included individuals who consistently reported no COVID-19 attributed symptoms during the entire study period. Because of the potential effects of SARS-CoV-2 vaccination on clinical symptoms of LC as well as the immunologic measurements conducted in this study, we restricted the participants to those who provided a post-COVID blood sample prior to having ever received a SARS-CoV-2 vaccination. Blood samples were collected between September 16, 2020, and April 6, 2021.

### Biospecimen Collection

At each visit, whole blood was collected in EDTA tubes followed by density gradient separation and isolation of PBMCs and plasma, as described previously ^31^. Serum was obtained concomitantly from serum-separator tubes. Serum and plasma were stored at -80°C and PBMCs were cryopreserved and stored in liquid nitrogen.

### Antibody Assays

Antibody responses against SARS-CoV-2 spike RBD were measured on sera using the Pylon COVID-19 total antibody assay (ET Health). The assay’s lower limit of detection was 10 relative fluorescence units (RFUs).

### SARS-CoV-2 peptide pool

Peptides used for T cell stimulation comprised a mix of overlapping 15-mers spanning the entire SARS-CoV-2 spike protein (PM-WCPV-S-1, purchased from JPT), and peptides corresponding to CD8+ T cell epitopes identified by T-scan ^67^ which were synthesized in-house (Table S6). The final concentration of 15-mer peptides was 300 nM and the final concentration of T-scan peptides was 450 nM.

### CyTOF antibody conjugation

CyTOF antibody conjugation was performed using the Maxpar® X8 Antibody Labeling Kit (Standard BioTools) according to manufacturer’s instructions with some modifications. Briefly, 0.1 mg Maxpar Polymer Reagent was first dissolved in 95 µl L-buffer, and then 5 µl lanthanide solution was added into the polymer. Following a 60-min incubation at room temperature (RT), the polymer mixture was transferred into a 3-kD Amicon^TM^ Ultra tube (Fisher) and centrifuged at 12,000 g for 25 min at RT. The retentate was then washed with 400 µl C-buffer, centrifuged again, and flowthrough was discarded. Next, 100 µg antibody was mixed with 200 µl R-buffer and transferred to a 50-kD Amicon^TM^ Ultra tube (Fisher), and then centrifuged at 12,000 g for 10 min at RT. Next, 100 µl TCEP (Pierce) diluted in R-buffer (4mM final concentration) was added to the buffer-exchanged antibody retentate on the column, followed by vortexing and then a 30-min incubation at 37°C. We then added 300 µl C-buffer to the antibody mix, centrifuged the mixture at 12,000 g for 10 min at RT, washed with 400 µl C-buffer, then centrifuged again. Next, 200 µl C-buffer was added to resuspend the polymer (in the 3-kD Amicon^TM^ Ultra tube), which was then transferred to the antibody column and placed in a new collection tube. Following a 60-min incubation at 37°C, 300 µl W-buffer was added into the antibody conjugation mix. The column was then centrifuged at 12,000 g for 10 min at RT and washed three times with 400 µl W-buffer through sequential centrifugation and discarding of flowthrough. Finally, 50 µl W-buffer was added to the column, which was centrifuged for 2 min at 1,000 g. The flowthrough in the bottom collection tube was then collected to recover the conjugated CyTOF antibody. The last collection step was repeated a total of two times and eluted material combined.

### Sample preparation for CyTOF

Sample preparation was performed similarly to methods previously described ^32, 33, 34, 35^ with some modifications. Briefly, PBMCs were isolated from fresh blood draws from the LIINC participants and cryopreserved. Upon revival, cells were rested overnight to allow for antigen recovery ^68^, and then divided equally into two aliquots. To the first aliquot, we added 3 µg/ml brefeldin A (BFA, to enable intracellular cytokine detection), the co-stimulation agonists anti-CD28 (2 µg/ml, BD Biosciences) and anti-CD49d (1 µg/ml, BD Biosciences), and the SARS-CoV-2 peptide pool prepared as described above. To the second aliquot, we added only 1% DMSO (Sigma) and 3 µg/ml BFA. Cells from both treatments were incubated at 37°C for 6 h. Thereafter, cells were treated with cisplatin (Sigma) as a live/dead distinguisher and then fixed in paraformaldehyde (PFA, Electron Microscopy Science) using methods similar to those recently implemented ^32, 33, 34, 35^. Briefly, 6 million cells were resuspended in 4 ml of PBS (Rockland) containing 2 mM EDTA (Corning) and 25 µM cisplatin (Sigma), and incubated for 60 seconds. Cells were then washed twice in 1 ml CyFACS (containing 0.1% Bovine Serum Albumin (Sigma) and 0.1% Sodium Azide (Sigma) in PBS) and fixed for 10 mins at room temperature in 1.2 ml 2% PFA diluted in CyFACS. Cells were then washed twice with 1 ml CyFACS, resuspended in 100 µl 10% DMSO (Sigma) diluted in CyFACS, and frozen at -80℃ until CyTOF staining. PBMCs from a healthy donor were also subjected to the same cisplatin/fixation protocol and then aliquoted, and served as bridge samples for batch correction.

CyTOF staining was performed similar to methods recently described, where we made precautions to avoid EDTA during all CyTOF staining steps as EDTA can interfere with chemokine receptor staining ^32, 33, 34, 35^. Cisplatin-treated and PFA-fixed specimens were barcoded using Cell-ID™ 20-Plex Pd Barcoding Kit (Standard BioTools) according to manufacturer’s instructions. After barcoding, cells from up to 20 different samples were then combined into a single sample, at a concentration of 6 million cells per sample in 100 µl. The cells were then blocked at 4℃ for 15 mins in 200 µl CyFACS buffer containing 3 µl rat serum (Invitrogen), 3 µl mouse serum (Invitrogen), and 0.6 µl human serum (Sigma). After two washes with CyFACS, cells were subjected to surface antibody staining by resuspending the cells in 100 µl of freshly-prepared cell surface antibody mix (Table S2). Staining was allowed to proceed for 45 mins at 4℃. After 3 washes with CyFACS, cells were fixed overnight in 100 µl 2% PFA diluted in PBS. The next day, cells were washed with CyFACS and permeabilized by resuspension in 200 µl Foxp3 fix/perm buffer (eBioscience), and incubated at 4℃ for 30 mins. The cells were pelleted and then blocked at 4℃ for 15 mins by addition of a pre-mixed solution of 15 µl mouse serum, 15 µl rat serum, and 70 µl permeabilization Buffer (eBioscience). The cells were then washed in 800 µl permeabilization buffer (eBioscience), and incubated at 4℃ for 45 mins in 100 µl freshly-prepared intracellular staining antibody mix (Table S2). After another two washes with CyFACS, cells were stained with 250 nM Intercalator-IR (Standard BioTools) at room temperature for 20 mins. Finally, after two additional washes with CyFACS, the cells were then fixed with 1 ml 2% PFA diluted in CyFACS.

### CyTOF data acquisition

The PFA-fixed samples were washed twice with CAS buffer (Standard BioTools) and then spiked with 10% (v/v) EQ™ Four Element Calibration Beads (Standard BioTools) diluted in CAS buffer, before loading onto a Helios CyTOF instrument (UCSF Parnassus Flow Core). A running speed of 200 to 400 events per second was maintained during sample collection, and the loading voltage was controlled between 4 and 5 to minimize clogging. Data were normalized to EQ beads by CyTOF software provided by Standard BioTools to batch-correct for instrument sensitivity during sample collection. Data matrices were exported as flow cytometry standard (fcs) files for data analyses as described below.

### T cell CyTOF data analyses

#### Data preprocessing

EQ bead-normalized CyTOF datasets were concatenated, de-barcoded, and normalized using CyTOF software provided by Standard BioTools (version 6.7) according to manufacturer’s instructions. Following arcsinh transformation of the data as described ^69^, cells were then analyzed by FlowJo (version 10.8.1, BD Biosciences). Intact (Ir191+Ir193+), live (Pt195-), singlet events were identified as described in Fig. S2A, B. Those events were then gated on T cells (CD3+) followed by sub-gating on CD4+ T cells and CD8+ T cells (Fig. S2A, B).

#### CyTOF antibody validation

Our CyTOF panel (Table S2) was designed to incorporate markers commonly used to define T cell subsets (e.g., CD4, CD8α, CD45RA, CD45RO, CCR7, CD27, CD95, PD1, CXCR5, CD25, CD127, Foxp3), activation states (e.g., CD38, ICOS, CD25, Ox40, Ki67, CD69, HLADR), exhaustion states (e.g., PD1, CTLA4, TIGIT), homing properties (e.g., CCR6, CCR5, CXCR4, CD62L, CD29, CCR7, CXCR5), and effector function (e.g., IFNγ, TNFα, IL2, IL4, IL6, IL17, MIP1β, Granzyme B, perforin). Some of these markers were previously associated with immune responses to COVID-19 ^34^. CyTOF antibodies were validated using methods previously described ^32, 33, 34, 35, 65, 69^, by demonstrating expected expression patterns among and between immune subsets. Antigen expression patterns detected using the CyTOF antibodies used in this study were similar to those previously observed ^32, 33, 34, 35, 65, 69^. Fig. S18 illustrates some examples of CyTOF antibody validation implemented for this study. CyTOF analysis of human lymphoid aggregate cultures generated from fresh tonsils were performed as described ^69^. The observed expression patterns among tonsillar T and B cells (Fig. S18A) were similar to those previously observed and validated ^69^. To validate detection of cytokines and other effector molecules, we stimulated PBMCs with 16 nM PMA (Sigma) and 1 μM ionomycin (Sigma), or 1 μg/ml LPS (eBioscience), for 4 hours in the presence of 3 μg/ml brefeldin A solution (eBioscience), and then cisplatin-treated and PFA-fixed the cells as described above. Cells from the two stimulation conditions were then combined. We observed the expected induction of cytokines or cytolytic markers among these cells (Fig. S18B) ^32, 33, 34, 35^. We observed preferential expression of Treg lineage marker Foxp3 among our gated population of Treg cells (Fig. S18C), and preferential expression of CD30 and Ki67 in memory as compared to naïve CD4+ T cells (Fig. S18D) as expected, thereby validating these lowly-expressed antigens. An example of peripheral Tfh gate establishment is depicted in Fig. S18E, where we gated on memory CD4+ T cells expressing high levels of CXCR5 and PD1.

#### Identification of SARS-CoV-2-specific T cells

For identification and definition of SARS-CoV-2-specific T cells, we compared unstimulated specimens to their peptide-stimulated counterparts. Effector cytokines (IFNγ, TNFα, IL2, IL4, IL6, IL17, MIP1β), cytolytic effectors (Granzyme B and perforin) and LAMP1 were assessed for the ability to identify antigen-specific T cells at the single-cell level. Of note, in our system, as the antibody against LAMP1 was added during intracellular staining and not at the time of peptide stimulation, it was not used as a degranulation marker, but rather only to quantitate intracellular LAMP1 expression. The following criteria were established to identify effector molecules appropriate for identifying SARS-CoV-2-specific T cells: 1) counts of positive cells in unstimulated sample (not receiving peptide) was less than 5 events, or the frequency of positive cells was lower than 0.1%; 2) counts of positive cells in the peptide-stimulated sample was not less than 5, or the frequency was higher than 0.1%; 3) differences in frequencies of positive cells between unstimulated and peptide-stimulated samples cells was not less than 0.01%, 4) the fold-change in frequencies of positive cells between unstimulated and peptide-stimulated samples cells was greater than 10; and 5) the aforementioned 4 criteria could identify SARS-CoV-2-specific T cells among >50% of participants. Effectors which fulfilled all five criteria for CD4+ T cells were IFNγ, TNFα, and IL2, and those which fulfilled all five criteria for CD8+ T cells were IFNγ, TNFα, and MIP1β. For a sub-analysis to identify responding cells that may only exist in a small subset of individuals, we removed criteria #5 and reduced the positive cell counts to number 3 within criteria #1 and #2. This approach allowed us to determine that SARS-CoV-2-specific CD4+ T cells producing IL6 were exclusively detected from LC (Fig. S3D). Of note, activation induced markers (AIM) such as Ox40, CD69, and CD38 were not used to identify SARS-CoV-2-specific T cells because they cannot specifically distinguish these cells from bystander T cells after a 6-hour stimulation ^34^. SARS-CoV-2-specific T cells were detected at a median of 163 cells (134 for CD4+ T cells, 29 for CD8+ T cells), and a mean of 221.7 cells (185.2 for CD4+ T cells, 36.4 for CD8+ T cells), per participant. SARS-CoV-2-specific T cells, once identified, were analyzed by Boolean gating ^70^ and exported for further analyses.

#### SPICE

SPICE analyses were performed similar to previously described methods ^71^. Briefly, CD4+ and CD8+ T cells were subjected to manual gating based on expression of cytokines used to define SARS-CoV-2-specific T cells (IFNγ, TNFα, IL2, and MIP1β, see above) using operations of Boolean logic. The dataset matrix generated from Boolean gating was then inputted into SPICE software (version 6.1) for polyfunctional analysis. The parameters for running the dataset were: iterations for permutation test = 10,000, and highlight values = 0.05. The parameters for the query structure were set as follows: values = frequency of single cytokine positive cells in total CD4+/CD8+ T cells (generated directly from FlowJo); category = IFNγ, TNFα, IL2, and MIP1β; overlay = patient type (LC vs. Non-LC); group = all other variables in the data matrix (including sex, PID, cell type, hospitalization status, and batch). All other parameters for SPICE analyses were kept as default.

#### T cell subsetting

Manual gating was performed using R (version 4.1.3). Briefly, arcsinh-transformed data corresponding to total or SARS-CoV-2-specific T cells were plotted as 2D plots using the CytoExploreR package. Statistical data were then exported for further analyses. Visualization of datasets by tSNE were performed using R (version 4.1.3), using methods similar to those previously described ^32, 33, 34, 35^. Briefly, CytoExploreR and tidyr packages were used to load the data. tSNE was performed using Rtsne and RColorBrewer packages on arcsinh-transformed markers. Total CD4+/CD8+ T cells were downsampled to n = 8000 (maximal cell number for individual samples) before tSNE analysis. The parameters for tSNE were set as follows: iteration = 1000, perplexity = 30, and theta= 0.5.

#### T cell clustering analysis

FCS files corresponding to total and SARS-CoV-2 specific CD4+ and CD8+ T cells were imported in R for data transformation. Packages of flowcore, expss, class, and openxlsx were loaded in R for training FCS files. Arcsinh-transformed data were then exported as csv files for clustering analyses. Biological (LC status, biological sex, hospitalization status) and technical (batch/run of processing) variables were visualized using the DimPlot function in the Seurat package ^72^. As batch effects associated with the processing run were evident, batch correction was performed across the 6 batches using the harmony ^73^ batch correction function RunHarmony, applied to the marker levels in cells. The optimal clustering resolution parameters were determined using Random Forests ^74^ and a silhouette score-based assessment of clustering validity and subject-wise cross-validation. This procedure is described in greater detail in George et al. ^64^. A generalized linear mixed model (GLMM) (implemented in the lme4 ^75^ package in R with family argument set to the binomial probability distribution) was used to estimate the association between cluster membership and LC status and the biological sex of the participant, with the participant modeled as a random effect. For each given participant, cluster membership of cells is encoded as a pair of numbers representing the number of cells in the cluster and the number of cells not in the cluster. Clusters having fewer than 3 cells were discarded. The sex-specific log odds ratio of cluster membership association with LC status was estimated using the emmeans ^76^ R package using the GLMM model fit. The estimated log odds ratio represents the change (due to LC status) in the average over all participants of a given sex in the log odds of cluster membership. The two-sided p-values corresponding to the null hypothesis of an odds ratio value of 1 was computed based on a Z-statistic in the GLMM model fit. These p-values were adjusted for multiple testing using the Benjamini-Hochberg method.

### FACS

Validation experiments using flow cytometry (FACS) was performed on total PBMCs from n=40 individuals in our cohort (25 LC, 15 non-LC). Sample preparation and cell revival were performed as described above for CyTOF preparation, and analyzed at baseline, or following SARS-CoV-2 peptide stimulation. For staining, cells were first treated with Zombie UV^TM^ or Zombie NIR^TM^ (BioLegend) as viability indicators, and then blocked with Human TruStain FcX^TM^ (BioLegend). FACS antibodies that were used in this study are shown in Table S7. For intracellular staining, cells were fixed with 2% PFA and incubated with Foxp3 fix/perm buffer (eBioscience) per manufacturer’s instructions, following completion of surface antibody staining. The BD Horizon^TM^ Brilliant stain buffer (BD Biosciences) was used according to manufacturer’s instructions when staining with antibodies conjugated to brilliant dyes. All cells were fixed in 2% PFA prior to analysis on a Fortessa X-20 (BD Biosciences). UltraComp eBeads™ Compensation Beads (Invitrogen) served as a single fluorescence dye control, and ArC™ Amine Reactive Compensation Beads (Invitrogen) served as a loading control for live/dead cell staining. FCS files were exported into FlowJo (BD, version 10.9.0) for further analyses. FACS data were arcsinh-scaled prior to analyses.

### RNAseq

RNAseq was performed on total PBMCs from n=36 individuals in our cohort using the AllPrep kit as per manufacturer’s instructions (Qiagen). RNA libraries, next generation Illumina sequencing, quality control analysis, trimming, and alignment to the human genome (hg19) were performed by Genewiz (Azenta Inc.). Briefly, following oligo dT enrichment, fragmentation and random priming, cDNA syntheses was completed. End repair, 5’ phosphorylation and dA-tailing were performed, followed by adaptor ligation, PCR enrichment, and sequencing on an Illumina HiSeq platform using PE150 (paired-end sequencing, 150 bp for reads 1 and reads 2). Raw reads (480 Gb in total) were trimmed using Trimmomatic (version 0.36) to remove adapter sequences and poor-quality reads. Trimmed reads were then mapped to Homo sapiens GRCh37 using star aligner (version 2.5.2b) ^77^. Log2 fold-changes were calculated between those with or without any LC symptoms. Two-sided P values corresponding to a null hypothesis of fold-change of 1 were calculated using DESeq2’s ^78^ Wald test, and were adjusted controlling for multiple testing using false discovery rates. Genes with an adjusted p-value < 0.05 and absolute log2 fold-change > 1 were considered as significantly differentially expressed genes (DEGs). Clustered heatmaps of DEG were constructed with groups of genes (rows) defined using the k-means algorithm to cluster genes into k clusters based on their similarity. K = 4 was determined using the HOPACH (Hierarchical Ordered Partitioning and Collapsing Hybrid) algorithm ^79^, which recursively partitions a hierarchical tree while ordering and collapsing clusters at each level to identify the level of the tree with maximally homogeneous clusters.

### scRNAseq

scRNAseq library preparation was performed on total PBMCs from n=12 individuals (4 non-LCs and 8 LCs) in our cohort using the Chromium Next GEM Single Cell 5’ Reagent Kits v2 (10x Genomics) according to manufacturer’s instruction. This was then followed by sequencing on the Illumina NovaSeq 6000 S4 300 platform (UCSF Center for Advanced Technology). All 12 samples were sequenced at a mean of >50K reads per cell (minimum 51K, maximum 120K, median 83K). A total of 1.3Tb sequencing data was collected. A median of 7888 cells was analyzed per donor (minimum 4189, maximum 9511). Demultiplexed fastq files were aligned to the human reference genome (GRCh38) using the 10x Genomics Cell Ranger v7.1.0 count pipeline ^80^, as described in the Cell Ranger documentation. The include-introns flag for the count pipeline was set to true to count reads mapping to intronic regions.

The filtered count matrices generated by the Cell Ranger count pipeline were processed using the R package for using Seurat ^72^. Each sample was pre-processed as a Seurat object, and the top 1% of cells per sample with highest numbers of unique genes, cells with <=200 unique genes, and cells >=10% mitochondrial genes were filtered out for each sample. The 12 samples were then merged into a single Seurat object, and normalization and variance stabilization was performed using sctransform86 with the “glmGamPoi” method ^81^ for initial parameter estimation.

Graph-based clustering was performed using the Seurat ^72^ functions FindNeighbors and FindClusters. First, the cells were embedded in a k-nearest neighbor (KNN) graph (with k=20) based on the Euclidean distance in the PCA space. The edge weights between two cells were further modified using Jaccard similarity. Next, clustering was performed using the Louvain algorithm ^82^ implementation in the FindClusters Seurat function. Clustering with 15 Principal Components (PCs) (determined based on location of elbow in the plot of variance explained by each of the top 25 PCs) and 0.1 resolution (determined using the resolution optimization method described above for CyTOF data clustering analyses) resulted in 11 distinct biologically relevant clusters (Clusters 0-11), which were used for further analyses.

Data visualization using Seurat in UMAP space for the 12 samples revealed no apparent batch effects due to age, sex (all were female), race, or hospitalization status of the subjects. The marker genes for each cluster were identified using the FindAllMarkers Seurat function. This algorithm uses the Wilcoxon Rank Sum test to iteratively identify DEGs in a cluster against all the other clusters. Marker genes were filtered to keep only positively expressed genes, detected in at least 25% of the cells, and with at least 0.5 log2 fold change. Cluster annotation was performed according to subset definitions previously established ^83, 84, 85^. Classification markers included CD19, MS4A1, and CD79A for B cells; CD3D, CD3E, CD5, and IL7R for CD4+ T cells; CD3D, CD3E, CD8A, CD8B, and GZMK (CTL subset) for CD8+ T cells; CD14, CD68, CYBB, S100A8, S100A9, S100A12, and LYZ for monocytes; CSF2RA, LYZ, CXCL8, and CD63 for granulocytes; and PF4, CAVIN2, PPBP, GNG11, and CLU for platelets.

The Counts-Per-Million (CPM) (pseudo-bulked) reads for ALAS2 and OR7D2 were assessed in each of the subjects using edgeR ^86^, and associations with group status were using the two sample Welch t-test, followed by multiple correction testing using the Holm ^87^ procedure. For establishing associations between clusters and group status, GLMM implemented in the lme4 R package was used separately for each cluster of cells. The model was performed with the family argument set to the binomial probability distribution, and with the ‘nAGQ’ parameter set to 10 corresponding to the number of points per axis for evaluating the adaptive Gauss-Hermite approximation for the log-likelihood estimation. Cluster membership of cells by sample was modeled as a response variable by a 2-dimensional vector representing the numbers of cells from a given sample belonging or not to the cluster under consideration. The corresponding sample from which the cell was derived was the random effect variable, and the group (non-LC, LC, ORD72^high^ LC, or ALAS2^high^ LC) was considered the fixed variable. The log odds ratio for all pairwise comparisons were estimated using the model fits provided to the emmeans function in the emmeans R package ^76^. The resulting p-values for the estimated log odds ratio and clusters were adjusted for multiple testing using the Benjamini-Hochberg method^88^. For associations of gene expression with group status within each cluster, the raw gene counts per cell were loaded as a SingleCellExperiment object. Cells from clusters 9 and 10 were not included in this analysis as the median number of cells across samples was less than 20 for each of these clusters. The aggregateData function in the muscat bioconductor package ^89^ was used to pseudo-bulk the gene read counts across cells for each cluster group. Genes with raw counts less than 10 in more than 8 samples were removed from the analyses. The pbDS function implementing the statistical methods in the edgeR package ^86^ was used to assess associations of gene expression with group identity in each of the clusters.

### Olink

We performed the Olink EXPLORE 384 inflammation Protein Extension Assay (PEA) from plasma from n=40 individuals in our cohort to characterize 384 unique plasma proteins associated with inflammation and immune signaling. PEA involves dual-recognition of two matched antibodies labelled with unique DNA oligonucleotides that simultaneously bind to specific target proteins. The simultaneous antibody binding leads to hybridization of unique DNA oligonucleotides that serve as templates for polymerase-dependent extension (DNA barcoding) followed by PCR amplification and NovaSeq (Illumina) DNA sequencing as published ^90, 91, 92, 93, 94^. A similar analysis pipeline was applied to the protein biomarkers as described for the gene expression data as above.

### Data visualization for RNAseq and Olink

To generate heatmaps, the R package HOPACH ^79^ was used to find the best cluster number. Gene expression values were log-transformed and centered using the average expression value for each gene. Genes were clustered by running the Kmeans algorithm using the best cluster number K found, and the results were plotted using the pheatmap package ^95^. For gene network analyses, the STRING interaction database was used to reconstruct gene networks using stringApp ^96^ for Cytoscape ^97^. For the network, the top 50 genes or 25 proteins with the lowest p-values were selected from the RNAseq data and Olink data, respectively. Then genes were subjected to stringApp with an interaction score cutoff = 0.5, and the number of maximum additional indirect interactors cutoff = 10. The analysis integrated STRING data with our gene inputs, resulting in a network of 24 nodes and 100 edges in for the RNAseq data, and a network of 26 nodes and 165 edges for the Olink data. In each network, a node corresponds to a gene, an edge represents the functional relevance between a pair of genes, with the thickness of each edge reflecting the confidence level. Node color indicates the degree of log2 fold-change and the difference between protein expression values for the RNAseq and Olink data, respectively.

## ACKNOWLEDGEMENTS

This work was supported by the Van Auken Private Foundation, David Henke, and Pamela and Edward Taft; philanthropic funds donated to Gladstone Institutes by The Roddenberry Foundation and individual donors devoted to COVID-19 research; the Program for Breakthrough Biomedical Research, which is partly funded by the Sandler Foundation; and Awards #2164 and #2208 from Fast Grants, a part of Emergent Ventures at the Mercatus Center, George Mason University. We acknowledge the NIH DRC Center Grant P30 DK063720 and the S10 1S10OD018040-01 for use of the CyTOF instrument, and the NIH S10 RR028962 and the James B. Pendleton Charitable Trust for use of the Fortessa X-20. This study was also funded by the Ministerium für Wissenschaft, Forschung und Kunst, Baden Württemberg, Germany (KNKC.031) and the Deutsche Forschungsgemeinschaft (DFG, German Research Foundation)—Projektnummer 316249678— SFB 1279. The parent cohort (LIINC) is supported by NIH/NIAID 3R01AI141003-03S1 and NIH/NIAID R01AI158013. We thank Stanley Tamaki, Vinh Nguyen, Pricsilla Sanchez and Claudia Bispo for CyTOF assistance at the Parnassus Flow Core, Jane Srivastava and Vaishanavi Saware for technical assistance in FACS, Merve Karacan and Nico Preising for technical assistance in peptide synthesis, Eliver Ghosn for guidance on annotation of cell clusters identified by scRNAseq, John Carroll for assistance on graphics, Françoise Chanut for editorial assistance, and Robin Givens for administrative assistance.

We are grateful to the study participants and their medical providers. We acknowledge current and former LIINC clinical study team members Tamara Abualhsan, Andrea Alvarez, Mireya Arreguin, Melissa Buitrago, Monika Deswal, Nicole DelCastillo, Emily Fehrman, Halle Grebe, Heather Hartig, Yanel Hernandez, Marian Kerbleski, Raushun Kirtikar, James Lombardo, Michael Luna, Lynn Ngo, Enrique Martinez Ortiz, Antonio Rodriguez, Justin Romero, Dylan Ryder, Ruth Diaz Sanchez, Matthew So, Celina Song, Viva Tai, Alex Tang, Cassandra Thanh, Fatima Ticas, Leonel Torres, Brandon Tran, Deepshika Varma, and Meghann Williams; and LIINC laboratory team members Amanda Buck, Joanna Donatelli, Jill Hakim, Nikita Iyer, Owen Janson, Brian LaFranchi, Christopher Nixon, Isaac Thomas, and Keirstinne Turcios. We thank Jessica Chen, Aidan Donovan, and Carrie Forman for assistance with data entry and review. We thank the UCSF AIDS Specimen Bank for processing specimens and maintaining the LIINC biospecimen repository. We are grateful to Elnaz Eilkhani and Monika Deswal for regulatory support. We are also grateful for the contributions of additional current and former LIINC leadership team members: Matthew Durstenfeld, Priscilla Hsue, Bryan Greenhouse, Isabelle Rodriguez-Barraquer, and Rachel Rutishauser.

## AUTHOR CONTRIBUTIONS

K.Y. designed the experiments, performed CyTOF, FACS, and scRNAseq experiments, conducted analyses, prepared figures and tables, and wrote the manuscript; M.J.P. designed the LIINC cohort, oversaw LIINC cohort procedures, interpreted data, and wrote the manuscript; X.L. developed pipelines for data analyses and performed scRNAseq; R.T. performed clustering and scRNAseq analyses; M.S. performed RNAseq and Olink analyses; J.N. prepared peptides and helped with experiments; A.A. and K.Y. prepared and analyzed CyTOF specimens, T.M. designed protocols for CyTOF analyses; R.H., K.A. and B.H. managed the LIINC cohort, recruited participants, collected clinical data, and collected biospecimens; U.A., M.L., D.V., K.A., T.D., and S.E.M. recruited LIINC participants, collected clinical data, and collected biospecimens; T.D. and S.E.M. processed specimens; L.S. and J.M. synthesized peptides; R.I. entered, cleaned, and performed quality control on LIINC data; S.L. and S.A.G. managed LIINC data and selected biospecimens; K.L.L. performed antibody assays; S.A.L. designed the CHIRP cohort and oversaw CHIRP cohort procedures; J.D.K. and J.N.M designed the LIINC cohort and interpreted LIINC clinical data; S.G.D. designed the LIINC cohort, oversaw cohort procedures, and interpreted LIINC clinical data; T.J.H. designed the LIINC cohort, oversaw cohort procedures, performed the RNAseq and Olink studies, interpreted data, prepared figures, and wrote the manuscript; N.R.R. conceived the study, performed supervision, conducted data analyses, prepared figures and tables, and wrote the manuscript. All authors have read and approved this manuscript.

## Data availability

The raw CyTOF datasets for this study corresponding to total and SARS-CoV-2-specific CD4+ and CD8+ T cells are publicly accessible through the following link: https://datadryad.org/stash/share/TE_QuY0JX23V2n2CIMO2PgsR6afIp6GGusdQ5nXVGnk. The raw bulk RNAseq and scRNAseq data from this study are deposited in the GEO (Gene Expression Omnibus) database: GSE224615 (for bulk RNAseq) and GSE235050 (for scRNAseq). All other raw datasets from this study are available from the corresponding authors upon request.

## Statistical tests

Unless otherwise indicated, permutation tests, two-tailed unpaired student’s t-tests, and Welch’s t test were used for statistical analyses. *p < 0.05, **p < 0.01, ***p < 0.001, ****p<0.0001, and n.s. non-significant. Error bars corresponded to standard deviation (SD). Graphs were plotted by GraphPad Prism (version 9.4.1). All measurements were taken from distinct samples, no samples were measured repeatedly to generate data. Where appropriate, p-values were corrected for multiple testing (across 3 pairwise comparisons) using the Holm procedure ^87^.

## Informed consent

All participants provided written informed consent and study protocols were approved by the UCSF Institutional Review Board.

---

**Fig. S1. Clinical characteristics of LC and non-LC individuals analyzed in this study. A.** Strategy of biospecimen selection for study. Individuals demonstrate variable recovery following COVID-19, with many experiencing symptom resolution within 4 months (typically <30 days) in a manner that is sustained (solid blue line). However, some experience symptoms that persist (solid red line). Symptoms can also newly develop (dotted red line) or diminish (dotted blue line) over time. Per WHO definition, symptoms persisting beyond 3 months post-infection are defined as LC ^98^. In order to minimize the effects of between-individual heterogeneity, we selected individuals demonstrating a consistent post-acute phenotype (LC or Non-LC) at two timepoints timed 4 and 8 months following initial SARS-CoV-2 infection, and performed biologic measurements on samples from the 8-month timepoint. All samples were collected and cryopreserved prior to the individuals ever receiving a SARS-CoV-2 vaccine or experiencing a clinically detected reinfection. **B.** Sex distribution of individuals from the LIINC cohort analyzed in this study. Overall, 55.8% of the participants were female; the non-LC group comprised 43.75% females and the LC group 62.96% females. **C.** The proportion of LC and non-LC individuals that were hospitalized at the time of acute COVID-19 infection. Overall, 20.9% of the participants were hospitalized: 12.5% among the non-LC individuals, and 25.9% among the individuals with LC. **D.** The number of sequelae symptoms among the individuals with LC. Number of sequelae symptoms at four (M4) and eight (M8) months are shown. *p<0.05 (paired sample t-test). **E**. Most individuals in the cohort did not exhibit comorbidities. The proportion of LC and non-LC individuals that exhibited the indicated comorbidities are indicated. **F**. Individuals with LC had significantly higher BMI than individuals without LC in the cohort. *p<0.05 (student’s t-test).

**Fig. S2. Cytokine and effector molecule expression on T cells in the absence or presence of SARS-CoV-2 peptides. A-B.** Gating strategy to identify T cell populations. Intact, live, singlet cells from a baseline (unstimulated) ***(A)*** or SARS-CoV-2 peptide-treated ***(B)*** samples were gated for T cells (CD3+) followed by sub-gating on CD4+ and CD8+ T cells as indicated. **B.** CD4+ T cells from a representative donor, in the absence or presence of SARS-CoV-2 peptides. The red box highlights the three cytokines used to define SARS-CoV-2-specific CD4+ T cells. **C.** CD8+ T cells from a representative donor, in the absence or presence of SARS-CoV-2 peptides. The red boxes highlight the three cytokines used to define SARS-CoV-2-specific CD8+ T cells.

**Fig. S3. SARS-CoV-2-specific T cell responses defined individually as those producing IFNγ, IL2, TNFα, and MIP1β do not differ between LC and non-LC individuals. A.** Shown are the proportions of SARS-CoV-2-specific CD4+ T cells as defined by cells producing IFNγ, IL2, or TNFα in response to SARS-CoV-2 peptide stimulation. n.s.: non-significant as determined by student’s t-test. **B.** Shown are the proportions of SARS-CoV-2-specific CD8+ T cells as defined by cells producing IFNγ, MIP1β, or TNFα in response to SARS-CoV-2 peptide stimulation. n.s.: non-significant as determined by student’s t-test. **C.** IL6-producing CD4+ T cells are observed after SARS-CoV-2 peptide stimulation in some donors. Shown are cells from a representative individual with LC. **D.** IL6+ SARS-CoV-2-specific CD4+ T cells are exclusively observed from participants with LC. *p<0.05 (Welch’s t-test). Results in panels *C* and *D* were pre-gated on live, singlet CD4+ T cells.

**Fig. S4. Gating strategy to define classical T cell subsets.** Shown are gating strategies to define the indicated subsets of CD4+ (**A**) and CD8+ (**B**) T cells.

**Fig. S5. Subset distribution of total and SARS-CoV-2-specific CD8+ T cells among LC and Non-LC individuals. A.** Tem frequencies trended higher among total CD8+ T cells from LC as compared to non-LC individuals (student’s t-test). **B.** Temra frequencies trended lower among SARS-CoV-2-specific CD8+ T cells from LC as compared to non-LC individuals (student’s t-test).

**Fig. S6. MSI of all CyTOF phenotyping markers among CD4+ T cells from individuals with and without LC.** Antigens are shown in the order listed in Table S2. Results are gated on live, singlet CD4+ T cells. No significant differences were observed between LC and non-LC individuals for any of the antigens (t-test with multiple correction by Sidak adjustment).

**Fig. S7. MSI of all CyTOF phenotyping markers among CD8+ T cells from individuals with and without LC.** Results are similar to that shown in Fig. S6, but gated on CD8+ T cells. No significant differences were observed between LC and non-LC individuals for any of the antigens (t-test with multiple correction by Sidak adjustment).

**Fig. S8. MSI of all CyTOF phenotyping markers among SARS-CoV-2-specific CD4+ T cells from individuals with and without LC.** Results are similar to that shown in Fig. S6, but gated on SARS-CoV-2-specific CD4+ T cells. No significant differences were observed between LC and non-LC individuals for any of the antigens (t-test with multiple correction by Sidak adjustment).

**Fig. S9. MSI of all CyTOF phenotyping markers among SARS-CoV-2-specific CD4+ T cells from individuals with and without LC.** Results are similar to that shown in Fig. S6, but gated on SARS-CoV-2-specific CD8+ T cells. No significant differences were observed between LC and non-LC individuals for any of the antigens (t-test with multiple correction by Sidak adjustment).

**Fig. S10. Activated T cells are not more abundant in individuals with LC.** The percentages of total CD4+ T cells **(A)**, total CD8+ T cells **(B)**, SARS-CoV-2-specific CD4+ T cells **(C)**, and SARS-CoV-2-specific CD8+ T cells **(D)** expressing acute activation markers CD38, HLADR, and Ki67 were not significantly different between individuals with and without LC. Shown are proportions of cells expressing each of these respective markers (first three columns), or co-expressing all three of these markers (last column). n.s. = non-significant as determined by student’s t-tests.

**Fig. S11. Sex-dimorphic CD4+ T cell cluster distribution in individuals with LC. A.** Cluster distribution among baseline CD4+ T cells as depicted by UMAP. **B.** Female individuals with LC harbor significantly lower frequencies (raw p-value = 0.03, FDR = 0.19) of cluster A1 cells relative to female non-LC individuals, and male individuals with LC harbor significantly lower frequencies (raw p-value = 0.02, FDR = 0.19) of cluster A4 cells relative to male non-LC individuals. P-values were derived from a GLMM fit (see Methods). Individual points represent the % of all cells from a given participant that belong to clusters A1 or A4. **C.** Relative to total CD4+ T cells, cluster A1 is characterized by expression of naïve cell markers (CD45RA^high^CD45RO^low^CD27^high^), and low expression of inflammatory tissue homing receptors (CD29^low^CXCR4^low^) but high expression lymph node homing receptors (CD62L^high^CCR7^high^). The activation markers HLA-DR and OX40 were also lowly expressed in cluster A1 cells. **D.** Relative to total CD4+ T cells, cluster A4 is characterized by expression of terminally differentiated effector memory T cell markers (CD45RA^low^CD45RO^high^CD27^low^CCR7^low^CD57^high^), and high expression of homing receptors for inflamed tissues (CD29^high^CXCR4^high^CCR5^high^) but low expression of lymph node homing receptors (CD62L^low^CCR7^low^). The cytolysis-associated markers perforin, granzyme, and LAMP1 were expressed at elevated levels in cluster A4 cells. ****p<0.0001 (paired t-test). For panels *C* and *D*, data are depicted as histograms showing distribution of marker expression among all participants, or as bar graphs depicting the distribution of the MSI of each marker within each individual participant. Comparisons are shown between cells belonging to the indicated cluster as compared to total CD4+ T cells.

**Fig. S12. Sex-dimorphic differential CD8+ T cell cluster distribution in individuals with LC. A.** Cluster distribution among baseline CD8+ T cells as depicted by UMAP. **B.** Female individuals with LC harbor significantly lower frequencies (raw p-value = 0.046, FDR = 0.22) of cluster B1 cells and significantly higher frequencies (p-value = 0.024, FDR = 0.22) of cluster B2 cells, relative to female non-LC individuals. P-values were derived from a GLMM fit (see Methods). Individual points represent the % of all cells from a given participant that belong to clusters B1 or B2. **C.** Relative to total CD8+ T cells, cluster B1 is characterized by expression of naïve cell markers (CD45RA^high^CD45RO^low^CD27^high^), and low expression of tissue homing receptors (CD29^low^CXCR4^low^) but high expression lymph node homing receptors (CD62L^high^CCR7^high^). The activation markers HLA-DR and OX40 were also lowly expressed in cluster B1 cells. **D.** Relative to total CD8+ T cells, cluster B2 is characterized by expression of terminally differentiated effector memory T cell markers (CD45RA^low^CD45RO^high^CD27^low^CCR7^low^CD57^high^), and high expression of homing receptors for inflamed tissues (CD29^high^CXCR4^high^CCR5^high^) but low expression of lymph node homing receptors (CD62L^low^CCR7^low^). The cytolysis markers perforin, granzyme, and LAMP1 were expressed at elevated levels in cluster B2 cells. ****p<0.0001 (paired t-test). For panels *C* and *D*, data are depicted as histograms showing distribution of marker expression among all participants, or as bar graphs depicting the distribution of the MSI of each marker within each individual participant. Comparisons are shown between cells belonging to indicated cluster as compared to total CD8+ T cells.

**Fig. S13. FACS validation that CD4+ T cells from individuals with LC preferentially express CXCR4, CXCR5, and CCR6. A.** Expression levels of CXCR4, CXCR5, and CCR6 as assessed by FACS significantly correlates with those as assessed by CyTOF. FACS data were reported as mean fluorescence intensity (MFI), while CyTOF data were reported as mean signal intensity (MSI). CyTOF data were arcsinh-transformed, while FACS data were arcsinh-scaled. Correlation estimates were identified as R values (Pearson correlation coefficients). **B.** FACS gating strategy to identify memory CD4+ T cells expressing various combinations of CXCR4, CXCR5, and CCR6. **C.** CXCR4+CXCR5+, CXCR5+CCR6+, and CXCR4+CCR6+ CD4+ T cells are significantly elevated in LC as compared to non-LC individuals. *p<0.05 (student’s t-test).

**Fig. S14. FACS validation that SARS-CoV-2-specific CD8+ T cells from individuals with LC preferentially express exhaustion markers PD1 and CTLA4.** FACS analysis demonstrating that SARS-CoV-2-specific CD8+ T cells co-expressing PD1 and CTLA4 were elevated in individuals with LC, while total PD1+CTLA4+ CD8+ T cells were not. *p<0.05 (student’s t-test). Note that in these FACS experiments, SARS-CoV-2-specific CD8+ T cells were defined as those specifically inducing IFNγ and/or TNFα in response to SARS-CoV-2 peptide stimulation, while those in the CyTOF experiments (Fig. 5) were defined as those specifically inducing IFNγ, TNFα, and/or MIP1β. The difference was because MIP1β antibody exhibited background staining in FACS and could not be used to define SARS-CoV-2-specific T cells.

**Fig. S15. Global differential gene and gene product expression in participants with and without LC. A.** Heatmap of the top 50 DEGs in PBMCs in LC compared to non-LC individuals. Note data are the same as those depicted in Fig. 7B, but separated by LC vs. non-LC status. **B.** Clustered heatmap of the top 25 differentially expressed plasma proteins from Olink Proximity Extension Assay. Note data are the same as those depicted in Fig. 7F, but separated by LC vs. non-LC status.

**Fig. S16. scRNAseq analysis of reveals OR7D2 and ALAS2 expression in multiple subsets of immune cells from individuals with LC. A, B.** UMAP of cells from LC (n=8) vs. non-LC (n=4) individuals *(A)*, and among the LC individuals those classified as OR7D2^high^ (n=4) vs. ALAS2^high^ (n=4) *(B)*, revealing lack of profound global differences in gene expression between any of these groups. **C.** OR7D2 expression is highest in the OR7D2^high^ LC group, and ALAS2 expression is highest in the ALAS2^high^ LC group. **D.** OR7D2 and ALAS2 are broadly expressed by multiple PBMC subsets in individuals with LC. Shown are the mean % of cells that were positive for OR7D2 or ALAS2 reads, grouped by cellular clusters defined in Fig. 7D. Note that both OR7D2 and ALAS2 transcripts were detected in all clusters except 9 and 10. OR7D2 was most highly expressed in Cluster 3 (monocytes), while ALAS2 was most highly expressed in Cluster 8 (B cells), with Cluster 5 (CD8+ T cells) and Clusters 0/7 (CD4+ T cells) also being relatively high expressors. **E, F.** Among all clusters expressing ORD72 or ALAS2, OR7D2 was preferentially expressed in the OR7D2^high^ LC group *(E)* while ALAS2 was preferentially expressed in the ALAS2^high^ LC group *(F)*. *p<0.05, **p<0.01, as determined by two-sample Welch t-tests, followed by multiple testing correction (across 3 pairwise combinations) using the Holm procedure.

**Fig. S17. Opposite associations of IL4 levels with T cells expressing inflammatory chemokine receptors in individuals with vs. without LC.** (**A**) The percentages of CXCR4+CXCR5+ CD4+ T cells are significantly positively associated with IL4 levels in individuals with LC (p<0.05), while these parameters negatively associate in individuals without LC (p<0.05). (**B**) The percentages of CXCR5+CCR6+ CD4+ T cells are significantly positively associated with IL4 levels in individuals with LC (p<0.05), while these parameters negatively associate in individuals without LC (p<0.01). Correlation estimates were defined as R values in the figures, and significance was determined using Pearson r t-tests.

**Fig. S18. Validation of CyTOF antibodies.** (**A**) Validation of CyTOF detection of antigens through analysis of expression patterns on tonsillar T cells and B cells. Most cells from the human lymphoid aggregate cultures generated from tonsils correspond to T cells and B cells ^69^. Therefore, as demonstrated in the schematic on the bottom, visualizing CD3 on the y-axis allows differentiation of T cells (top population) from B cells (bottom population). The expression patterns of the indicated antigens on T and B cells are similar to those previously reported and validated^69^. (**B**) Validation of CyTOF detection of effector molecule expression was performed by using PBMCs stimulated with PMA/ionomycin or LPS (see Methods). As demonstrated in the schematic, visualizing CD3 on the y-axis allows differentiation of T cells (top population) from other immune subsets including monocytes and other innate immune cells. The expression patterns of the shown effectors are as expected and consistent prior studies ^32, 33, 34, 35^. (**C**) Validation of Treg gating. Compared to naïve CD4+ T cells (Tn: CD3+CD4+CD45RO-CD45RA+CCR7+CD95-), the Treg population (defined as CD3+CD4+CD45RO+CD45RA-CD25+CD127-) expressed significantly higher levels of the Treg lineage marker Foxp3, thereby validating their Treg identity. ****p<0.0001 (paired t-test). (**D**) CyTOF detection of CD30 and Ki67 were validated by demonstrating that memory CD4+ T cells (Tm: CD3+CD4+CD45RO+CD45RA-) expressed significantly higher levels of these markers as compared to naïve CD4+ T cells, as expected. ****p<0.0001 (paired t-test). (**E**) Illustration of Tfh gate on PBMC samples used for validation. Cells were pre-gated on memory CD4+ T cells.

**Table S1:**
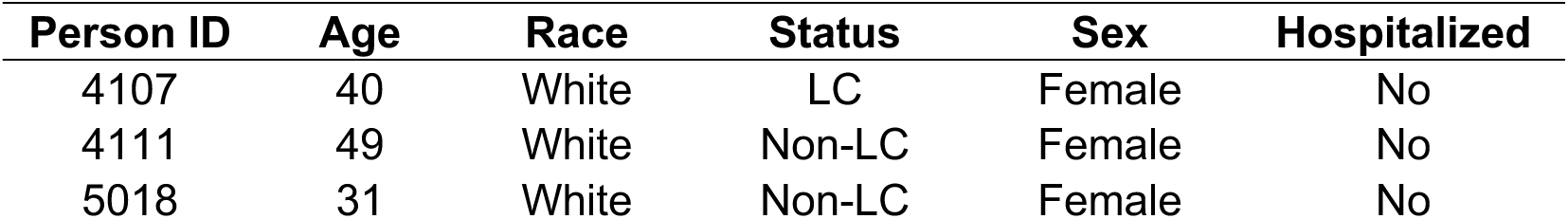
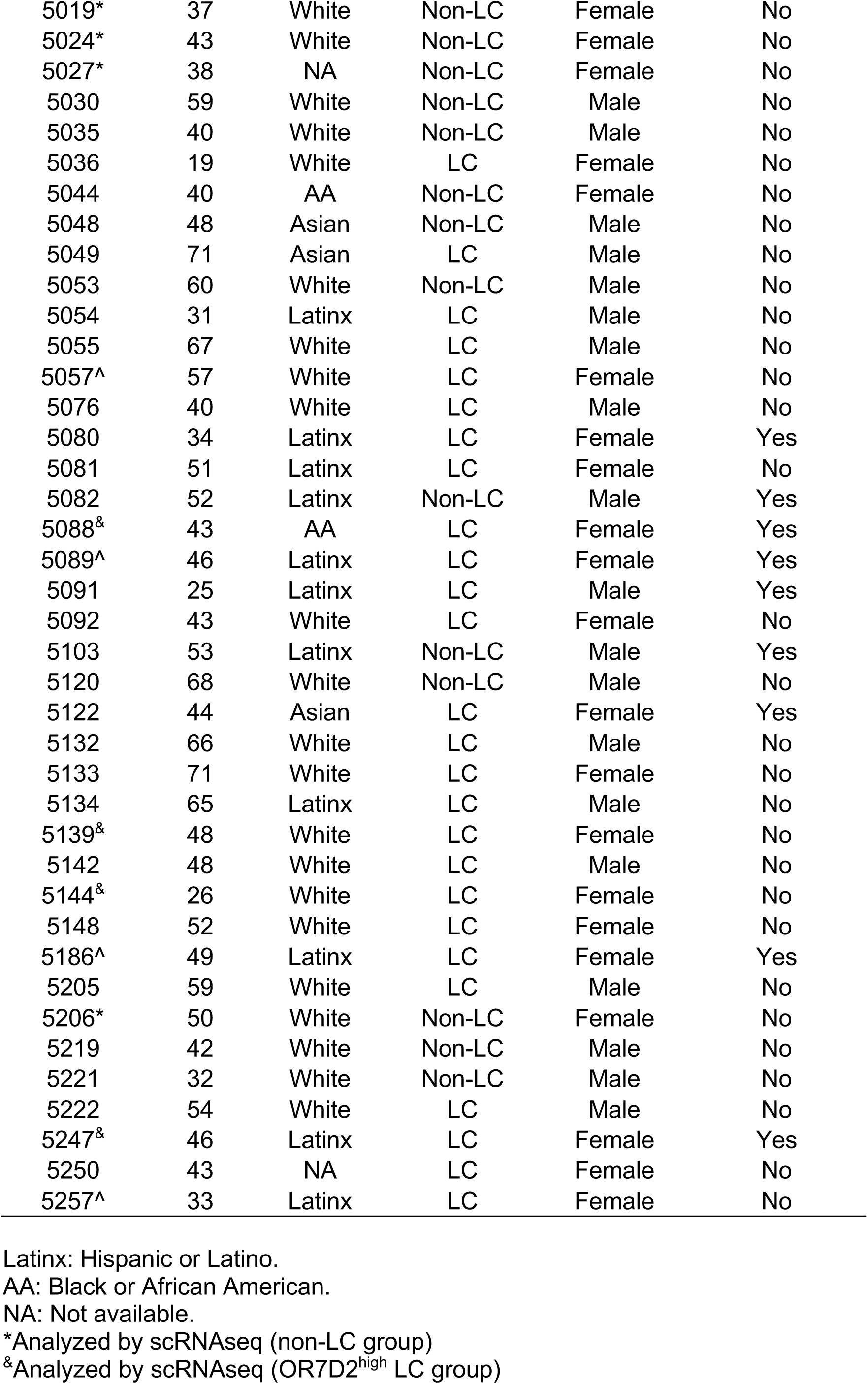
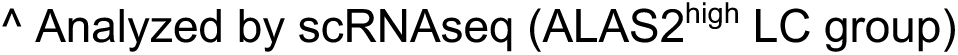
Participants from the LIINC cohort.

| Person ID | Age | Race | Status | Sex | Hospitalized |
| --- | --- | --- | --- | --- | --- |
| 4107 | 40 | White | LC | Female | No |
| 4111 | 49 | White | Non-LC | Female | No |
| 5018 | 31 | White | Non-LC | Female | No |
| 5019* | 37 | White | Non-LC | Female | No |
| 5024* | 43 | White | Non-LC | Female | No |
| 5027* | 38 | NA | Non-LC | Female | No |
| 5030 | 59 | White | Non-LC | Male | No |
| 5035 | 40 | White | Non-LC | Male | No |
| 5036 | 19 | White | LC | Female | No |
| 5044 | 40 | AA | Non-LC | Female | No |
| 5048 | 48 | Asian | Non-LC | Male | No |
| 5049 | 71 | Asian | LC | Male | No |
| 5053 | 60 | White | Non-LC | Male | No |
| 5054 | 31 | Latinx | LC | Male | No |
| 5055 | 67 | White | LC | Male | No |
| 5057^ | 57 | White | LC | Female | No |
| 5076 | 40 | White | LC | Male | No |
| 5080 | 34 | Latinx | LC | Female | Yes |
| 5081 | 51 | Latinx | LC | Female | No |
| 5082 | 52 | Latinx | Non-LC | Male | Yes |
| 5088& | 43 | AA | LC | Female | Yes |
| 5089^ | 46 | Latinx | LC | Female | Yes |
| 5091 | 25 | Latinx | LC | Male | Yes |
| 5092 | 43 | White | LC | Female | No |
| 5103 | 53 | Latinx | Non-LC | Male | Yes |
| 5120 | 68 | White | Non-LC | Male | No |
| 5122 | 44 | Asian | LC | Female | Yes |
| 5132 | 66 | White | LC | Male | No |
| 5133 | 71 | White | LC | Female | No |
| 5134 | 65 | Latinx | LC | Male | No |
| 5139& | 48 | White | LC | Female | No |
| 5142 | 48 | White | LC | Male | No |
| 5144& | 26 | White | LC | Female | No |
| 5148 | 52 | White | LC | Female | No |
| 5186^ | 49 | Latinx | LC | Female | Yes |
| 5205 | 59 | White | LC | Male | No |
| 5206* | 50 | White | Non-LC | Female | No |
| 5219 | 42 | White | Non-LC | Male | No |
| 5221 | 32 | White | Non-LC | Male | No |
| 5222 | 54 | White | LC | Male | No |
| 5247& | 46 | Latinx | LC | Female | Yes |
| 5250 | 43 | NA | LC | Female | No |
| 5257^ | 33 | Latinx | LC | Female | No |

Latinx: Hispanic or Latino.
AA: Black or African American.
NA: Not available.
\*Analyzed by scRNAseq (non-LC group)
&Analyzed by scRNAseq (OR7D2<sup>high</sup> LC group)
^ Analyzed by scRNAseq (ALAS2<sup>high</sup> LC group)

**Table S2:** CyTOF antibodies used in study.

| Antibody | Clone | Catalog | Metal | Manufacturer |
| --- | --- | --- | --- | --- |
| CD196/CCR6 | 11A9 | 3141014A | 141Pr | Standard BioTools |
| IL4* | MP4-25D2 | 3142002B | 142Nd | Standard BioTools |
| CD38 | HIT2 | 303535 | 143Nd | BioLegend |
| CD195/CCR5 | NP6G4 | 3144007A | 144Nd | Standard BioTools |
| CD30 | BerH8 | 555827 | 145Nd | BD |
| CD8α | RPAT8 | 3146001B | 146Nd | Standard BioTools |
| CXCR4 | 12G5 | 306523 | 147Sm | BioLegend |
| CD278/ICOS | C398.4A | 3148019B | 148Nd | Standard BioTools |
| CD25 | 2A3 | 3149010B | 149Sm | Standard BioTools |
| MIP1β* | D211351 | 3150004B | 150Nd | Standard BioTools |
| CD107a/LAMP1* | H4A3 | 3151002B | 151Eu | Standard BioTools |
| TNFα* | Mab11 | 3152002B | 152Sm | Standard BioTools |
| CD62L/L-selectin | DREG56 | 3153004B | 153Eu | Standard BioTools |
| CD95 | 50825 | MAB326100 | 154Sm | R&D |
| CD279/PD1 | EH12.2H7 | 3155009B | 155Gd | Standard BioTools |
| CD29 | TS2/16 | 3156007B | 156Gd | Standard BioTools |
| CTLA4* | 14D3 | 5012919 | 157Gd | eBioscience |
| CD134/OX40 | ACT35 | 3158012B | 158Gd | Standard BioTools |
| CD197/CCR7 | G043H7 | 3159003A | 159Tb | Standard BioTools |
| CD28 | CD28.2 | 3160003B | 160Gd | Standard BioTools |
| Ki-67* | B56 | 3161007B | 161Dy | Standard BioTools |
| CD69 | FN50 | 3162001B | 162Dy | Standard BioTools |
| IL6* | MQ2-13A5 | 501115 | 163Dy | BioLegend |
| CD45RO | UCHL1 | 3164007B | 164Dy | Standard BioTools |
| CD127/IL7Ra | A019D5 | 3165008B | 165Ho | Standard BioTools |
| IL2* | MQ117H12 | 3166002B | 166Er | Standard BioTools |
| CD27 | L128 | 3167006B | 167Er | Standard BioTools |
| IFNγ* | B27 | 3168005B | 168Er | Standard BioTools |
| CD45RA | HI100 | 3169008B | 169Tm | Standard BioTools |
| CD3 | UCHT1 | 3170001B | 170Er | Standard BioTools |
| CD185/CXCR5 | RF8B2 | 3171014B | 171Yb | Standard BioTools |
| CD57 | HCD57 | 3172009B | 172Yb | Standard BioTools |
| Granzyme B* | GB11 | 3173006B | 173Yb | Standard BioTools |
| CD4 | SK3 | 3174004B | 174Yb | Standard BioTools |
| Perforin* | BD48 | 3175004B | 175Lu | Standard BioTools |
| Foxp3* | 206D | 320102 | 176Yb | BioLegend |
| TIGIT | MBSA43 | 3209013B | 209Bi | Standard BioTools |
| IL17* | BL168 | 512331 | 89Y | BioLegend |
| HLA-DR | TU36 | Q22158 | Qdot(112Cd) | Invitrogen |

\*Intracellular staining.

**Table S3:**
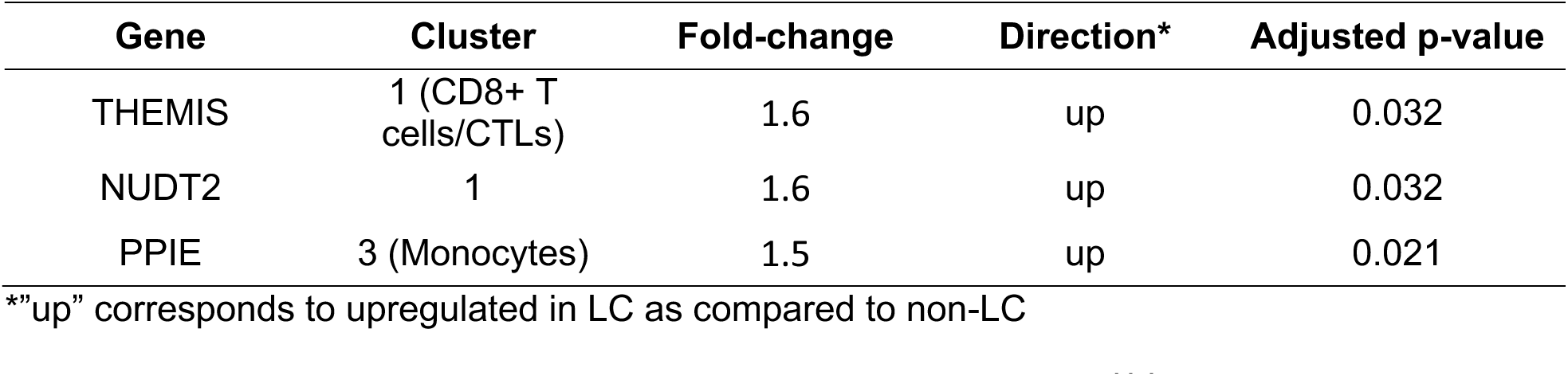
scRNAseq of differentially expressed genes in LC (n=8) vs. non-LC (n=4) (p<0.05)

**Table S4:** scRNAseq of differentially expressed genes in OR7D2^high^ (n=4) vs. non-LC (n=4) (p<0.05)

| Gene | Cluster | Fold-change | Direction* | Adjusted p-value |
| --- | --- | --- | --- | --- |
| HIST1H2AM | 0 (CD4+ T cells) | 2.7 | up | 0.001 |
| HIST2H2AC | 0 | 1.8 | up | 0.034 |
| CR1 | 0 | 3.4 | up | 0.034 |
| FGD5-AS1 | 0 | 1.4 | up | 0.034 |
| SVIL | 0 | 1.6 | up | 0.034 |
| PIP4K2A | 0 | 1.3 | up | 0.036 |
| NORAD | 0 | 1.4 | up | 0.036 |
| NUDT2 | 0 | 1.6 | up | 0.038 |
| IFFO2 | 0 | 1.5 | up | 0.039 |
| DENND1B | 0 | 1.3 | up | 0.039 |
| AC147067.1 | 0 | 1.8 | up | 0.039 |
| RASSF6 | 0 | 3.4 | up | 0.039 |
| ARL15 | 0 | 1.3 | up | 0.039 |
| ARRDC3-AS1 | 0 | 1.7 | up | 0.039 |
| AHNAK | 0 | 1.8 | up | 0.039 |
| HIST1H2AK | 0 | 2.1 | up | 0.044 |
| SPCS3 | 0 | 1.3 | up | 0.047 |
| HIST1H2AM | 1 (CD8+ T cells/CTLs) | 1.6 | up | 0.004 |
| ITGB1 | 5 (CD8+ T cells) | 1.4 | up | 0.026 |
| AHNAK | 5 | 8.0 | up | 0.026 |
| S100A6 | 5 | 2.9 | up | 0.037 |
| HIST1H1E | 5 | 1.8 | up | 0.037 |
| HIST1H2AM | 5 | 1.7 | up | 0.037 |
| H1FX | 5 | 1.5 | up | 0.037 |
| KLF6 | 5 | 1.8 | up | 0.037 |
| MYO1F | 5 | 3.2 | up | 0.037 |
| TTC39C | 5 | 1.6 | up | 0.040 |
| IGKV2-24 | 8 (B cells) | 1.7 | up | 0.011 |
| APOO | 0 | 2.4 | down | 0.034 |
| MTFP1 | 0 | 1.7 | down | 0.036 |
| AC090360.1 | 0 | 1.6 | down | 0.039 |
| APOO | 5 | 2.1 | down | 0.026 |
| PECAM1 | 5 | 1.6 | down | 0.037 |
| APOO | 7 (CD4+ T cells) | 143.1 | down | 0.027 |
| RGPD2 | 8 (B cells) | 5.1 | down | 0.002 |

\*\*"up" corresponds to upregulated in OR7D2<sup>high</sup> LC as compared to non-LC, "down" corresponds to down-regulated in OR7D2<sup>high</sup> LC as compared to non-LC

**Table S5:**
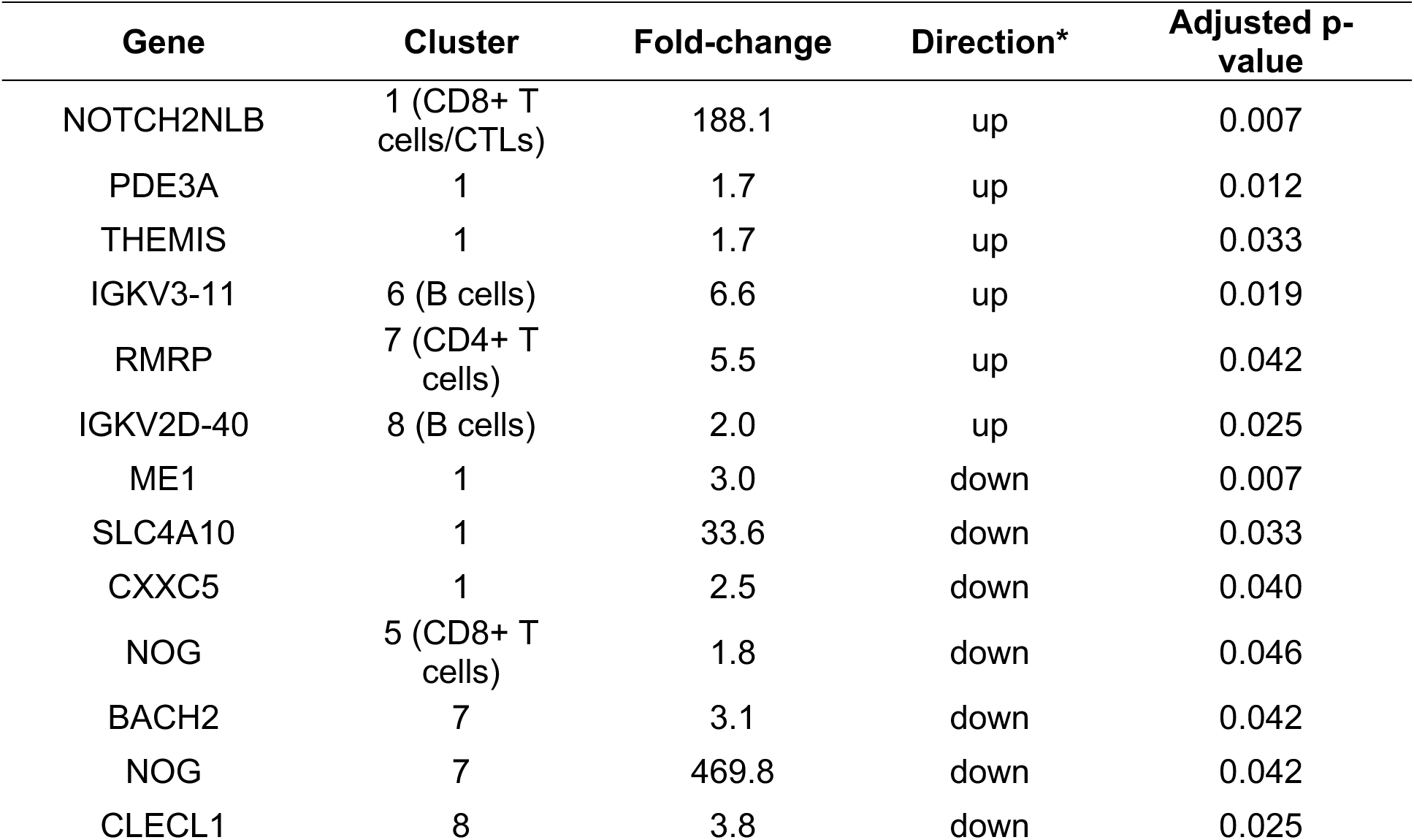
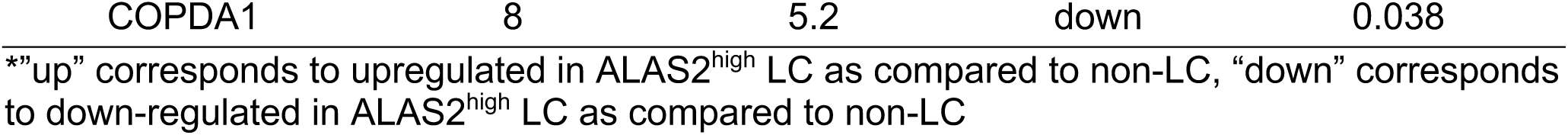
scRNAseq of differentially expressed genes in ALAS2^high^ (n=4) vs. non-LC (n=4) (p<0.05)

**Table S6:**
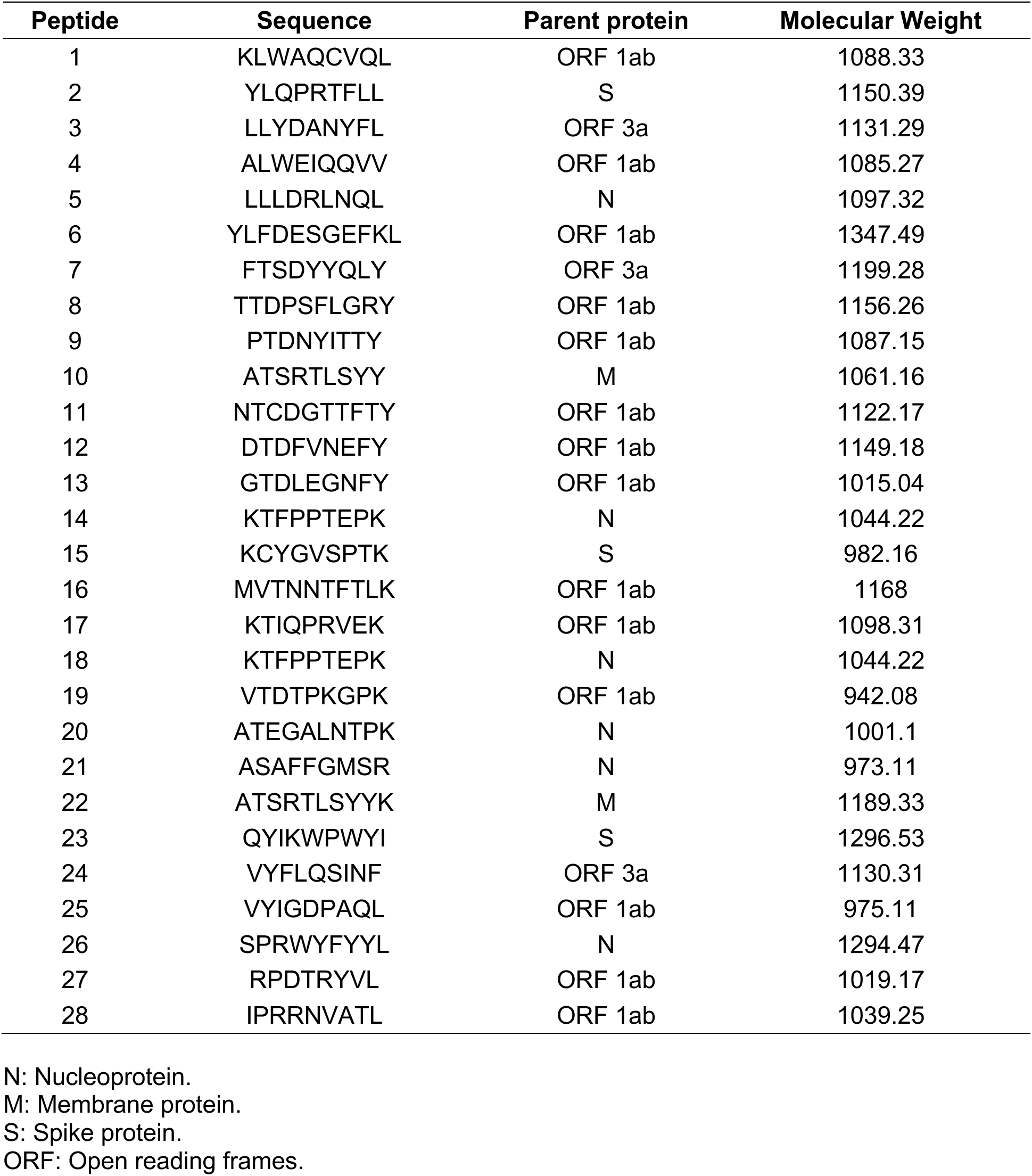
T-scan peptides used in study.

**Table S7:** Flow cytometry antibodies used in study.

| **Antibody** | **Clone** | **Catalog** | **Fluorochrome** | **Manufacturer** |
| --- | --- | --- | --- | --- |
| CD3 | UCHT1 | 11-0038-41 | FITC | Invitrogen |
| CD4 | SK3 | 566356 | BV750 | BD Biosciences |
| CXCR4 | 12G5 | 17-9999-41 | APC | Invitrogen |
| CXCR5 | J252D4 | 356919 | BV421 | BioLegend |
| CCR6 | 11A9 | 562724 | BV605 | BD Biosciences |
| CD45RO | UCHL1 | 304227 | APC Cy7 | BioLegend |
| live/dead | NA | 423107 | zombie UV | BioLegend |
| CD3 | UVHT1 | 612941 | BUV496 | BD Biosciences |
| CD8a | RPA-T8 | 301005 | FITC | BioLegend |
| PD-1 | EH12-2H7 | 329965 | BV750 | BioLegend |
| CTLA4* | L3D10 | 349913 | PE-Cy7 | BioLegend |
| IFNγ* | B27 | 566395 | BB700 | BD Biosciences |
| TNFα* | Mab11 | 502937 | BV650 | BioLegend |
| MIP-1β* | D21-1351 | 562900 | BV421 | BD Biosciences |
| live/dead | NA | 423105 | zombie NIR | BioLegend |
| CD28 | L293 | 340975 | NA | BD Biosciences |
| CD49d | L25 | 340976 | NA | BD Biosciences |

\*Intracellular staining.

