## Supplementary Figures for "Long COVID manifests with T cell dysregulation, inflammation, and an uncoordinated adaptive immune response to SARS-CoV-2"

### Figure S1

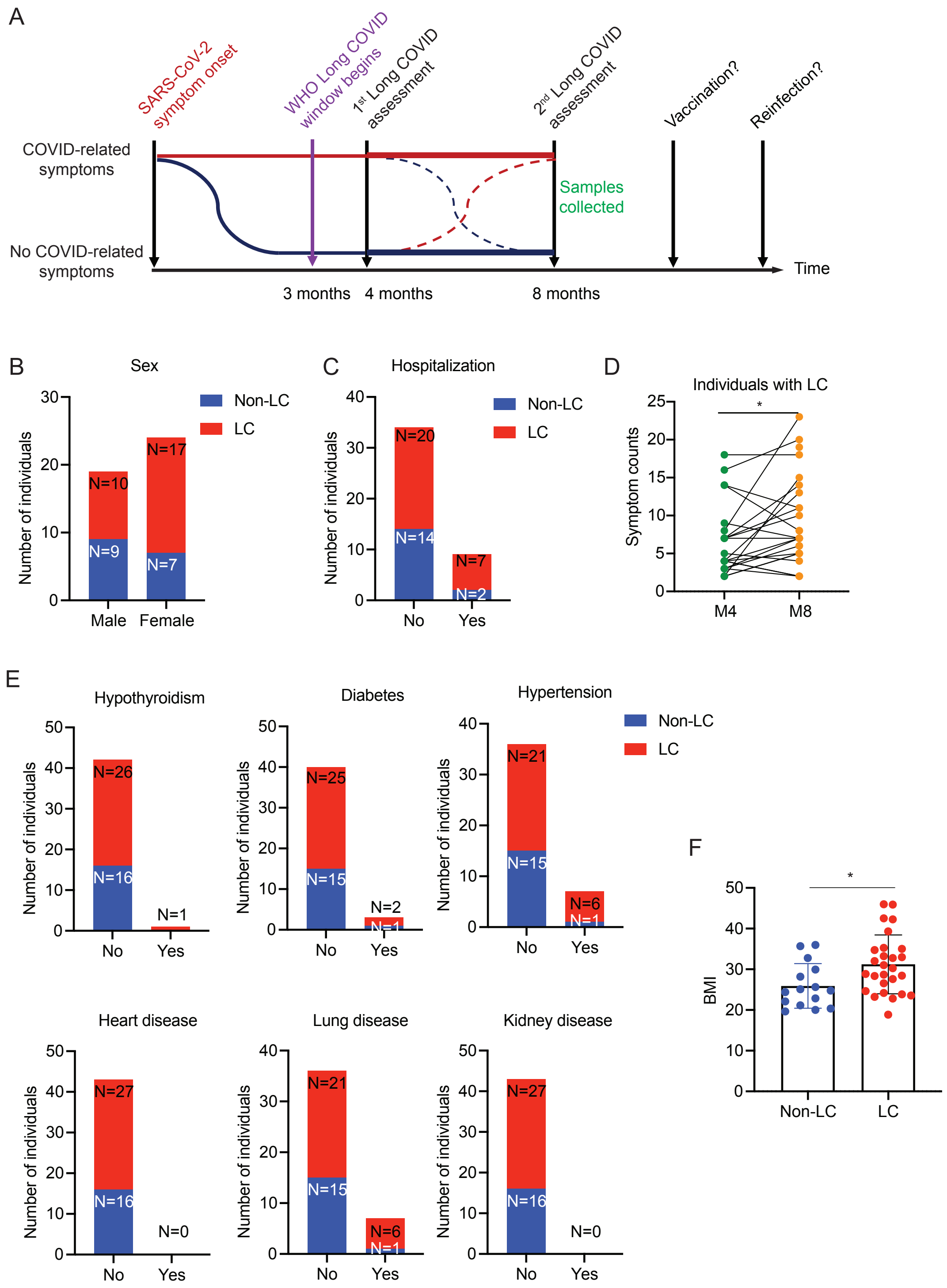
A
SARS-CoV-2 symptom onset
WHO Long COVID window begins
1st Long COVID assessment
2nd Long COVID assessment
Vaccination?
Reinfection?
COVID-related symptoms
No COVID-related symptoms
Samples collected
Time
3 months
4 months
8 months
B
Sex
Non-LC
LC
Number of individuals
N=10
N=17
N=9
N=7
Male
Female
C
Hospitalization
N=20
N=14
N=7
N=2
No
Yes
D
Individuals with LC
Symptom counts
M4
M8
E
Hypothyroidism
N=26
N=16
N=1
Diabetes
N=25
N=15
N=2
N=1
Hypertension
N=21
N=15
N=6
N=1
Heart disease
N=27
N=16
N=0
Lung disease
N=21
N=15
N=6
N=1
Kidney disease
N=27
N=16
N=0
F
BMI
Non-LC
LC

Figure S2

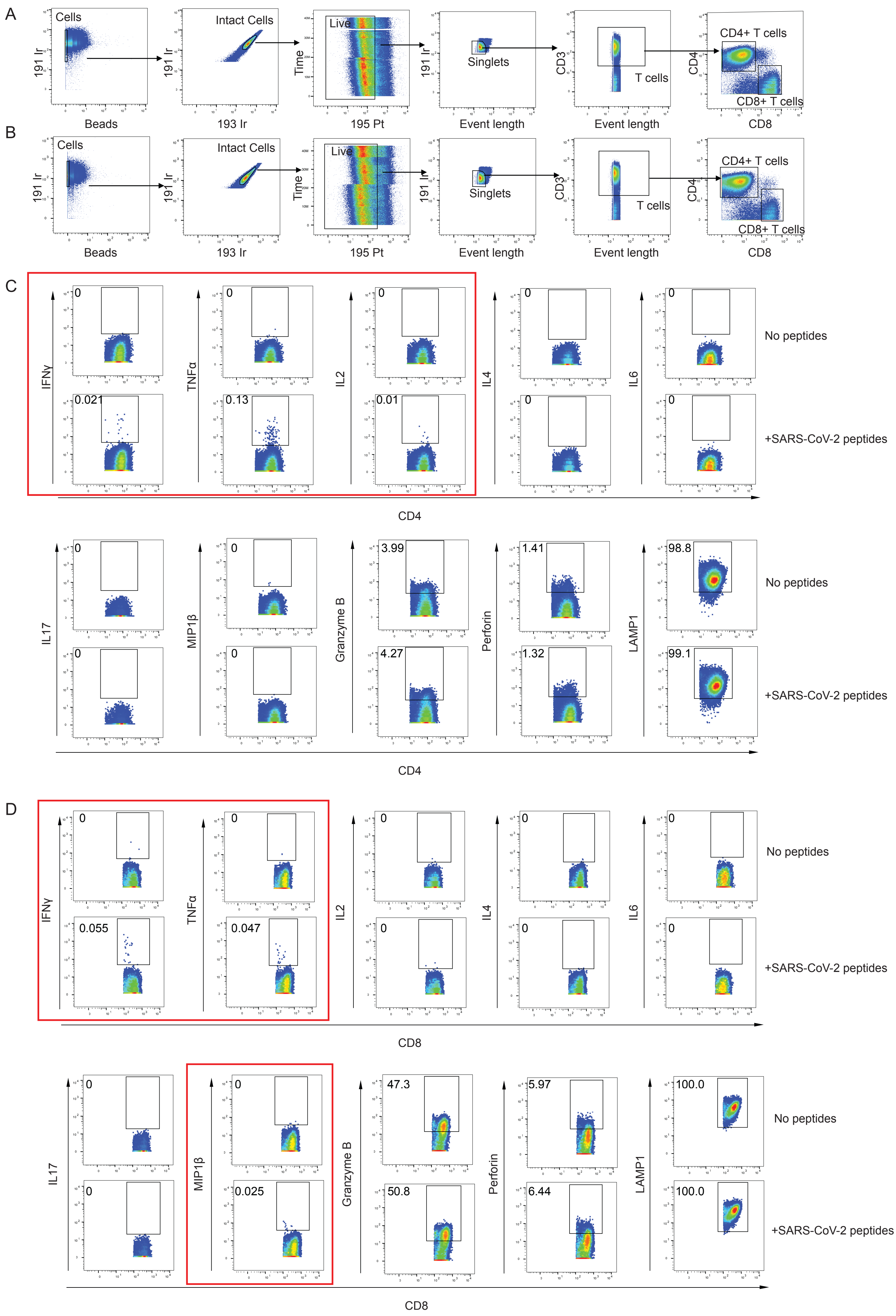
A
Cells
191 Ir
Beads
Intact Cells
193 Ir
Live
Time
195 Pt
Singlets
Event length
CD3
T cells
CD4+ T cells
CD4
CD8+ T cells
CD8
B
Cells
191 Ir
Beads
Intact Cells
193 Ir
Live
Time
195 Pt
Singlets
Event length
CD3
T cells
CD4+ T cells
CD4
CD8+ T cells
CD8
C
IFNγ
0
0.021
TNFα
0
0.13
IL2
0
0.01
IL4
0
0
IL6
0
0
No peptides
+SARS-CoV-2 peptides
CD4
IL17
0
0
MIP1β
0
0
Granzyme B
3.99
4.27
Perforin
1.41
1.32
LAMP1
98.8
99.1
No peptides
+SARS-CoV-2 peptides
CD4
D
IFNγ
0
0.055
TNFα
0
0.047
IL2
0
0
IL4
0
0
IL6
0
0
No peptides
+SARS-CoV-2 peptides
CD8
IL17
0
0
MIP1β
0
0.025
Granzyme B
47.3
50.8
Perforin
5.97
6.44
LAMP1
100.0
100.0
No peptides
+SARS-CoV-2 peptides
CD8

### Figure S3

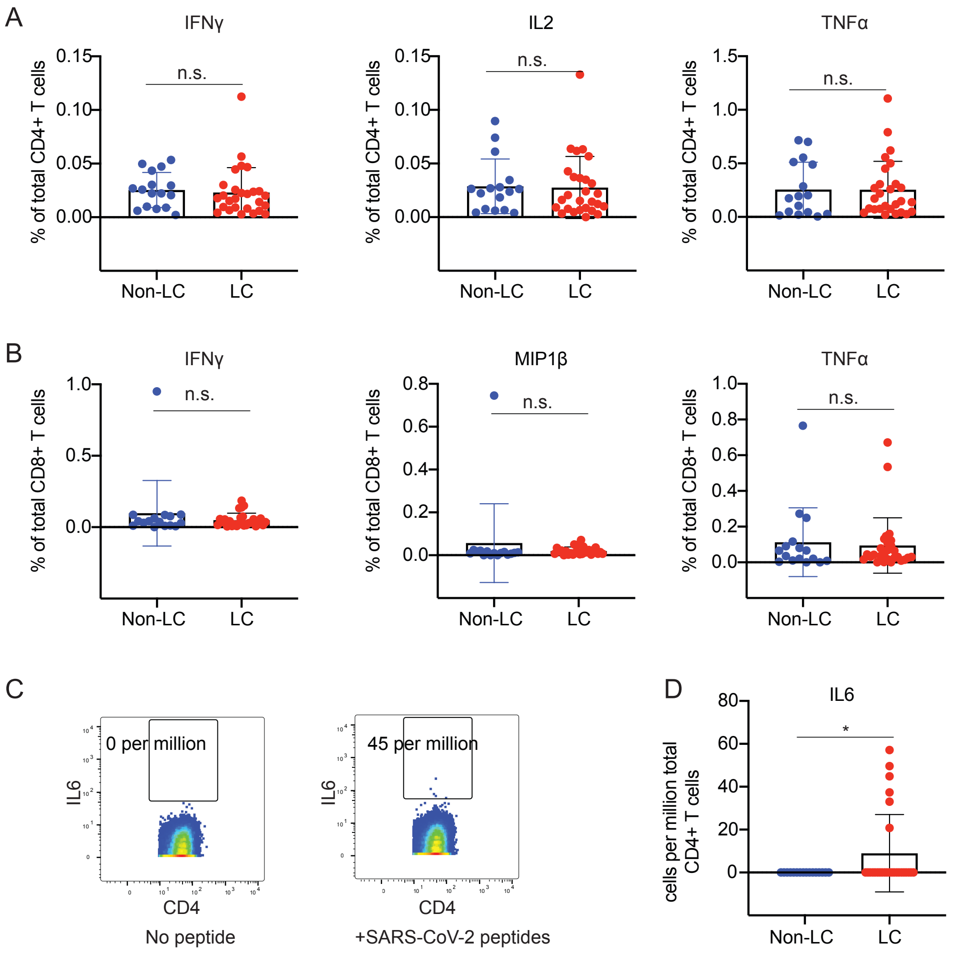
A
IFNγ
IL2
TNFα
% of total CD4+ T cells
n.s.
Non-LC
LC
B
MIP1β
% of total CD8+ T cells
C
0 per million
45 per million
IL6
CD4
No peptide
+SARS-CoV-2 peptides
D
cells per million total CD4+ T cells
*

Figure S4

A **Total CD4+ T cells**

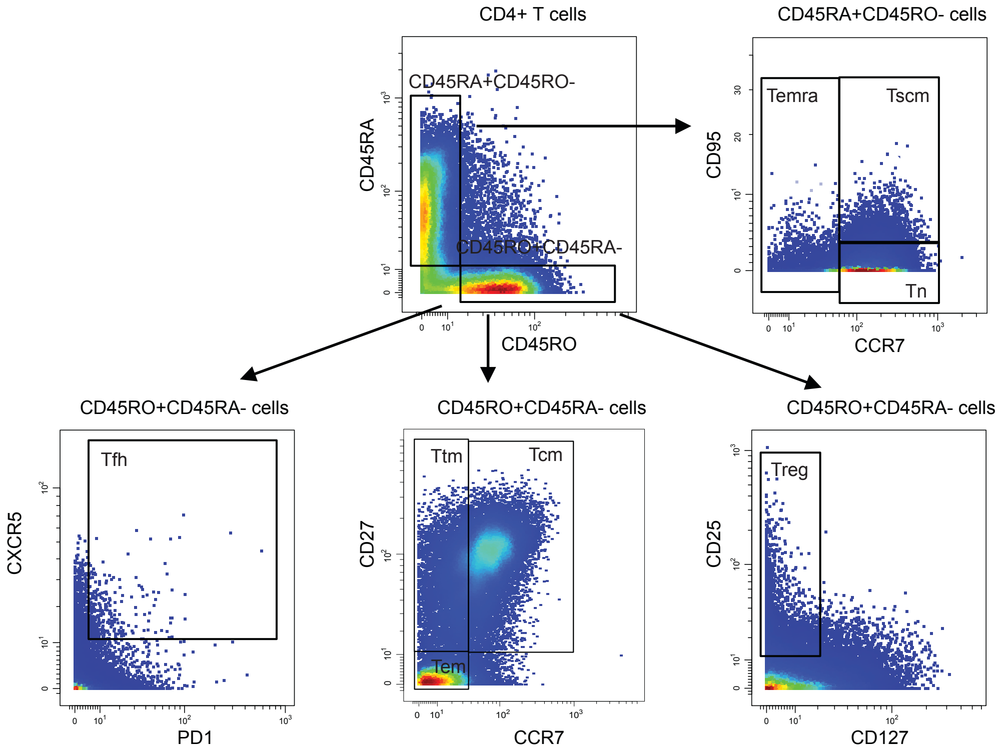


B **Total CD8+ T cells**

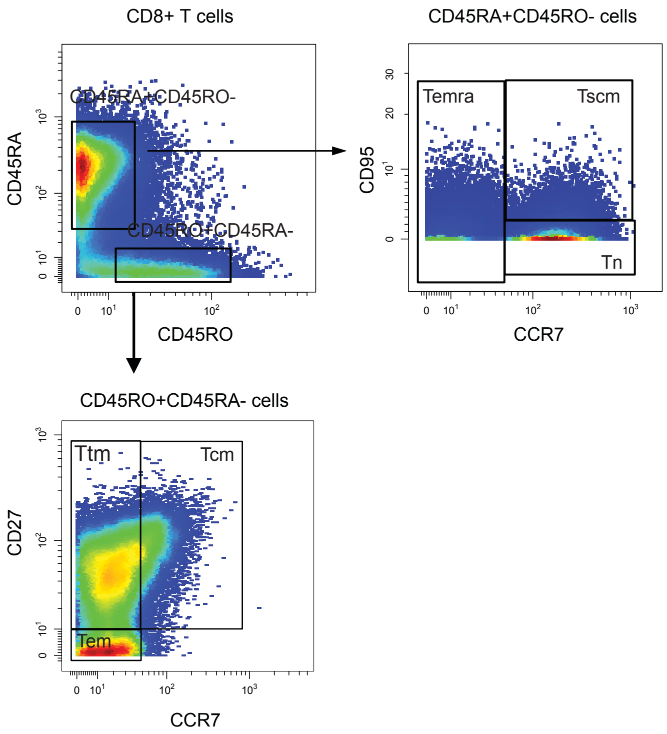

Figure S5

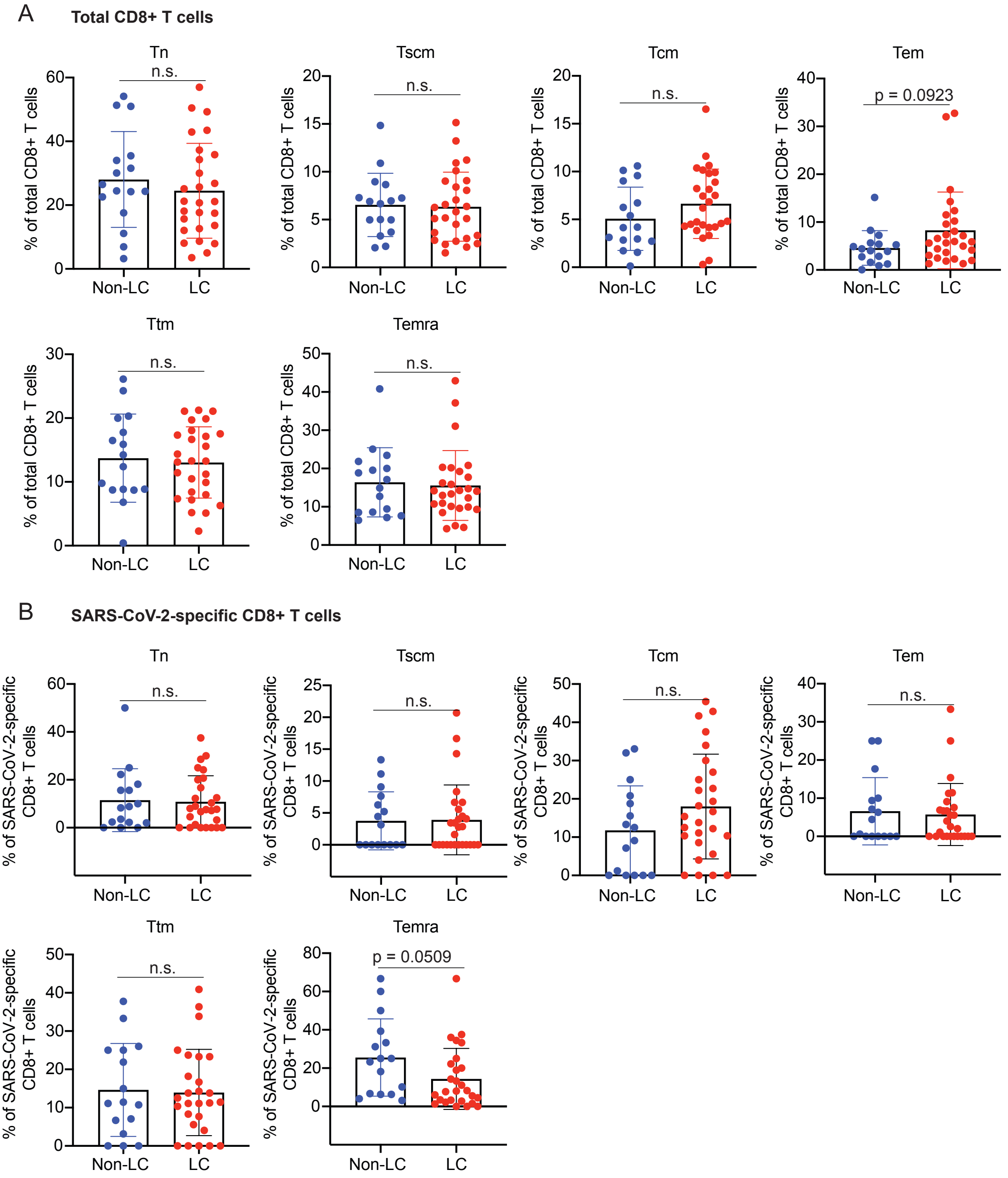
A
Total CD8+ T cells
Tn
Tscm
Tcm
Tem
Ttm
Temra
% of total CD8+ T cells
n.s.
p = 0.0923
Non-LC
LC
B
SARS-CoV-2-specific CD8+ T cells
% of SARS-CoV-2-specific CD8+ T cells
p = 0.0509

Figure S6

**Total CD4+ T cells**

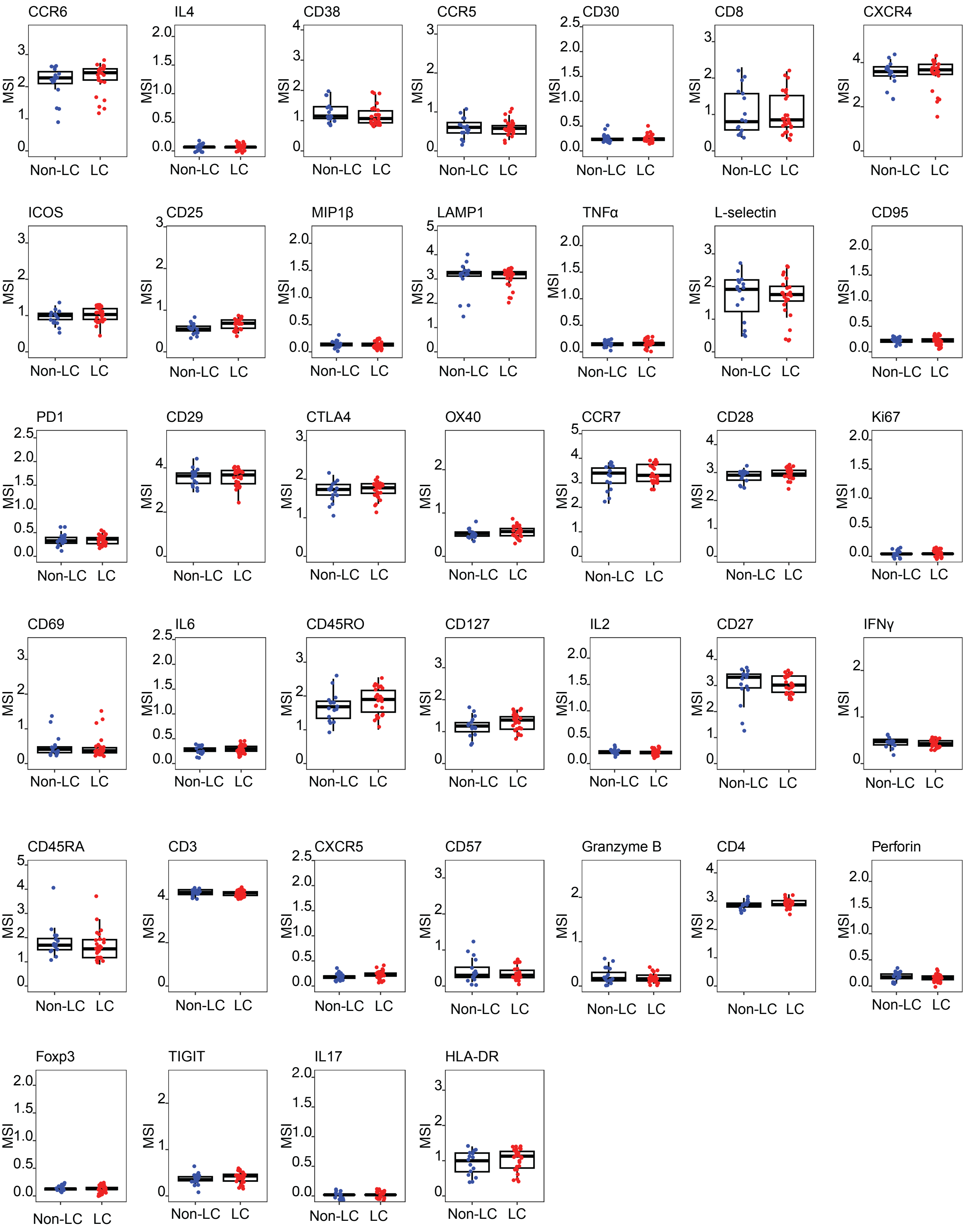

Figure S7

**Total CD8+ T cells**

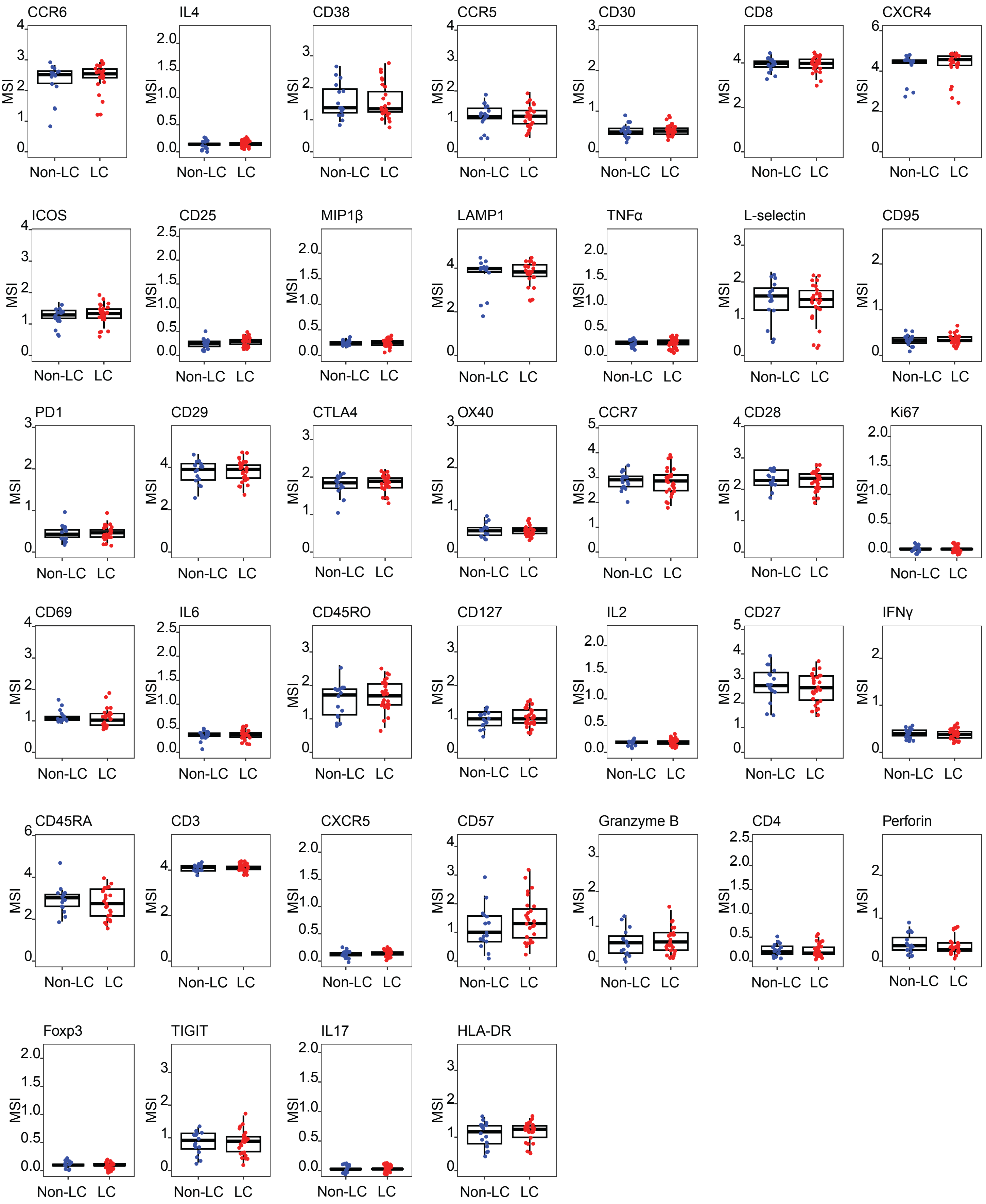

Figure S8

**SARS-CoV-2-specific CD4+ T cells**

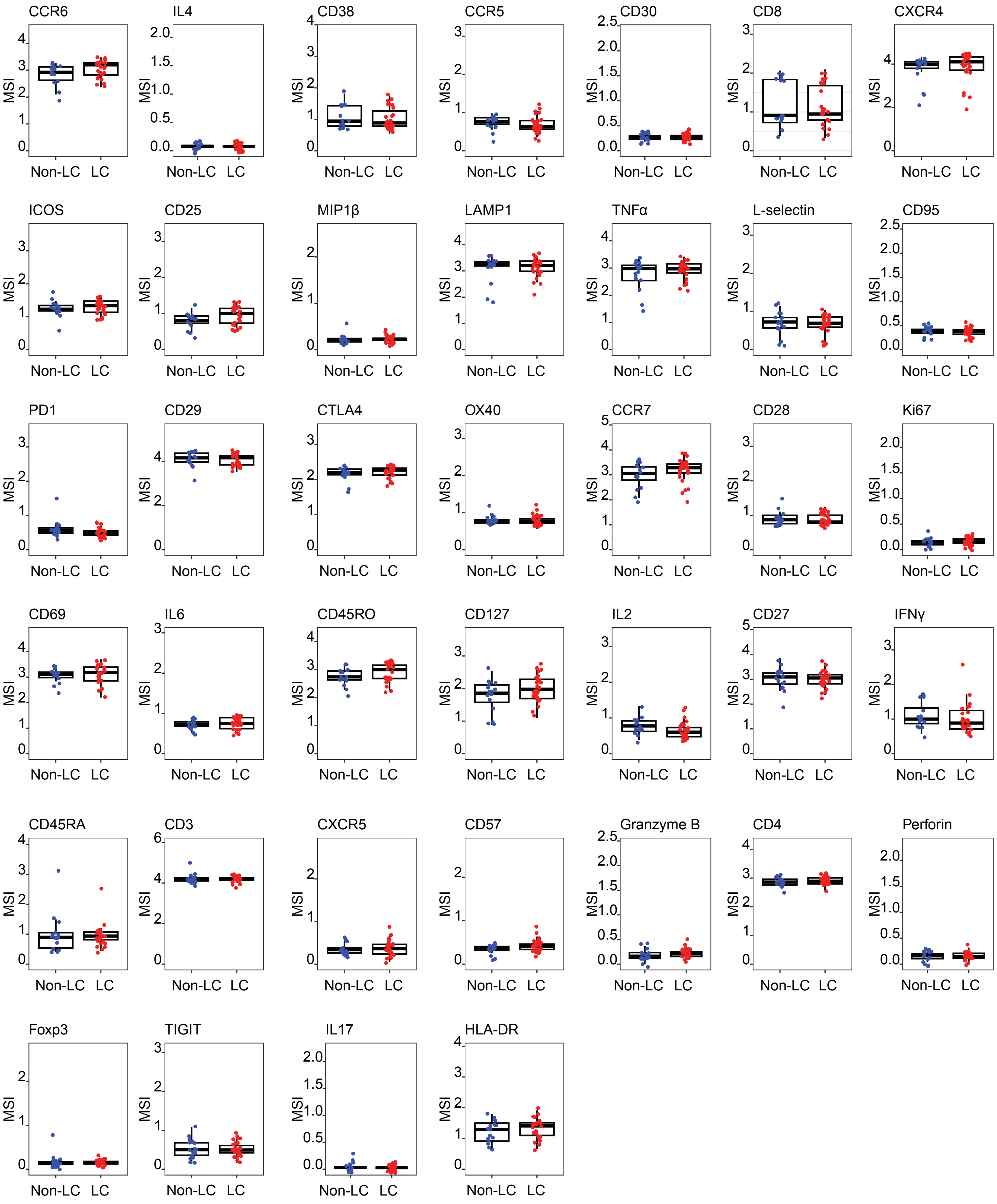

Figure S9

**SARS-CoV-2-specific CD8+ T cells**

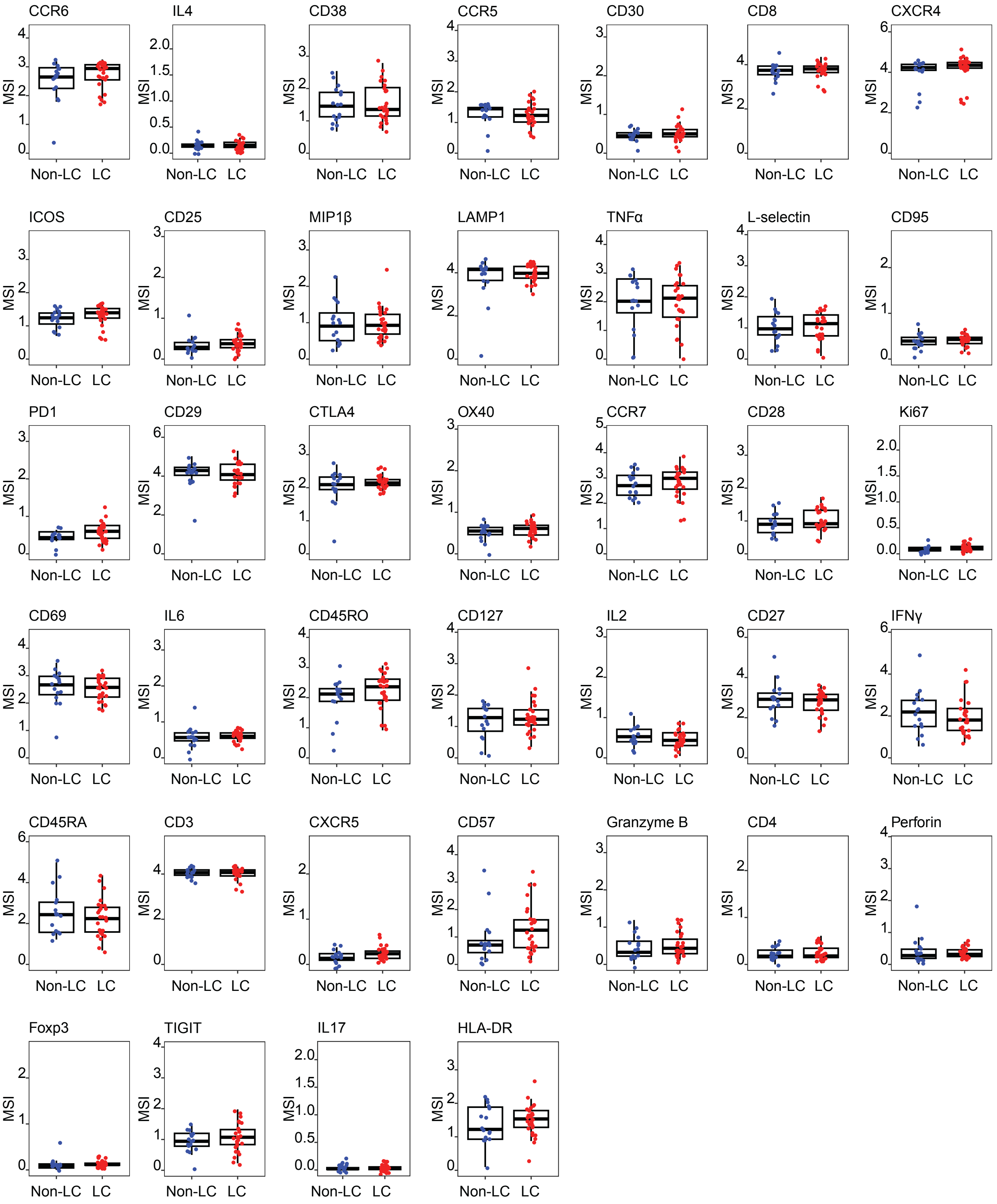

Figure S10

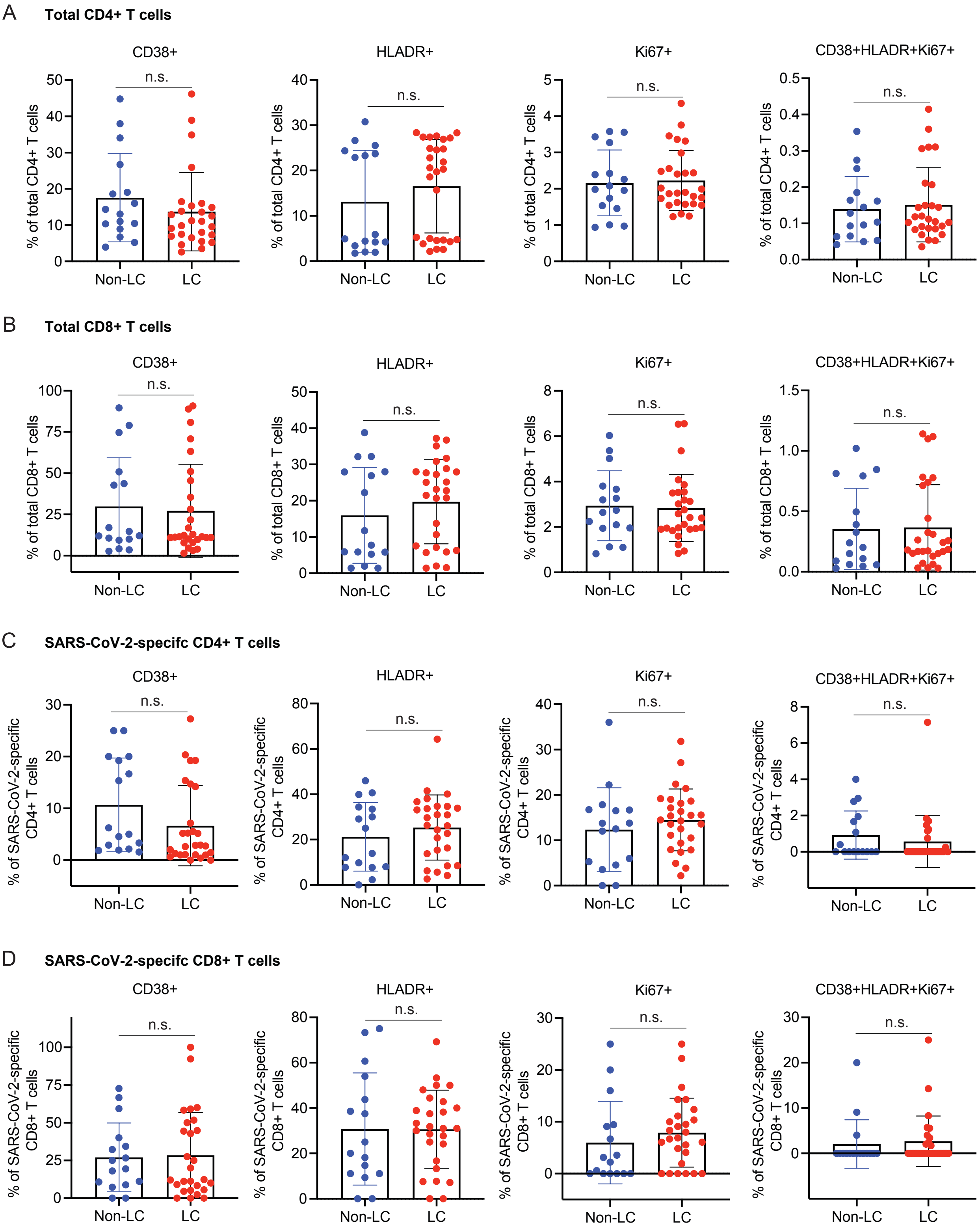
A
Total CD4+ T cells
CD38+
HLADR+
Ki67+
CD38+HLADR+Ki67+
% of total CD4+ T cells
n.s.
Non-LC
LC
B
Total CD8+ T cells
CD38+
HLADR+
Ki67+
CD38+HLADR+Ki67+
% of total CD8+ T cells
n.s.
Non-LC
LC
C
SARS-CoV-2-specifc CD4+ T cells
CD38+
HLADR+
Ki67+
CD38+HLADR+Ki67+
% of SARS-CoV-2-specific CD4+ T cells
n.s.
Non-LC
LC
D
SARS-CoV-2-specifc CD8+ T cells
CD38+
HLADR+
Ki67+
CD38+HLADR+Ki67+
% of SARS-CoV-2-specific CD8+ T cells
n.s.
Non-LC
LC

### Figure S11

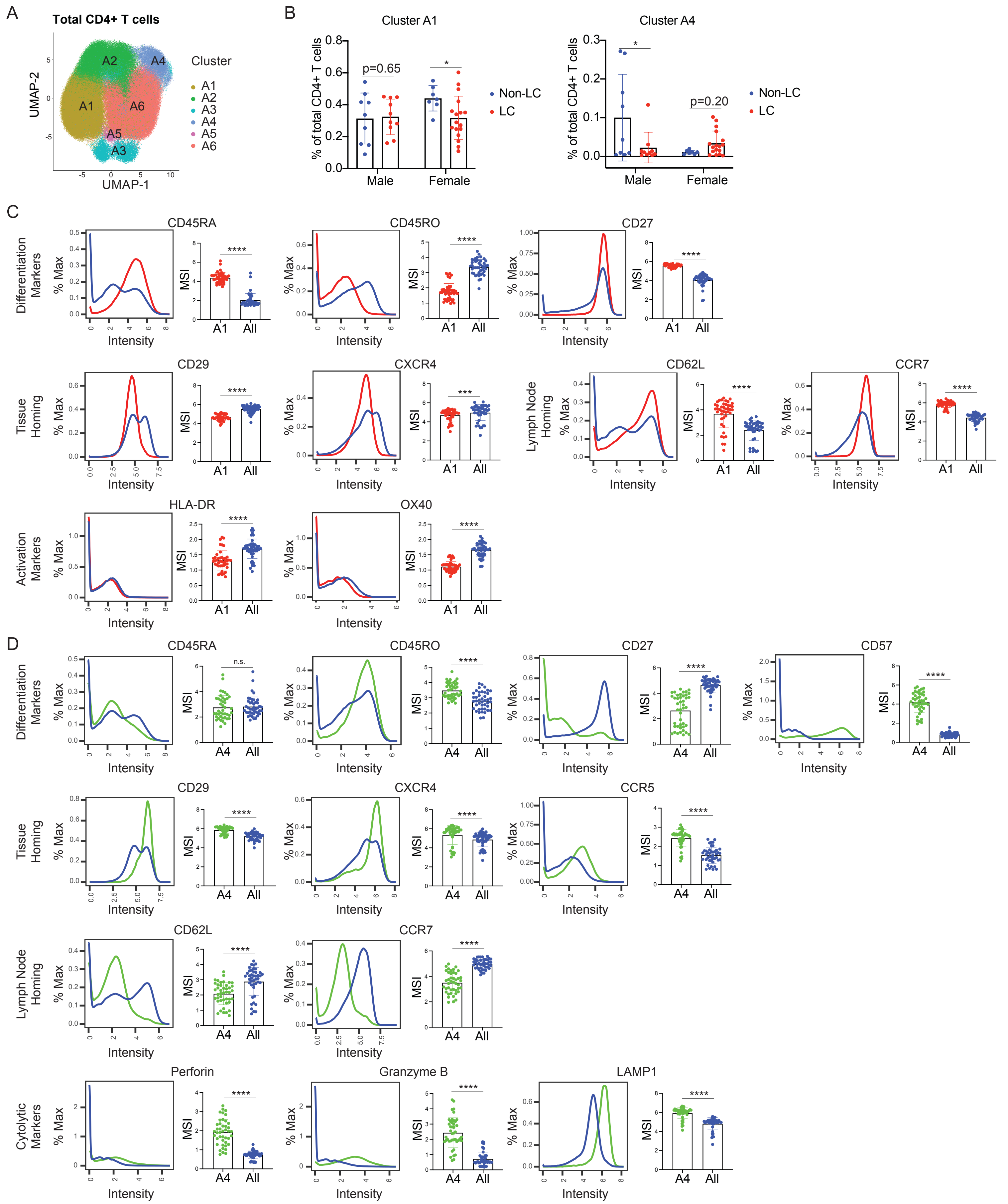
A
Total CD4+ T cells
UMAP-2
UMAP-1
Cluster
A1
A2
A3
A4
A5
A6
B
Cluster A1
Cluster A4
% of total CD4+ T cells
p=0.65
p=0.20
Non-LC
LC
Male
Female
C
Differentiation Markers
Tissue Homing
Lymph Node Homing
Activation Markers
CD45RA
CD45RO
CD27
CD29
CXCR4
CD62L
CCR7
HLA-DR
OX40
% Max
MSI
Intensity
A1
All
D
Differentiation Markers
Tissue Homing
Lymph Node Homing
Cytolytic Markers
CD45RA
CD45RO
CD27
CD57
CD29
CXCR4
CCR5
CD62L
CCR7
Perforin
Granzyme B
LAMP1
n.s.
A4
All

### Figure S12

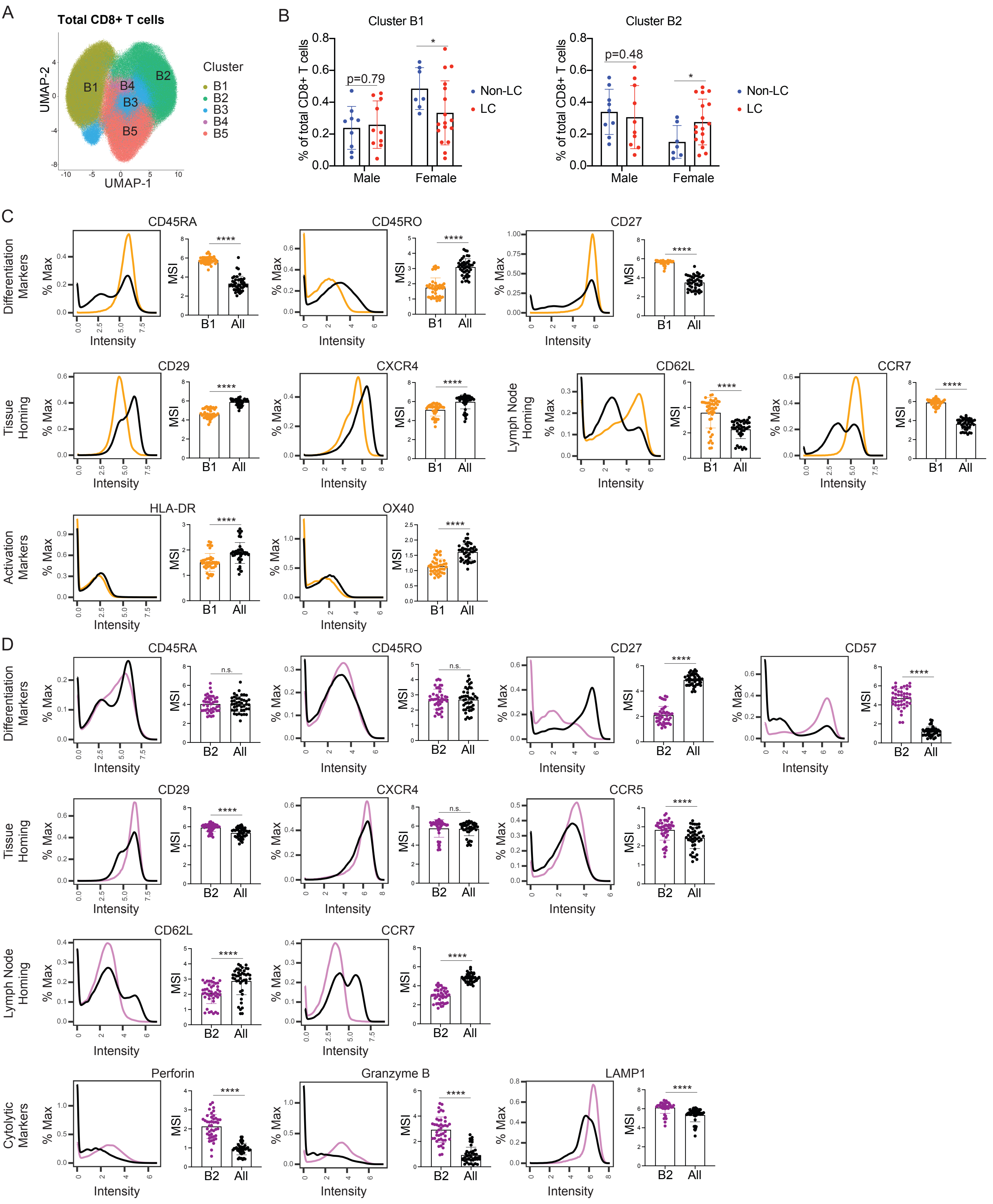
A
Total CD8+ T cells
UMAP-2
UMAP-1
B1
B2
B3
B4
B5
Cluster
B
Cluster B1
Cluster B2
% of total CD8+ T cells
p=0.79
p=0.48
*
Male
Female
Non-LC
LC
C
Differentiation Markers
Tissue Homing
Lymph Node Homing
Activation Markers
CD45RA
CD45RO
CD27
CD29
CXCR4
CD62L
CCR7
HLA-DR
OX40
% Max
MSI
Intensity
****
B1
All
D
Differentiation Markers
Tissue Homing
Lymph Node Homing
Cytolytic Markers
CD45RA
CD45RO
CD27
CD57
CD29
CXCR4
CCR5
CD62L
CCR7
Perforin
Granzyme B
LAMP1
n.s.
****
B2
All

Figure S13

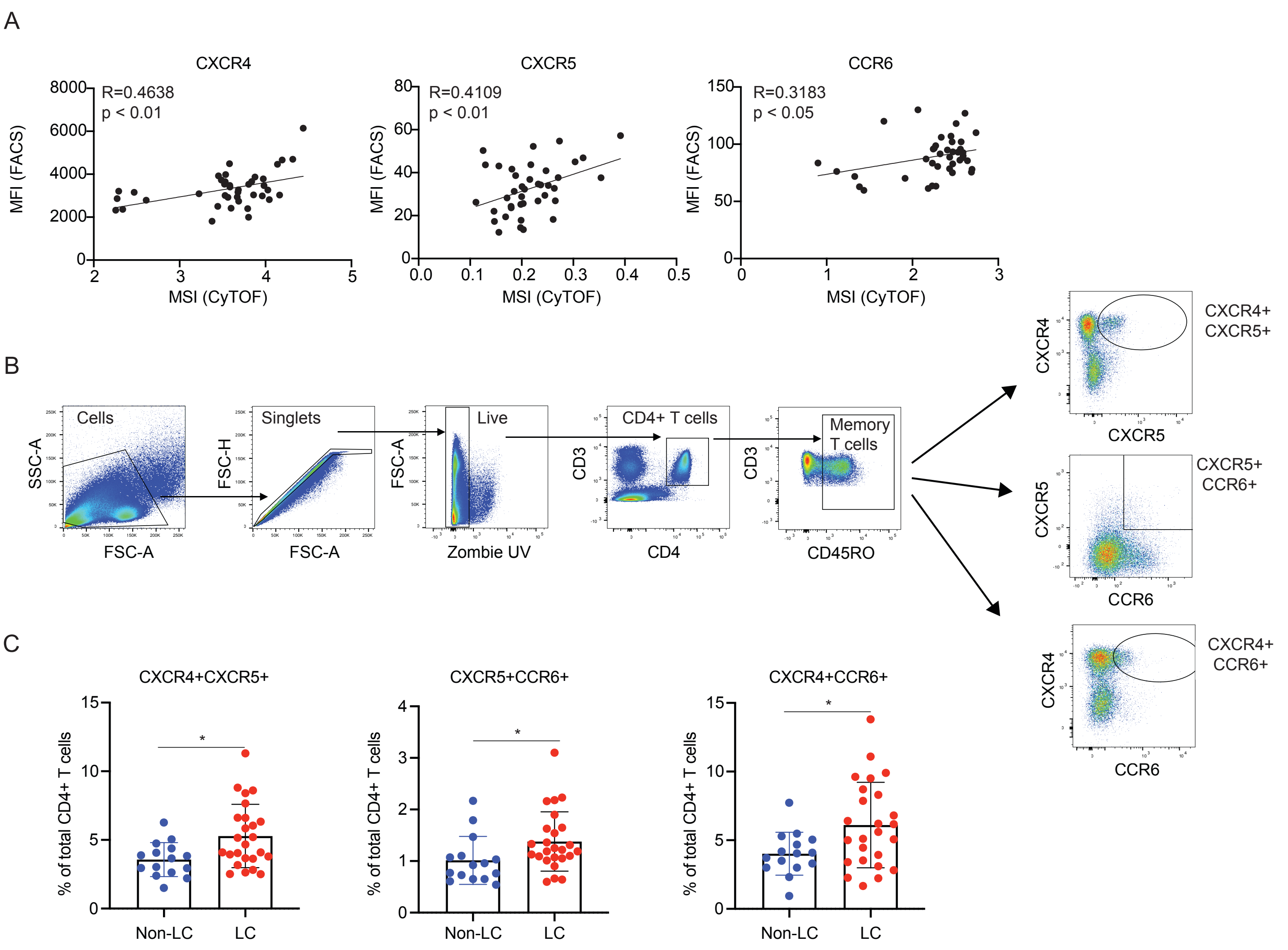
A
CXCR4
R=0.4638
p < 0.01
MFI (FACS)
MSI (CyTOF)
CXCR5
R=0.4109
p < 0.01
CCR6
R=0.3183
p < 0.05
B
Cells
SSC-A
FSC-A
Singlets
FSC-H
Live
Zombie UV
CD4+ T cells
CD3
CD4
Memory T cells
CD45RO
CXCR4+ CXCR5+
CXCR4
CXCR5
CXCR5+ CCR6+
CCR6
CXCR4+ CCR6+
C
CXCR4+CXCR5+
CXCR5+CCR6+
CXCR4+CCR6+
% of total CD4+ T cells
Non-LC
LC
*

### Figure S14

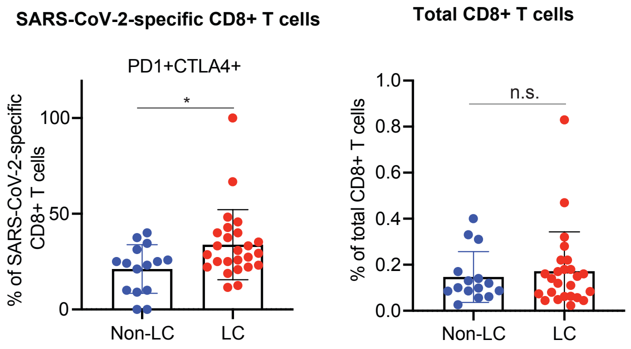
SARS-CoV-2-specific CD8+ T cells
PD1+CTLA4+
% of SARS-CoV-2-specific CD8+ T cells
*
100
50
0
Non-LC
LC
Total CD8+ T cells
% of total CD8+ T cells
n.s.
1.0
0.8
0.6
0.4
0.2
0.0
Non-LC
LC

Figure S15

A

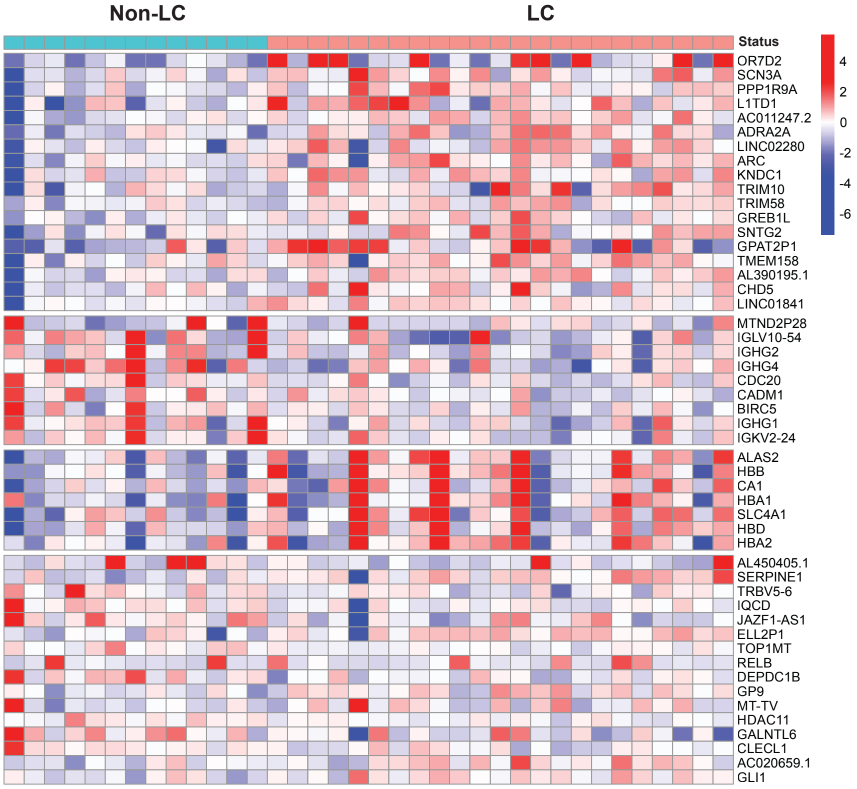


B

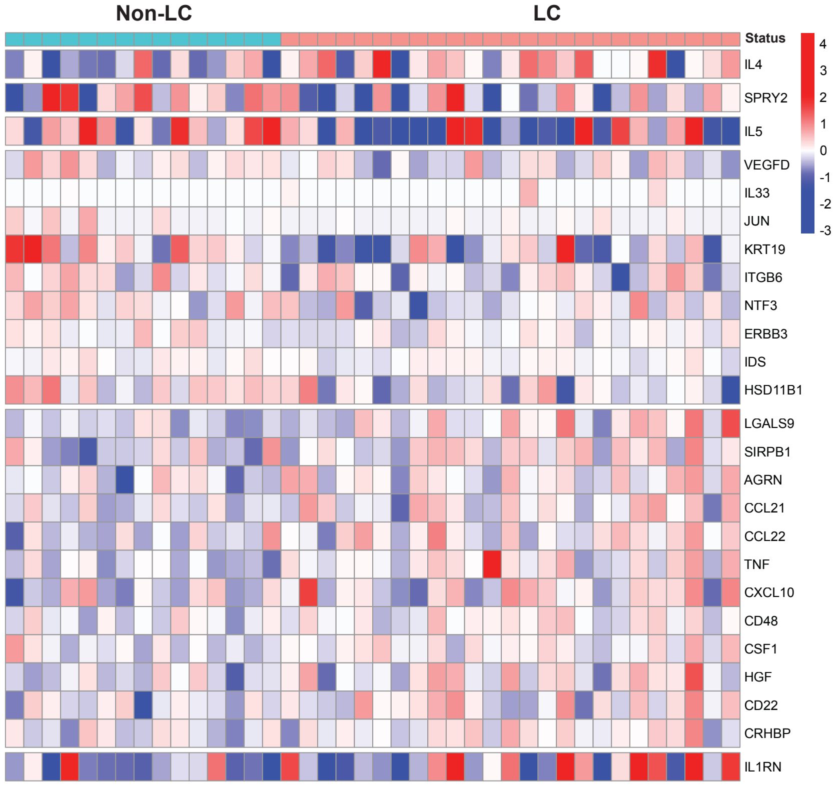

### Figure S16

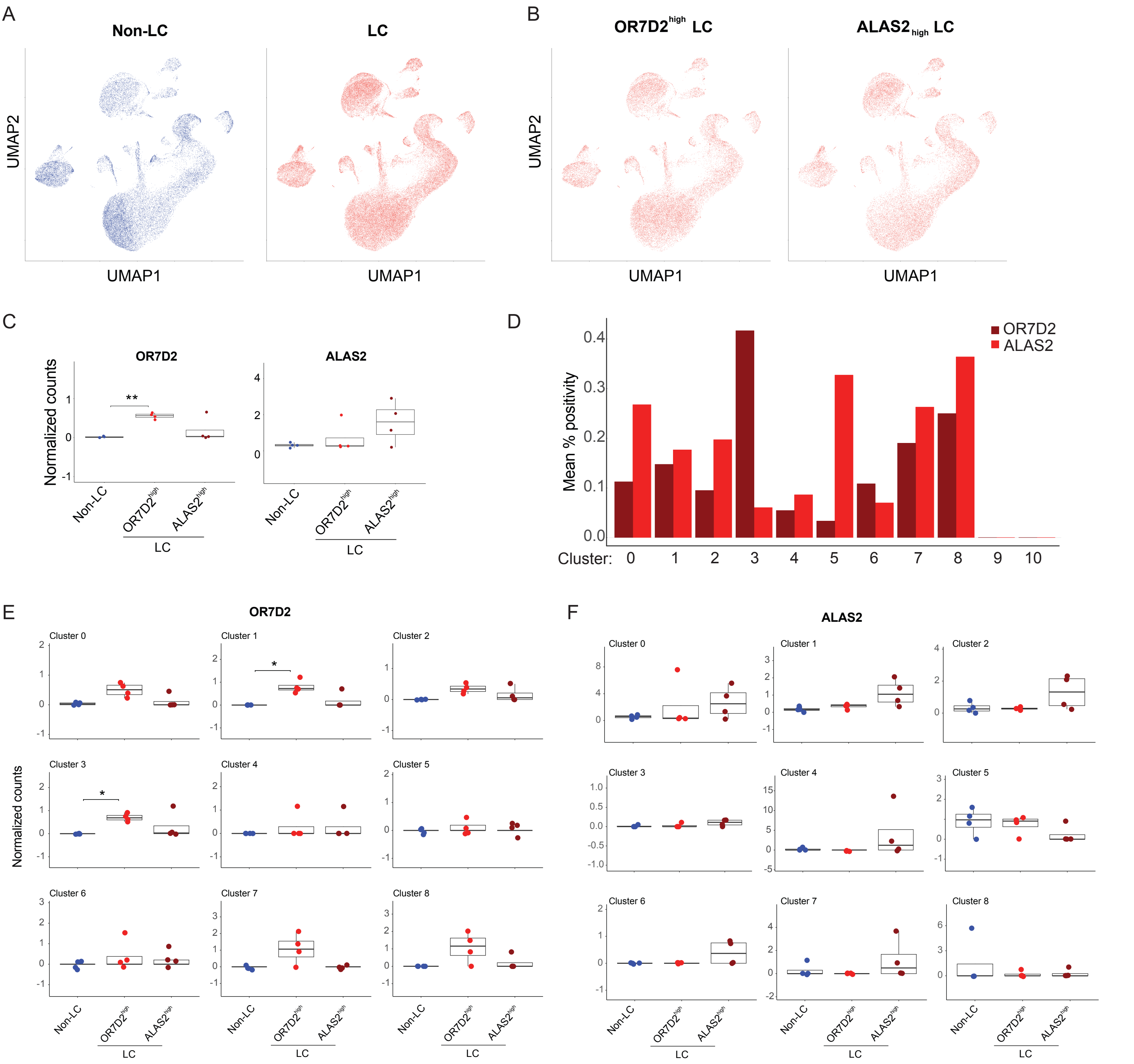
A
Non-LC
LC
UMAP2
UMAP1
B
OR7D2high LC
ALAS2high LC
C
OR7D2
ALAS2
Normalized counts
Non-LC
OR7D2high
ALAS2high
LC
D
Mean % positivity
OR7D2
ALAS2
Cluster: 0 1 2 3 4 5 6 7 8 9 10
E
OR7D2
Cluster 0
Cluster 1
Cluster 2
Cluster 3
Cluster 4
Cluster 5
Cluster 6
Cluster 7
Cluster 8
F
ALAS2

Figure S17

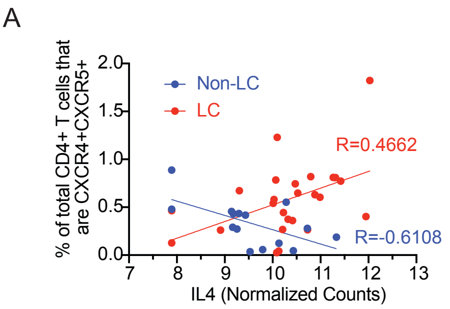
A
% of total CD4+ T cells that are CXCR4+CXCR5+
2.0
1.5
1.0
0.5
0.0
7
8
9
10
11
12
13
IL4 (Normalized Counts)
Non-LC
LC
R=0.4662
R=-0.6108

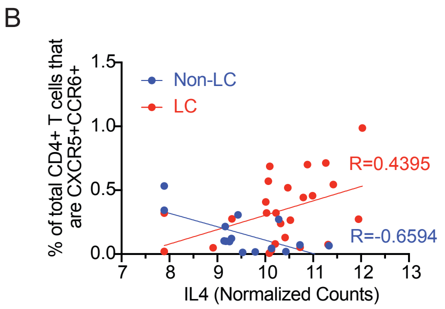
B
% of total CD4+ T cells that are CXCR5+CCR6+
1.5
1.0
0.5
0.0
7
8
9
10
11
12
13
IL4 (Normalized Counts)
Non-LC
LC
R=0.4395
R=-0.6594

### Figure S18

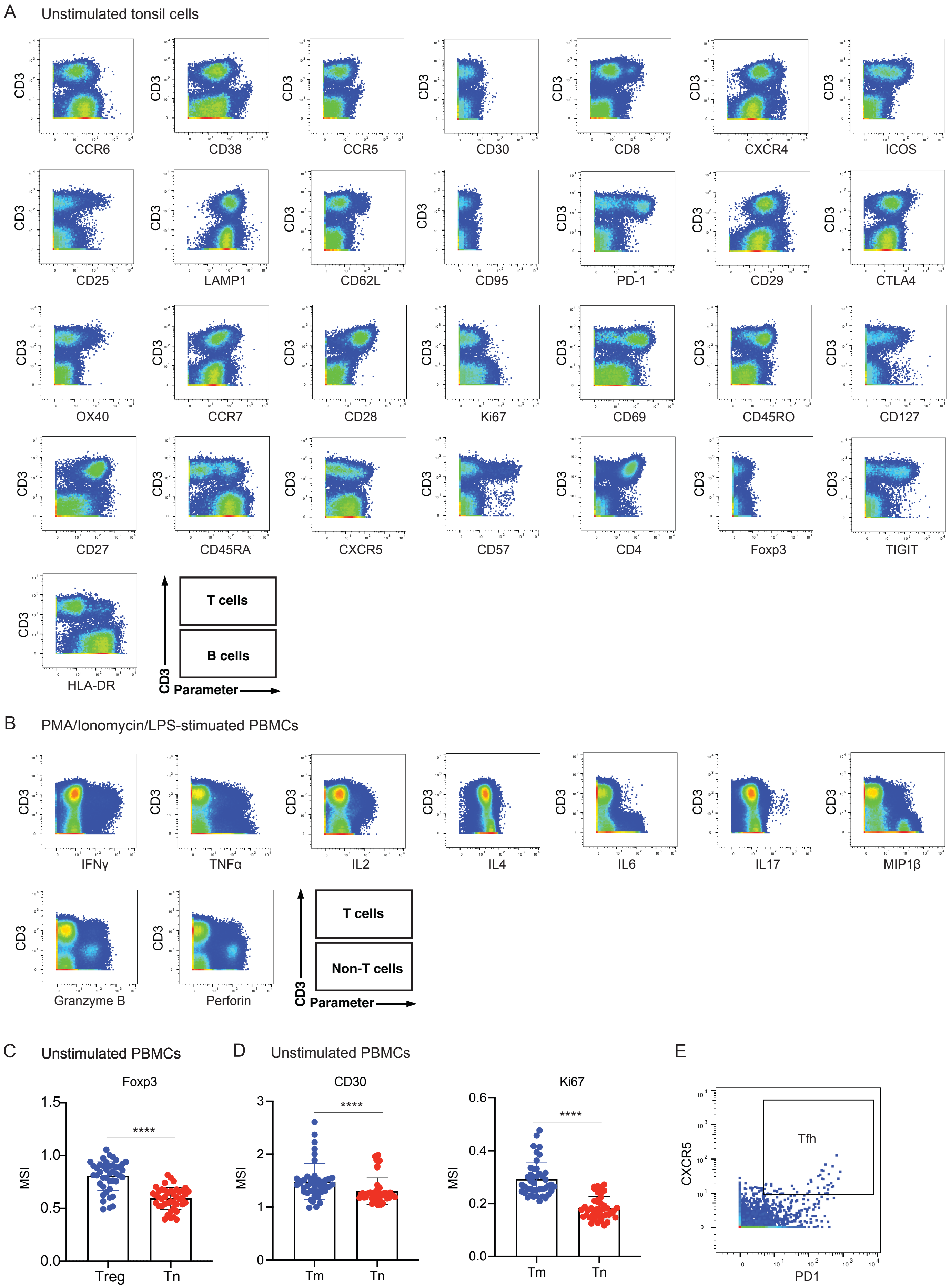

A
Unstimulated tonsil cells
CD3
CCR6
CD38
CCR5
CD30
CD8
CXCR4
ICOS
CD25
LAMP1
CD62L
CD95
PD-1
CD29
CTLA4
OX40
CCR7
CD28
Ki67
CD69
CD45RO
CD127
CD27
CD45RA
CXCR5
CD57
CD4
Foxp3
TIGIT
HLA-DR
T cells
B cells
Parameter
B
PMA/Ionomycin/LPS-stimulated PBMCs
IFNγ
TNFα
IL2
IL4
IL6
IL17
MIP1β
Granzyme B
Perforin
T cells
Non-T cells
Parameter
C
Unstimulated PBMCs
Foxp3
MSI
****
Treg
Tn
D
Unstimulated PBMCs
CD30
Ki67
Tm
Tn
E
Tfh
CXCR5
PD1
